# Zero-shot design of a *de novo* metalloenzyme

**DOI:** 10.64898/2026.04.23.720277

**Authors:** Gina El Nesr, Simon L. Dürr, Irimpan I. Mathews, Qi Wen, Kewei Zhao, Ritimukta Sarangi, Ursula Rothlisberger, Fanny Sunden, Po-Ssu Huang

**Affiliations:** Biophysics Program, Stanford University, Stanford, CA; Institute of Chemical Sciences and Engineering, École Polytechnique Fédérale de Lausanne (EPFL), Lausanne, Switzerland; Institute of Life Sciences, HES-SO Valais-Wallis, Sion, Switzerland; Department of Bioengineering, Stanford University, Stanford, CA; Stanford Synchrotron Radiation Lightsource SLAC National Laboratory, Menlo Park, CA; Department of Biochemistry, Stanford University, Stanford, CA

## Abstract

The *de novo* design of enzymes remains a central challenge, requiring consideration of catalytic mechanism and optimization across biochemical and biophysical criteria. Here we present dEVA (design by EVolutionary Algorithm), a multi-objective protein design framework built on principles drawn from evolutionary biology. We apply dEVA to the zero-shot, *de novo* design of metalloenzymes by optimizing the coordination sphere of catalytic metals. We characterize a bi-zinc metalloenzyme that exhibits promiscuous hydrolytic activity towards both phosphomonoesters and phosphodiesters. This design achieves a rate enhancement ((k_cat_/K_M_)/k_w_) up to 3 × 10^13^, comparable to characterized natural phosphatases. dEVA offers a general and modular strategy for the programmable design of protein function without dependence on natural templates or evolutionary information.

## Main Text

Enzymes catalyze chemically demanding reactions with rate accelerations spanning many orders of magnitude (*1, 2*). Achieving high levels of catalytic proficiency comparable to natural enzymes remains a long-standing challenge in computational protein design (*3-5*). Early attempts to design biocatalysts resulted low initial catalytic rates, requiring subsequent directed evolution (*6-9*). Current approaches leverage deep learning and apply a process called motif scaffolding, in which a set of residues proposed to directly facilitate catalysis (i.e., the catalytic motif) is computationally generated or directly extracted from natural enzyme structures and placed into designed protein backbones (*10-12*). While this workflow is intended to preserve the motif geometry to hopefully recapitulate catalytic function, optimizing for the motif alone does not guarantee catalytic competence.

Enzyme activity, however, is well established to extend beyond the catalytic motif (*13-15*). Second-shell residues around the motif play an important role in mediating affinity, specificity, and reactivity (*16*). These contributions are well documented in natural enzymes but are largely absent in motif scaffolding approaches (*17*). Broader methods that attempt to capture the likely determinants of protein function either optimize for sequence-structure compatibility with codesign models (*18-20*), iteratively optimize for structure prediction metrics (*21-23*), or incorporate additional scores via either weighted sum (*24*) or sequential filtering (*11-12,25*). The resulting designs often favor features that score well across individual metrics or optimize for a single score (*26,27*). We hypothesized that a framework capable of jointly satisfying sequence constraints alongside the physicochemical requirements of the active site would enable the design of *de novo* enzymes that catalyze these chemically demanding reactions.

We were inspired by how nature evolved highly proficient function: through iterative selective pressure. We developed dEVA (design by EVolutionary Algorithm), a general and modular protein design protocol that simultaneously optimizes desired properties within a given structure. dEVA is built on the non-dominated sorting genetic algorithm (NSGA-II) (*28*), a multi-objective optimization framework that iteratively enriches for sequences that best satisfy competing objectives (**Methods, Fig. 1A, Fig S1**). Given a structure and a set of objective functions, dEVA explores sequence space to find an optimal solution. These design objectives can be defined as scoring functions or deep learning models, tailored to the desired properties.

**Fig. 1.**
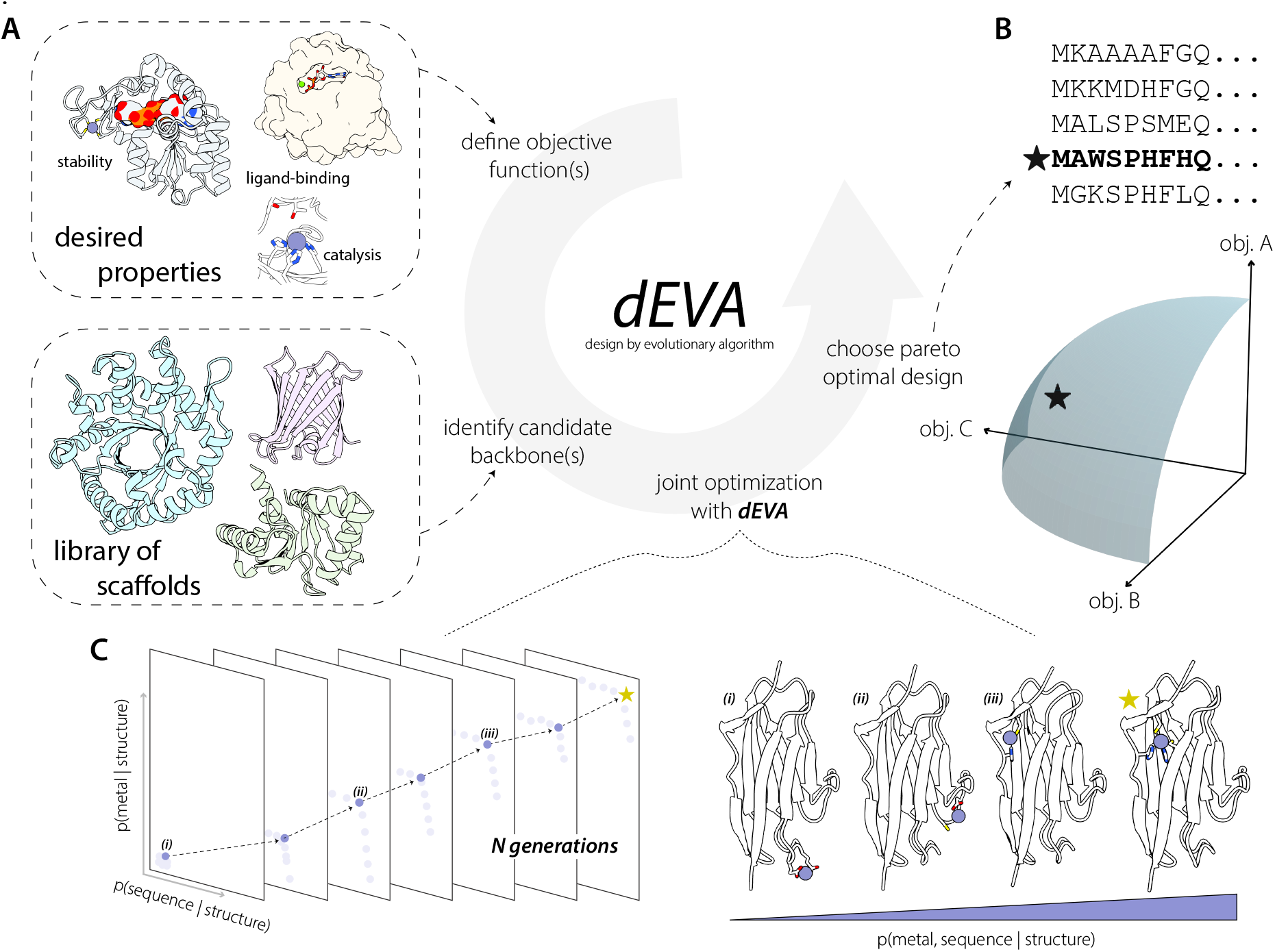
Overview of dEVA. (**A**) This multi-objective algorithm is a fixed-backbone, multi-objective optimization algorithm that can incorporate any number of defined objectives. (**B**) Optimal solutions are identified at the knee point (star) along the Pareto front. (**C**) dEVA follows a standard non-dominated genetic sorting algorithm (NSGA-II) (*28*) where the initial population of sequences is generated, evaluated across multiple objectives, and iteratively refined through mutation and crossover. The knee point represents the optimal tradeoff between sequence likelihood and metal-binding probability. Illustrated are representative designs sampled from along the Pareto front. Moving from *i* to *iii*, the predicted metal-binding geometry improves as the solutions approach the knee point, with *star* representing the optimal design at the knee point.

dEVA operates zero-shot: it imposes no requirement for structure prediction, natural templates, nor predefined motifs to generate designs, although these can be incorporated as additional objectives. Rather than optimizing for a single score, the designs favor solutions across them all. Optimal solutions are identified as designs representing the best, balanced tradeoffs across all defined objectives, referred to as the knee point of the “Pareto front” (**Methods, Fig 1B**).

Here, we sought to apply dEVA to design a *de novo* metalloenzyme. In metalloenzymes, the catalytic motif is commonly defined as the residues directly coordinating the metal. Previous attempts at designing metalloproteins have leveraged either motif scaffolding or rational design to achieve the desired metal binding (*29-34*). A key attribute that has remained difficult is incorporating second-shell interactions into the designs (*35*). Metalloenzymes impose an additional layer of complexity wherein the metal must occupy a catalytically competent coordination sphere to facilitate the chemical reaction, stabilize a nucleophile, or mediate electron transfer (*36*). We started by developing dEVA’s ability to design metal binding sites as a foundation from which we addressed the more demanding challenge of designing metalloenzymes.

### The design of proteins with *de novo* metal-binding sites

We began by posing metalloprotein design as a fixed-backbone design problem, where the goal is to identify sequences and metal configurations that are mutually adapted given a structural scaffold. This is achieved by defining two objectives in dEVA: *p(sequence* | *metal, structure*) and *p(metal* | *sequence, structure*). For the former sequence objective, we used LigandMPNN (*37*), trained to consider non-protein atoms. For the latter metal design objective, we used Metal3D (*38*) to predict metal location from the local chemical environment. At each iteration of the dEVA design protocol, LigandMPNN proposes mutations and sidechain conformations followed by Metal3D predicting the location of the metal ion(s). After *N* iterations, the final population reaches the Pareto front (**Fig. 1C, Fig. S2**).

We generated a diverse library of non-all-helical scaffolds using Protpardelle (*39,40*) and applied dEVA to unconditionally design metal-binding sites. From those, we selected eight designs spanning a range of lengths, topologies, and coordination motifs for in-depth experimental characterization (**Methods, Fig. S3-S5, Table S1-2**), where the full sequence and proposed metal-binding site was designed entirely by dEVA. All eight proteins were well-expressed in *E. coli* and purified as soluble monomers confirmed by SDS-PAGE and size exclusion chromatography (SEC) (**Fig. S6**). Characterization of all eight designs—including binding affinities, alanine point mutants, circular dichroism (CD) spectra in apo and holo states, and inductively coupled mass spectrometry (ICP-MS) data—is reported in the Supplementary Information (**Figs S7-S9, Table S3**).

Among all the designs, our best is desH2C2, an immunoglobulin-like fold in which dEVA designed a His_2_Cys_2_ coordination site without a motif template (**Fig. 2A-B**). desH2C2 binds zinc tightly (K_d_ = 37±4 nM, **Fig. 2C**) and ICP-MS confirms 1:1 stoichiometry (**Table S3, Fig 3D**).

**Fig. 2.**
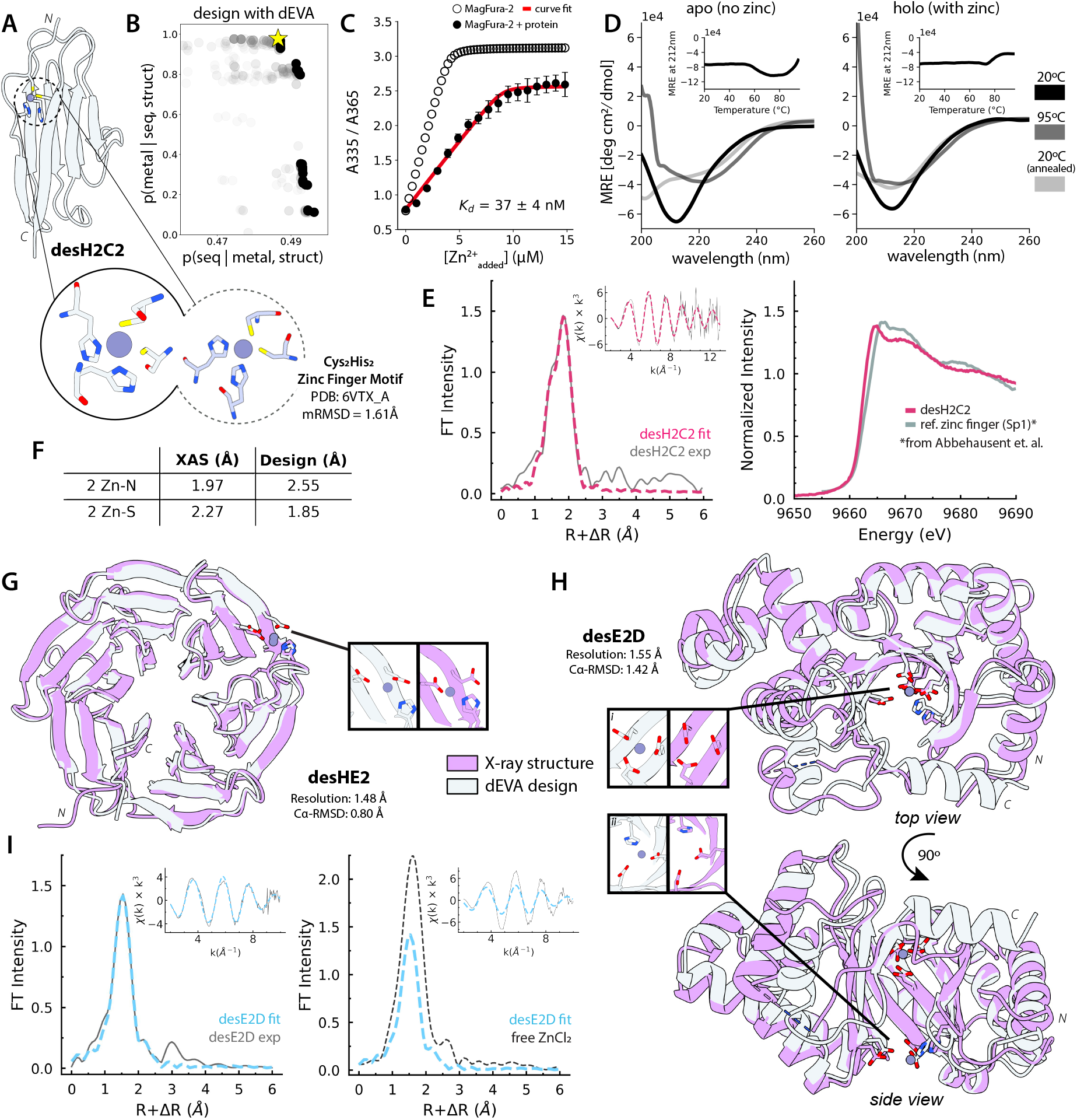
Designing de novo metalloproteins. (**A**) Cartoon representation of the designed metalloprotein desH2C2, where the gray circle represents the modeled zinc. The blow-out compares the designed motif (left) and its nearest neighbor zinc-finger motif in the PDB (right).(**B**) The yellow star indicates the chosen design from dEVA, which is also the Pareto-optimal solution sitting along the Pareto front. (**C**) The protein binds zinc tightly (K_d_=37±4nM) determined by competition titration against MagFura-2. Open circular dots represent experimental curve of MagFura-2 alone, black dots represent average experimental value, and red line represents best competition binding curve fit. Error bars represent relative error across n = 5 replicates. (**D**) Circular dichroism spectra with inset are thermal denaturation curves. The design is unstable in the apo form but shows increased thermostability in the presence of zinc.(**E**) X-ray absorption spectroscopy (XAS) of desH2C2 confirms metal coordination geometry. Comparison of the normalized Zn K-edge spectrum of desH2C2 (pink) with a digitized canonical zinc finger motif (*41*) (grey) suggests the design is a *de novo* binding motif. (**F**) The fits are consistent with 2 histidines and 2 cysteines and closely match the design. (**G**) Crystal structure of (magenta) superimposed with design model (white) for desHE2 (PDB: 12WO, 1.48Å resolution, Cα-RMSD 0.80Å) and (**H**) desE2D (PDB: 12WN, 1.55Å resolution, Cα-RMSD 1.42Å). (**I**) Zn K-edge XAS spectra of desE2D confirms that zinc is bound to the protein ligands despite not being visible in the crystal structure.

**Fig. 3.**
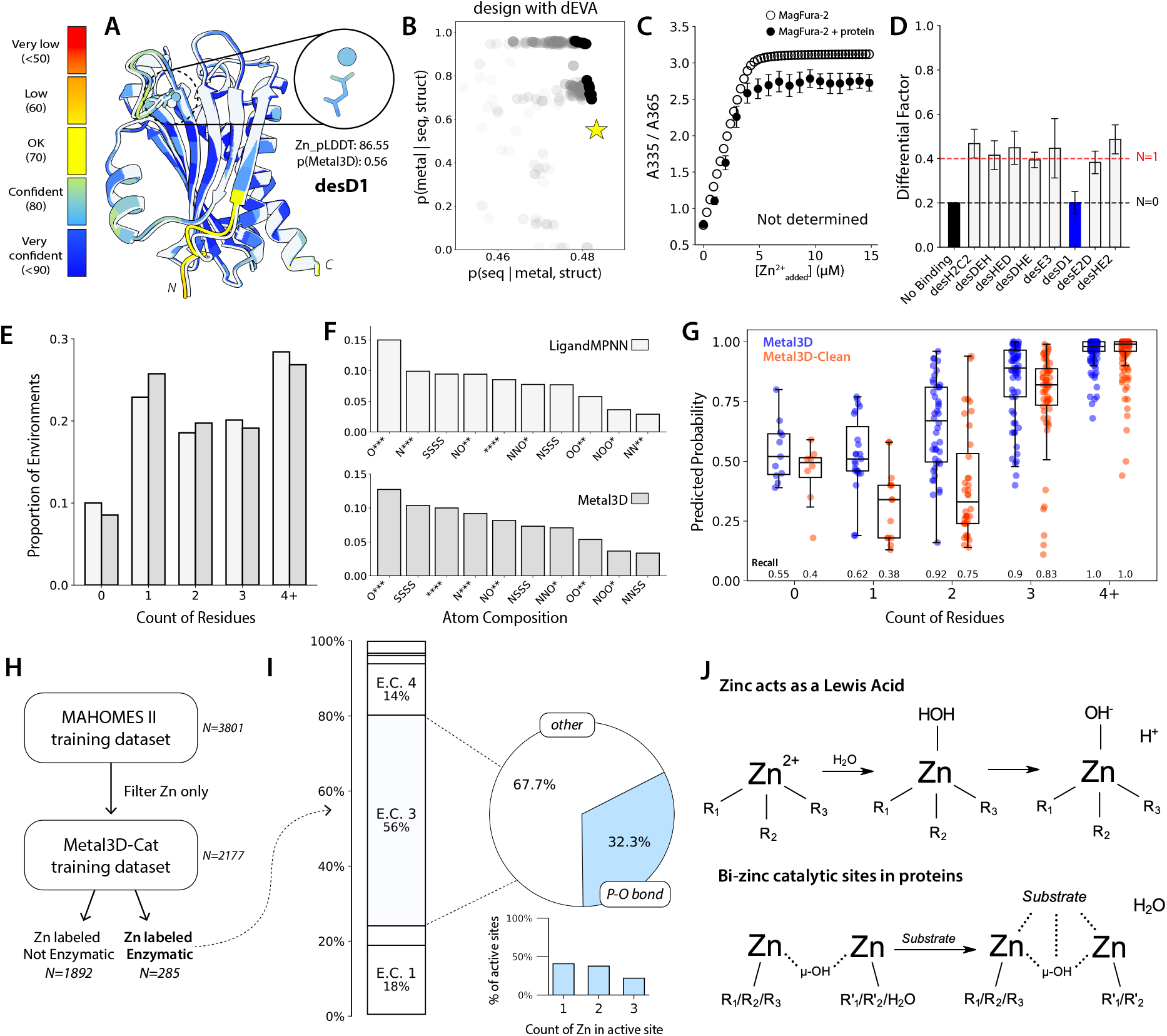
dEVA is sensitive to its models’ underlying training data. (**A**) AlphaFold-3 assigns high confidence to single ligand-coordinated zinc as indicated by desD1. Model in white overlaid with the predicted model, colored by pLDDT. (**B**) dEVA assigns the designed structure as a low-probability metal binding site. (**C**) The design has no detectable binding by MagFura-2 competition titration and (**D**) no detectable binding in ICP-MS. Error bars represent relative error for n=5 and n=3 replicates, respectively. (**E**) >54% of protein structures in the PDB have two or less coordinating ligands within 3.0Å of the zinc. (**F**) The majority of these structures have a zinc coordinated by a single carboxylate. The star (*) represents an open coordination sphere. (**G**) Retraining Metal3D (blue) by removing all metals coordinated with two or fewer ligands led to Metal3D-Clean (orange), reducing recall for under-coordinated metals while retaining recall for well-coordinated sites. Points represent all data with predicted probability greater than 0.0; boxes indicate IQR with median; whiskers extend to 1.5xIQR. (**H**) Curation of an enzymatic zinc training dataset. (**I**) Zinc sites span enzymatic classes, with the majority exhibiting hydrolytic behavior, including hydrolysis of a phosphate-oxygen bond. (**J**) Zinc act as a Lewis acid, and bicatalytic sites can coordinate and facilitate hydrolysis.

Zinc confers thermal stability absent in the apo form by CD (**Fig. 2D**). To resolve the local metal environment, we turned to X-ray absorption spectroscopy (XAS), a powerful solution-state structural technique that characterizes the metal coordination shell. For desH2C2, the Zn K-edge extended X-ray absorption fine structure (EXAFS) established coordination number and distances to 2 Zn-N and 2 Zn-S (**Fig. 2E, Table S4**). The normalized Zn K-edge spectrum reveals subtle whiteline differences compared to canonical zinc fingers (*41*) (**Fig. 2F**), potentially supporting the design’s distinct structural context and rotamer orientations. Together, these results suggest that dEVA generated a *de novo* coordination geometry, distinct from structurally characterized, zinc-containing proteins in the Protein Data Bank (PDB).

Crystal structures of three other designs indicate that dEVA can generate sequences for diverse and complex backbones independent of structure prediction (**Table S10**). desHE2, a *de novo* asymmetric β-propeller, was solved to 1.48Å with the backbone closely matching the designed model (PDB: 12WO, **Fig. 2G**). The designed zinc ion was resolved in the electron density and the metal positioned within 0.90Å of the designed location. desE2D, a *de novo* half-open TIM barrel, diffracted at 1.55Å and showed excellent backbone agreement with the design (PDB: 12WN, **Fig. 2H**), with deviations localized to a less ordered C-terminus. Likewise, a crystal structure of desDEH further confirms backbone fidelity (PDB: 12WM, **Fig. S10A**). In the latter two cases, the metal site was ambiguous in the electron density. However, XAS confirms the presence of zinc, with spectral features distinct from free ZnCl_2_; the EXAFS further shows shorter first-shell distance to free ZnCl_2_, confirming protein-bound metal (**Fig. 2H, Fig. S10B**), with ICP-MS indicating 1:1 stoichiometry (**Fig. 3C**). The EXAFS oscillation amplitude is lower show lower intensities than the desH2C2 design (**Fig. 2I**), suggesting ligation by lighter atoms (nitrogen or oxygen) consistent with the designed atom composition (**Table S4, Fig. S10C**).

As a negative control to probe the boundaries of dEVA’s design capabilities, we examined desD1 where the chosen design is far from the knee point (**Fig. 3A-B**). The designed site contains only a single ligand within binding distance—a coordination environment insufficient for stable metal chelation. Experimentally, there is no detectable binding; competition titration against MagFura-2 showed no apparent zinc binding, and ICP-MS confirmed no zinc association (**Fig. 3C-D**).

### dEVA is sensitive to the models’ training data composition

Despite our designs showing experimental success and consistency with the designs, we expected dEVA to create high affinity sites because of its additional consideration of the second-shell environment. However, our results show that most of the designs have weak affinity, suggesting that something was not adequately captured during the design protocol. Post-hoc, we predicted the structure of our negative control design desD1. Despite excellent overall structural agreement with the design, AlphaFold-3 (*42*) assigned high-confidence to the predicted metal (Zn pLDDT: 86.55) and positioned it in the same place as the low-confidence design (**Fig 3A**). By standard computational metrics and filtering criteria, this design would have been classified as a success; the ambiguity between the experimental data and structure prediction prompted us to evaluate what these models learned by examining their training data.

We began with the structural training datasets of both LigandMPNN and Metal3D—the two models used for dEVA metalloprotein design—and analyzed the coordination environments within 3.0Å of all annotated zincs (**Fig. S11-S12**). Across both datasets, we found that approximately 10% of sites had zero coordinating ligands, and over 54% had two or fewer residues (**Fig. 3E**). The distribution of the atom compositions reflects this heterogeneity; the most prevalent coordination pattern is a single, monodentate oxygen with three unoccupied coordination sites, while well-defined tetrahedral environments and catalytic sites are comparatively rare (**Fig. 3F**). Closer inspection of the depositions reveals that many of the poorly coordinated sites may reflect crystallization additives or soaking agents rather than biologically relevant co-factors. Zinc ions exhibit non-specific surface binding, particularly near aspartates, glutamates, and histidines (*43*). As a result, some depositions report over 20 resolved zinc ions in their electron density (e.g. PDB: 2EJC), while others contain only a few zinc ions tightly associated with just one ligand (e.g. PDB: 3IVB) (**Fig. S13**). These artefacts explain the disproportionate prevalence of carboxylate-mediated zincs in deep learning training datasets and the resulting confident predictions.

To address this directly, we trained Metal3D-Clean and excluded all zinc sites with fewer than three coordinating residues within 3.0Å. Evaluating Metal3D and Metal3D-Clean on the same held-out test set reveals a clear and expected tradeoff: Metal3D-Clean assigns substantially lower predicted probabilities and recall values for poorly coordinated sites (0-2 residues) while preserving performance on well-coordinated sites (**Fig. 3G**). As model behavior is directly shaped by training data composition, dEVA’s designs are therefore sensitive to the data its underlying models are trained on.

### The design of a *de novo* bi-zinc metalloenzyme

We next sought to exploit this sensitivity by tailoring the objectives for dEVA to enable the zero-shot design of a functional metalloenzyme. dEVA therefore needed a model that approximates *p(catalytic metal* | *sequence, structure*). This compelled us to train Metal3D-Cat on only annotated, catalytic zincs from MAHOMES-II (*44*) to discriminate the first- and second-shell chemical environments that govern zinc-mediated catalysis (*35, 45*) (**Methods, Fig. 3H**).

Analysis of the training data indicates that over half of the annotated zinc sites belong to hydrolases; of these, the largest fraction catalyzes the hydrolysis of phosphate-oxygen bonds, one-third of the total (**Fig. 3I**). While most hydrolytic active sites employ a single catalytic zinc, many zinc-containing enzymes use a bi- or tri-nuclear zinc center. In mono-nuclear zinc hydrolases, zinc acts as a Lewis acid to activate a water; in binuclear-zinc sites, the two metal centers play complementary roles in activating the nucleophile, orienting the substrate, and stabilizing the charge in the transition state (*46, 47*) (**Fig. 3J**).

To design a *de novo* metalloenzyme, we began by constructing a library of unconditionally generated backbones and filtered to a subset of TIM-barrel scaffolds—a fold whose functional plasticity has made it abundant across all six enzyme classes (*48*) For each scaffold, an initial full sequence was designed with ensemble Caliby (*49*), a Potts-based sequence design model that conditions on synthetic structural ensembles to remove non-structural sequence bias. Then, dEVA was employed to explore catalytic zinc configurations within the barrel cavity. dEVA identified only three Pareto-optimal solutions; these three designs (**Fig. S14**)—desA, desB, and desC—were individually screened for hydrolytic activity against phospho-ester substrates (**Fig. 4A**), reflecting the chemical reaction most represented in the Metal3D-Cat training data.

**Fig. 4.**
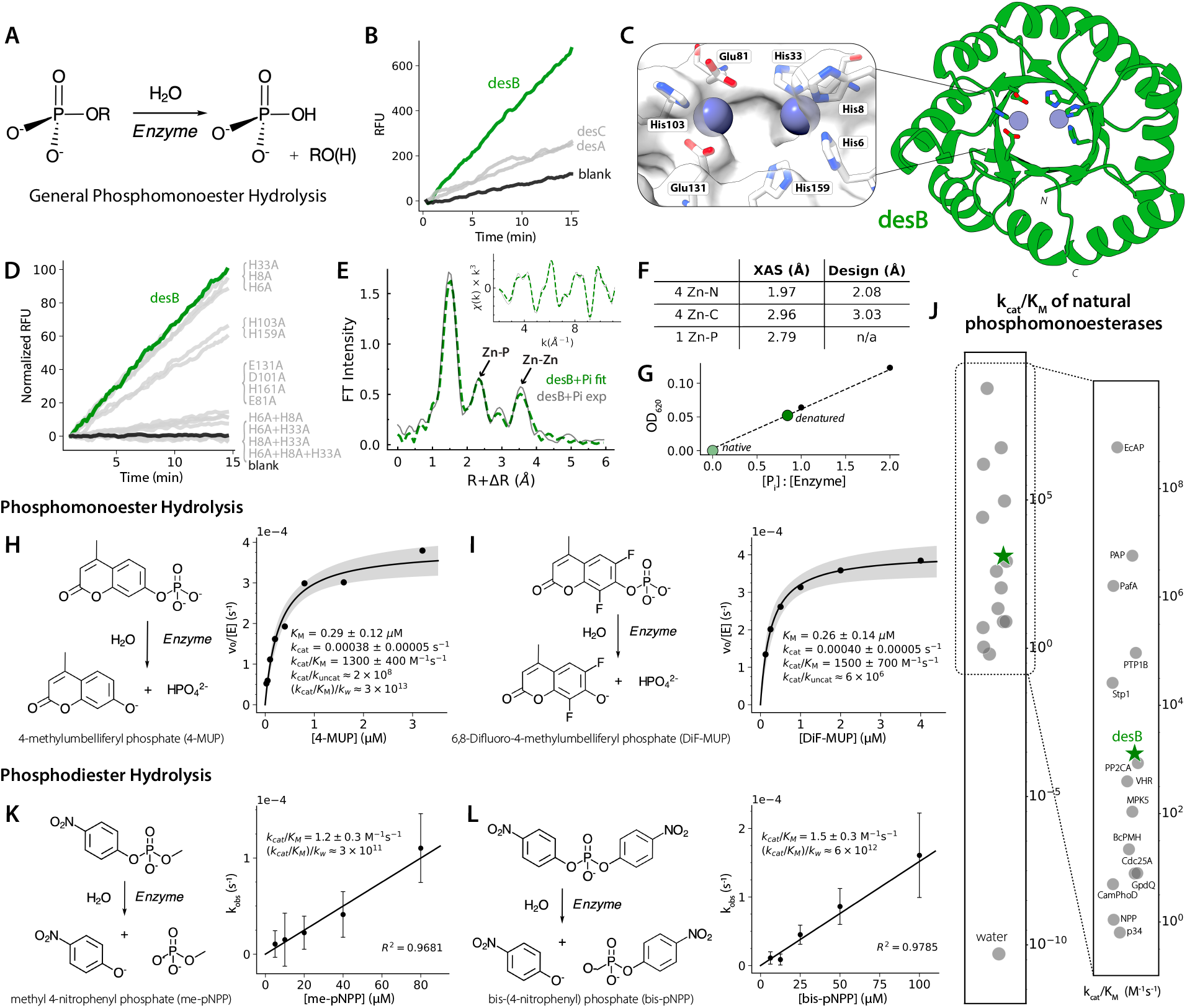
Zero-shot design of a de novo binuclear zinc metalloenzyme. (**A**) General mechanism of phosphomonoester hydrolysis (**B**) Initial fluorescence-based screen of desA, desB, and desC against 4-MUP; desB (green) shows the highest activity. (**C**) Cartoon representation of the dEVA design model for desB, a *de novo* TIM barrel harboring a bi-zinc active site (blue spheres). (**D**) Screen of the alanine mutants of first- and second-shell active site residues confirms catalytic dependence on the designed bi-zinc site. (**E**) Zn K-edge EXAFS of desB. Structural analysis of the bi-zinc active site; experimental (black) and fitted (green) spectra fit with a Zn-Zn scattering path at 4.3Å (see **Table S4**). (**F**) Comparison of EXAFS distances and design model. (**G**) Malachite green phosphate assay; phosphate is released only upon denaturation and scales proportionally with enzyme concentration. (**H, I**) Michaelis-Menten kinetics of desB for phosphomonoester substrates 4-MUP and DiF-MUP. Shaded region represents ±1 standard error. Measurements were performed in biological triplicate. (**J**) Comparison of desB catalytic efficiency (k_cat_/K_M_) against literature values for natural phosphomonoesterases. (**K, L**) Linear kinetics of desB for me-pNPP and bis-pNPP. Measurements were performed in biological duplicate.

Of the three candidates, desB emerged as the most active, catalyzing phosphomonoester hydrolysis with a rate significantly above background (**Fig 4B**). This is particularly notable as desB harbors a bi-nuclear zinc site (**Fig. 4C**)—the same bi-zinc site that underlies the exceptional efficiency of many naturally occurring metallohydrolases (*50*). We proceeded to characterize this design.

desB is well-folded and monomeric, exhibiting high thermostability as assessed by SEC and CD, respectively (**Fig. S15-S16A**). The zinc site of desB has an entirely *de novo* motif; motif searches against the PDB using both Folddisco (*51*) and RCSB Structure Motif Search (*52*) returned no structural motif neighbors, and BLAST (*53*) returned no sequence homologs. The novelty of this active site is further supported by the absence of bi-zinc sites among structural neighbors identified by Foldseek (*54*).

To establish that activity depends on the designed metal-binding site, we checked that EDTA chelation reduces activity (**Fig. S16B**). However, metal chelation also destabilized the protein (**Fig. S16C**), precluding determination of zinc-binding affinity. Alanine scanning mutagenesis of the first- and second-shell residues showed that mutations to the designed site reduced activity (**Fig. 4D, Fig. S17, Table S6**). Metal stoichiometry was independently determined by ICP-MS, confirming that desB coordinates two zinc ions per monomer (**Table S7**). pH profiling revealed optimal activity under alkaline conditions (**Fig. S18)**, consistent with a binuclear zinc site in which two proximal zinc ions lower the pKa of metal-bound water to ∼7, thereby generating a hydroxide nucleophile for attack at alkaline pH (*55, 56*) (**Fig. 3J**).

We then used XAS with orthophosphate as a ground-state analog to probe the catalytic configuration of the zinc site. desB showed high X-ray fluorescence counts that is consistent with two zinc ions. Notably, the Zn-Zn distance detected in solution (∼4.3Å) falls within the range of observed catalytically competent binuclear zinc hydrolases (*50*). Fitting the Zn K-edge EXAFS yielded scattering paths consistent with 4 Zn-N, 1 Zn-P, and 1 Zn-Zn interactions (**Fig. 4E-F, Fig. S19**). The prominent Zn-P contribution indicates direct coordination of phosphate to the metal center and suggests a closely associated, bridged zinc configuration, consistent with phosphoryl-transfer catalysis. Phosphate binding was independently confirmed by a malachite green assay, with bound phosphate detected only upon denaturation (**Fig. 4G, Fig. S20**). Phosphate inhibits desB activity with a K_i,app_ of 1.0±0.2µM, similar to the inhibition constants reported for a subset of natural alkaline phosphatases (*57*) (**Fig. S21**).

The hydrolysis of phosphomonoesters is among the most energetically demanding reactions in biology, with uncatalyzed half-lives >500,000 years (*1*). desB overcomes this barrier with catalytic efficiencies (k_cat_/K_M_) of 1300±400 and 1500±700 M^-1^s^-1^ for 4-MUP and DiF-MUP, respectively (**Fig. 4H-I**). Compared to literature values, desB activity is within the range of natural phosphomonoesterases (**Fig. 4J, Table S8**). Its rate enhancement ((k_cat_/K_M_)/k_w_) is 3 × 10^13^ for 4-MUP, the highest reported rate enhancement for any *de novo* designed hydrolase to date (*33*). desB continually turns over substrate, as confirmed by product accumulation over hours at sub-stoichiometric enzyme concentrations (**Fig. S22**).

Remarkably, desB also catalyzes the hydrolysis of phosphodiesters, a chemically more demanding reaction class (with uncatalyzed half-lives >13 million years (*1*)) distinguished by a different charge state and transition state geometry. desB hydrolyzes both me-pNPP and bis-pNPP with a rate enhancement of 3 × 10^11^ and 6 × 10^12^, respectively (**Fig. 4K-L**). Their k_cat_/K_M_ of 1.2±0.3 and 1.5±0.3 M^-1^s^-1^ makes desB also comparable to natural enzymes with characterized phosphodiesterase activity (**Fig. S23, Table S9**). This substrate promiscuity suggests that the catalytic efficiency is not limited by the size of the diester moiety, consistent with the original design of an exposed active site.

We solved the X-ray crystal structure of desB at 2.29Å resolution (PDB: 36BS), confirming the designed TIM-barrel fold (Cα-RMSD: 2.37Å) and revealing the binuclear zinc site in the electron density with the designed first-shell ligand chemistry intact (**Fig 5A-B, Fig. S24, Table S10**). The Zn-Zn separation of 5.9Å agrees closely with the design model (5.8Å), contrasting with the compressed 4.3Å distance observed upon phosphate binding. Comparison of the phosphate-free and phosphate-bound Zn K-edge XAS further shows that the two states differ in their zinc coordination environments (**Fig. 5C, Fig. S19, Table S4**). Together, these observations support a possible mechanism in which desB adopts an open, solvent-exposed inactive state and becomes catalytically competent upon ligand coordination at the binuclear zinc center. This behavior parallels proposed mechanisms of many natural binuclear metallohydrolases (*58*), including phosphoesterases, where substrate coordination drives formation of a zinc configuration with reduced Zn-Zn separation poised for catalysis.

**Fig. 5.**
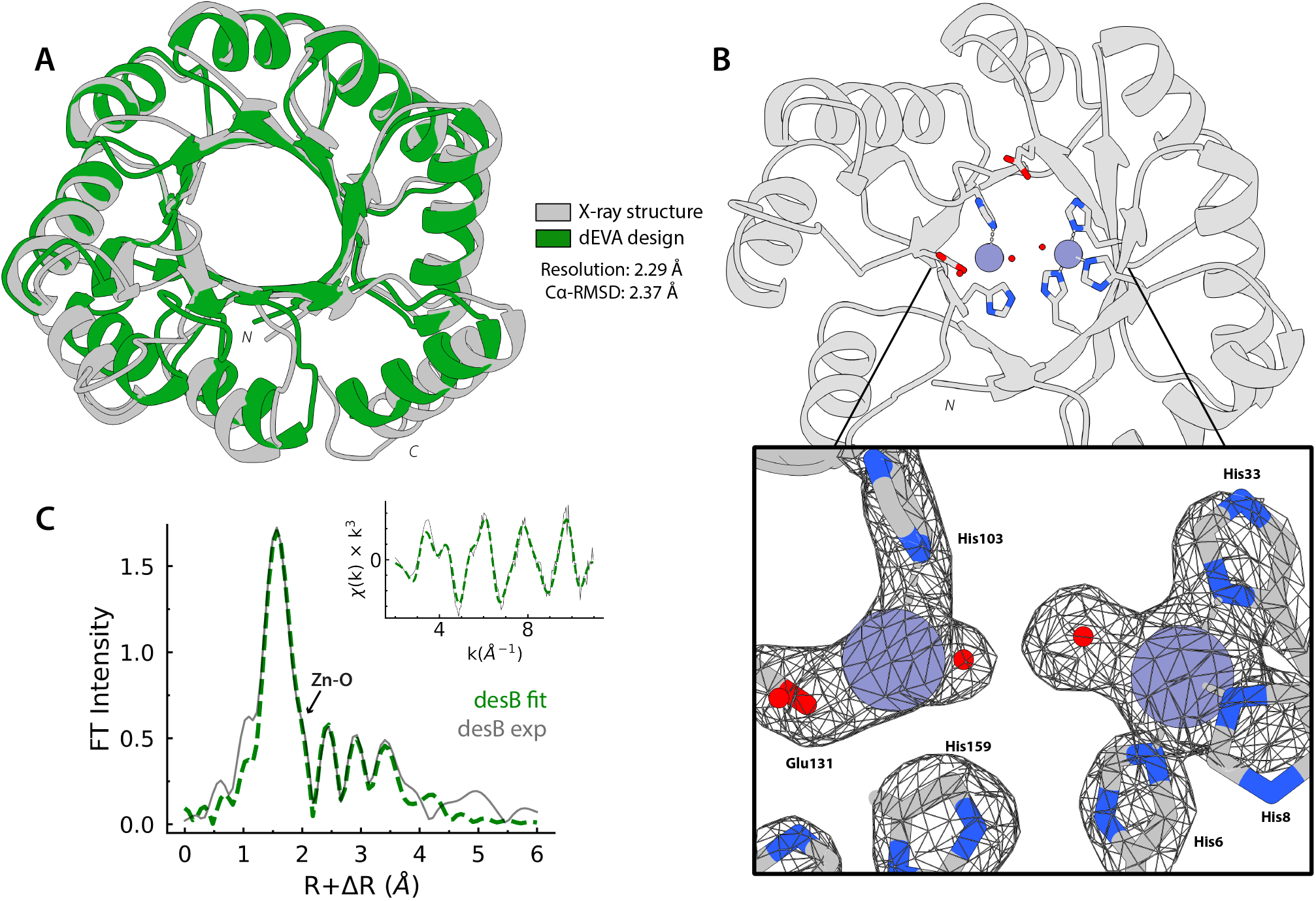
Crystal structure of desB matches closely with design model. (**A**) Crystal structure (gray) superimposed to the design model (green), showing close overall agreement (PDB: 36BS, 2.29Å resolution, Cα-RMSD 2.37Å). (**B**) Overview of the desB crystal structure highlighting the zinc coordination site (blue spheres) and coordinating residues and water ions (small red circles). Electron density map (2mFo-DFc, contoured at 2.0σ) of the zinc coordination site shows well-defined density around the coordinating residues and zinc ions, supporting the designed bi-zinc active site. (**C**) Zn K-edge EXAFS of desB supports the coordinating residues in the bi-zinc active site; experimental (black) and fitted (green).

## Discussion

The design of *de novo* enzymes represents one of the most demanding tests of deep learning methods. While previous enzyme designs have relied on scaffolding known catalytic motifs, this constrains solutions to geometries that evolution has already conceived. We demonstrate that this constraint is not necessary; by starting from no natural template, no theozyme geometry, and no evolutionary data, dEVA yielded a *de novo* metalloenzyme in zero-shot. The resulting design has no structural precedent and catalytic efficiency comparable to natural phosphatases—catalyzing two of the most energetically demanding reactions in biology.

This work draws inspiration from how natural selection forged highly proficient function. In nature, the evolution of enzymes is widely thought to have begun with “generalist” catalysts that, under millennia of selective pressure, became highly efficient and substrate-specific machines (*59*). Analogously, the design of *de novo* functional proteins may begin with the design of promiscuous catalysts as a bed-ground for future protein engineering efforts. desB is one such catalyst: its active site sits in a non-specific open cavity, and its promiscuity for model phosphomonoester and phosphodiester substrates is unpaired compared to the promiscuity of natural enzymes (*60*). This opens the door for future engineering of desB to tune its specificity and tailor its catalysis.

To navigate sequence space, dEVA similarly draws from evolutionary biology. Natural selection explores sequence space through iterative mutation and selection, implicitly sampling epistatic interactions. By leveraging a genetic algorithm, dEVA enables exploration of these interdependencies—though its success ultimately rests on both the quality of the training data and sensitivity of the models used for each defined objective (*61*). Our findings here represent a cautionary tale on training deep learning models without consideration of the physicochemical reality of macromolecules. Continued development of models that learn the biophysical and biochemical determinants of protein function will enable better formulation of the design objectives necessary for efficient catalysis. dEVA offers a starting point for balancing these objectives and we envision as a flexible platform for functional protein design. Ultimately, the design of diverse enzymes will benefit from accessing other catalytic solutions yet to be conceived by natural evolution.

## Acknowledgments

We thank Dan Herschlag for helpful discussions regarding alkaline phosphatases and for providing many of the reagents used in these experiments. We thank Alex Hoffnagle for help setting up metalloprotein metal-binding affinity characterization. We graciously thank Giovanni Aviles and Nicholas Freitas (UCSF and UC Berkeley) for providing plasmid to use as controls for setting up initial experimental assays. We graciously thank Micah Olivas (Fordyce Lab, Stanford) and Mikkel Madsen (Lin Lab, Stanford) for providing reagents and substrates for experimental assays. We thank Richard Shuai for running the ensemble Caliby scripts for the initial sequence designs, as well as Paul Rujigrok, Jingjia Liu, and Carla Perez for helpful discussions.

G.E.N. acknowledges funding from National Science Foundation Graduate Research Fellowship (NSF GRF). S.L.D acknowledges funding from an EPFL-Stanford Firmenich Fellowship. This research was funded in part by the Swiss National Science Foundation [200020_219440]. F.S. acknowledges funding by NIH (R01 GM064798). Additional support for this research comes from Merck Research Laboratories (MRL) Scientific Engagement and Emerging Discovery Science (SEEDS) Program and NIH (R01GM147893) awarded to P.-S.H. and Department of Energy-Basic Energy Sciences Field Work Proposal 101018 awarded to R.S.

Work was performed in part in the Stanford SIGMA Facility with support from the Stanford Doerr School of Sustainability and Stanford Nano Shared Facilities (SNSF)/Stanford Nanofabrication Facility (SNF) under National Science Foundation award ECCS-2026822 (RRID: SCR_02329), and at the Vincent Coates Foundation Mass Spectrometry Laboratory, Stanford University Mass Spectrometry (RRID:SCR_017801) utilizing the Thermo Exploris 240 LC/MS system (RRID:SCR_022216). Use of the Stanford Synchrotron Radiation Lightsource, SLAC National Accelerator Laboratory, is supported by the U.S. Department of Energy, Office of Science, Office of Basic Energy Sciences under Contract No. DE-AC02-76SF00515. The SSRL Structural Molecular Biology Program is supported by the DOE Office of Biological and Environmental Research, and by the National Institutes of Health (NIH), National Institute of General Medical Sciences (P30GM133894). The contents of this publication are solely the responsibility of the authors and do not necessarily represent the official views of NIGMS or NIH. For the purposes of Open Access, a CC BY public copyright license is applied to any Author Accepted Manuscript (AAM) version arising from this submission.

## Author Contributions

G.E.N., S.L.D., U.R., and P.-S.H. conceived the research. G.E.N. and S.L.D. built the dEVA algorithm. G.E.N. performed model training, computational evaluations, and generated designs. G.E.N., and Q.W. performed experimental characterization. G.E.N. and F.S. performed enzyme characterization. I.I.M. solved the crystal structures. Y.H. and R.S. performed and interpreted XAS experiments. G.E.N. performed data analysis and visualization. P.-S.H., F.S., U.R., and R.S., supervised the research. G.E.N. and P.-S.H wrote the manuscript with assistance from all co-authors.

## Code availability

dEVA and all associated model weights are available as a modular framework via GitHub at https://github.com/ProteinDesignLab/dEVA.

## Data availability

All data are available in the main text or as supplementary materials, including all reference literature values and ordered protein sequences and mutants. Protein crystal structure coordinates and structure factors are available in the Protein Data Bank with PDB IDs 12WM, 12WN, 12WO, and 36BS.

