## Supplementary Material for "Zero-shot design of a *de novo* metalloenzyme"

**Supplementary Material for**
**Zero-shot design of a *de novo* metalloenzyme**

Gina El Nesr[1], Simon L. Dürr[2,3,4], Irimpan I. Mathews[5], Qi Wen[4], Kewei Zhao[5], Ritimukta Sarangi[5], Ursula Rothlisberger[2], Fanny Sunden[6], Po-Ssu Huang[4]

[1]Biophysics Program, Stanford University, Stanford, CA
[2]Institute of Chemical Sciences and Engineering, École Polytechnique Fédérale de Lausanne (EPFL), Lausanne, Switzerland
[3]Institute of Life Sciences, HES-SO Valais-Wallis, Sion, Switzerland
[4]Department of Bioengineering, Stanford University, Stanford, CA
[5]Stanford Synchrotron Radiation Lightsource SLAC National Accelerator Laboratory, Menlo Park, CA
[6]Department of Biochemistry, Stanford University, Stanford, CA

#### Materials and Methods

Table of Contents


**Introduction to Supplementary Materials** **5**
**Computational algorithms and model training.**
*The dEVA algorithm.* *5*
*Identifying crystallographic artefacts in deep learning training datasets.* *6*
*Metal3D-Clean* *7*
*Evaluating Metal3D and Metal3D-Clean* *7*
*Metal3D-Cat* *8*
*E.C. distribution of Metal3D-Cat training dataset* *8*
*Identifying similar sequences, structures, and motifs* *8*
**Phase I: Metalloprotein design and characterization** **10**
*Computational design of metalloproteins.* *10*
*Metalloprotein protein expression and purification.* *10*
*Single-site alanine mutagenesis.* *11*
*Circular dichroism and thermal denaturation.* *11*
*Metal-binding affinity by MagFura-2 competition titration.* *11*
*Cobalt Substitution as a Spectroscopic Probe of Metal-Binding.* *12*
*Determination of zinc content by ICP-MS.* *12*
**Phase II: Metalloenzyme design and characterization** **14**
*Computational design of metalloenzymes.* *14*
*Metalloenzyme protein expression and purification.* *14*
*Single-site alanine mutagenesis.* *16*
*Kinetic analyses.* *16*
Initial design screen. 17
Kinetic screen under variable pH. 17
Kinetic screen under variable temperature. 18
Kinetic screen under variable ionic strength. 18
Phosphomonoesterase: 4-MU standard curve. 19
Phosphomonoesterase: 4-MUP kinetics. 19
Phosphomonoesterase: DiF-MU standard curve. 19
Phosphomonoesterase: DiF-MUP kinetics. 20
Phosphomonoesterase: desB mutants. 20
Phosphodiesterase: me-pNPP kinetics. 20
Phosphodiesterase: bis-pNPP kinetics. 20
Michaelis-Menten kinetics. 21
Determining Apparent Inhibition Constants ($K_{i,app}$). 22
*Attempting to prepare apo-desB by zinc chelation.* *22*
*Circular dichroism and thermal denaturation.* *22*
*Preparation of zinc-supplemented sample for ICP-MS and XAS.* *22*
*Determination of zinc content by ICP-MS.* *23*
*Quantifying free orthophosphate in native and denatured desB.* *23*
*Liquid-chromatography mass spectrometry (LC-MS).* *24*
**Structural determination methods.** **25**
*X-ray crystallography.* *25*
*X-ray absorption spectroscopy (XAS).* *26*

Additional preparation of the metalloproteins. 26
Additional preparation of metalloenzyme desB. 27
Performing XAS. 27

Introduction to Supplementary Materials

The principal hypothesis motivating our two-phased approach is that the design of robust metal-binding sites is essential for the design of catalytic metal sites. We therefore separated the design process (and describe the methods associated with the design processes) into two phases: the design of metalloproteins and the design of metalloenzymes, henceforth referred to as Phase I and Phase II. These ultimately distinguish the difference in computational approaches and experimental purification/characterization strategies. Structural characterization methods via x-ray crystallography and x-ray absorption spectroscopy are reported separately.

We chose to combine all the computational algorithm development, model training, and computational evaluations discussed in the manuscript into one separate section, as we believe these algorithms and their respective evaluations standalone as advances in the computational modeling of proteins, broadly.

Proteins referred to as "metalloproteins" represent the non-catalytic, initial eight designs used to probe metal-binding design capabilities. Proteins referred to as "metalloenzymes" represent the catalytic designs used to probe metalloenzyme capabilities.

Computational algorithms and model training.

The dEVA algorithm.

dEVA is implemented using Python. The algorithm requires defining at least two objectives and defining an objective evaluation file in order of evaluation and mutations. Each file must contain an initialization and scoring function, both of which are tailored for the objectives. The score is not restricted to bounded intervals or defined by an upper maxima and lower minima. More information can be found on GitHub in the README.md of https://github.com/gelnesr/dEVA. dEVA closely follows the procedures of a non-dominated sorting genetic algorithm II (NSGA-II) (*28*).

The specific pipeline used for design in this paper is illustrated in **Figure S1** and is as follows; An initial structure is parsed and LigandMPNN (*37*) and Metal3D (38*)* are used to calculate initial fitness of the starting sequence for the backbone and metal binding. Then the genetic algorithm procedure is initialized by generating *i* new sequences by sampling from the LigandMPNN distribution using sampling temperature 0.5 and LigandMPNN-OpenFold for rebuilding the rotamers of the new sequence. The population size *N,* the number of generations *M,* and conditioning information are user-defined starting parameters. For each individual *i* using the generated structure Metal3D is run and from the generated metal probability density the metal ions are added to the structure. The Metal3D default parameters (distance threshold 7 and max probability 0.1) are used. The maximum probability of any one of the metals predicted in the structure as well as the LigandMPNN score are used for the optimization. The parent sequence generated in this step as well as their fitness are then used for tournament selection to determine the sequences that will be used to generate *N* children.

For each parent among four randomly sampled sequence the best sequence is chosen according to crowding distance and tournament probability 0.9. The two parent sequences are then mixed using crossover exchanging one fragment of random length. Then 4 mutations were introduced in each sequence. After generating the children, *2N* individuals exist of which only *N* are selected for the next generation *m+1*. For this first non-dominated sorting is used to determine the non-domination rank of all individuals and sort them in Pareto fronts. The individuals for the next generation are chosen according to their non-domination rank until *N* individuals are chosen. First, all individuals with non-domination rank 0 are chosen, then all with rank 1 and so forth. If not all individuals with a given rank fit into the remaining individuals for the $m^{th}$-generation crowding distance comparison is used to choose individuals that best cover the fitness space. This procedure is iterated *M* times. For the results reported here we co-optimize the probability of sequence from LigandMPNN and probability of metal from Metal3D.

Integration with LigandMPNN (and ProteinMPNN) retain features including fixed and variable residue identities, omitted amino acid identities, and weighting of specific amino acid types at all or defined positions. Use of these features is described later. The code for LigandMPNN and ProteinMPNN was modified for better ease of use such generating a specific score function compatible with the pipeline. This version is made available on GitHub.

Extension to other fitness scores such as structure prediction metrics, hydrogen-bonding evaluation, or other metrics—mathematical equations, deep learning models, or otherwise— is trivial so long as a single floating-point score be attributed to the method. Computation time scales with complexity of compute for each specific metric and number of generations and individuals defined in the genetic algorithm. The expected asymptotic upper bound for an NSGA-II algorithm is $O(NK^2)$, where *N* is the population size and *K* are the number of defined objectives.

Identifying crystallographic artefacts in deep learning training datasets.

As described in main text, the experimental results compelled the re-evaluation of the training datasets used as objectives in Phase I metalloprotein design.
The training dataset for LigandMPNN was downloaded from https://github.com/dauparas/LigandMPNN/blob/main/training/train.json and the training dataset for Metal3D was downloaded from https://doi.org/10.5281/zenodo.7015849/ Both datasets contain PDB ID overlap in part because of their curation strategy.

Individual PDB files were downloaded via the RCSB REST API and zinc ions were identified from the pdb-formatted coordinate files by scanning all ATOM and HETATOM records for residue names matching the element symbol “ZN”. For each detected metal, the atomic coordinates (x,y,z), chain identifier, residue sequence number, occupancy, and B-factor were extracted directly from the file. Because pdb files vary in labeling of water molecules, all water molecules were ignored in subsequent occupancy evaluations to allow for standardization of the datasets. The first biological assembly was used, and no additional structures were generated from crystallographic symmetry. This approach evaluates the deposited PDB coordinates directly, consistent with common practices in structure-based protein modeling pipelines.

For each non-metal, non-water residues within a Euclidean distance cutoff of 3.0Å were identified as coordinating residues. Distances were computed between the metal center and every heavy atom of each candidate residue. The coordinating shell was summarized by recording the unique set of residues (by chain, residue sequence number, and residue name), the closest atom name per residue, and the total count of coordinating residues. Coordinating residue identifiers were formatted as concatenated residue name, sequence number, and chain (e.g. HIS64A), separated by semicolons. For each identified metal site, the total number of ligands coordinating the atom was calculated and recorded. Reported evaluations compare the count of residues as identified by the ligand atoms and residue identities described above. No statistics were computed beyond overall count per coordinating environment.

For each coordinating residue, the closest-approach atom was mapped to an element symbol: oxygen (O) for atoms with names beginning with OD, OE, OG, OH, or O; nitrogen (N) for ND, NE, NH, NZ, or N. sulfur (S) for SD, SG, or S. Patters were standardized to a four ligand coordination target by padding under-coordinated sies with a wildcard character (*) and truncated "over-coordinated" sites. Non-wildcard symbols were sorted alphabetically before concatenation to produce a canonical, order-independent pattern string (e.g. NNOS, NOSS, NS**).

<u>Metal3D-Clean</u>

The same PBD training dataset used in training the original Metal3D model was used, with different pre-processing, described here. Unlike the original Metal3D pipeline, protein structures were not additionally expanded from the asymmetric unit using crystallographic symmetry operators. For structures where metal coordination spans biological assembly interfaces, we assumed the first biological assembly would adequately capture these interactions where relevant. During training, zinc atoms with fewer than three coordinating residues (within a 3.0Å cutoff) were masked and excluded from the training objective entirely. The model was trained as described in the original Metal3D, but using a filter size of 4, and on an NVIDIA A100 for 10 epochs.

<u>Evaluating Metal3D and Metal3D-Clean</u>

Metal3D and Metal3D-Clean were evaluated against the metal positions found in the pdb files in the test set, with the distinction that Metal3D-Clean was trained on the further pre-processed dataset. For each metal site found in the proteins in the test set, the nearest predicted metal was identified by minimum Euclidean distance. A prediction was counted as a true positive if this distance fell within 2.0Å of the original metal site position. For true positives, the prediction confidence was recorded. Otherwise, the prediction was counted as not predicted. False positives were not considered in this analysis as the objective was to specifically evaluate recovery of crystallographically annotated metal sites (i.e., recall) rather than overall prediction precision.

To assess model performance as a function of coordination environment complexity, the true metal sites were stratified by the number of coordinating protein residues within 3.0Å, with sites having four or more coordinating residues grouped into a single bin (≥4). For each coordination bin, the per-model recall was computed as a proportion of true sites predicted within 2.0Å matching the distance threshold. Prediction confidence scores for the true positive sites were visualized as a jittered strip plot overlaid with box-and-whisker plots showing the median, interquartile range, and 1.5xIQR whiskers. Per-bin recall values were annotated.

Metal3D-Cat

The same PBD training dataset used in training MAHOMES (*44*) was used, with different pre-processing, described here. First, all identified proteins that did not contain zinc were discarded, filtering the entire dataset from 3,801 identified structures and chains to 2,177. The filtered training and test dataset split used in MAHOMES was used as is with no further preprocessing for the training of Metal3D-Cat.

Zinc metals that were identified as "enzyme" in the original training dataset comprised the positive training class; therefore, zinc metals that were identified as "non-enzyme" were masked during model training. Further training details of Metal3D-Cat are analogous to the training procedure of Metal3D-Clean, described above.

E.C. distribution of Metal3D-Cat training dataset

To characterize the functional diversity of the Metal3D training dataset, the EC numbers were retrieved programmatically for all unique PDB entries annotated as enzymes in the filtered Metal3D-Cat training dataset. For each entry, polymer entry identifiers were first obtained via the RCSB REST API, and the EC number was retrieved for each polymer entity. EC numbers recorded in the `pdbx_ec` field was parsed and normalized to handle both comma- and semicolon-delimited entries. Entries returning no enzyme class (EC) annotation was recorded as missing.

EC numbers were classified at both the first level (enzyme class, e.g. hydrolase, transferase) and second level (enzyme subclass) by splitting on the dot-delimited EC string. Counts were aggregated across all entities per structure. The bond type cleaved during hydrolysis was manually annotated for structures in the hydrolase class by parsing the reaction string for common bond patterns (P-O, C-O, C-N, etc). We note that many of these types of bonds could also be broken in other classes but were outside of the scope of this evaluation.

Identifying similar sequences, structures, and motifs

Sequence similarity searches were performed using the Basic Local Alignment Search Tool (BLAST). The query protein sequence was used to search against the NCBI non-redundant (nr) protein database using BLASTp (*53*) with default parameters. Hits were ranked based on E-value, percent identity, and alignment coverage. The sequence with the highest percent identity is reported where BLAST sequence similarity is indicated.

Structural similarity searches were conducted to identify nearest neighbors in the Protein Data Bank (PDB) using TM-score alignment from Foldseek (*54*). The output from deVA was used as the query structure and was compared against all available entries using TM-align (**Figure S4**). Structural alignments were performed using default parameters, and similarity between structures was quantified using TM-score. The sequence with highest structural similarity as evaluated by TM-score was reported. The RMSD reported per structure was calculated using CEalign.

Structural motif searches were performed to identify nearest neighbor active site architecture in the Protein Data Bank (PDB). The query structure was analyzed using Folddisco (*51*) and RCSB PDB Motif Search (*52*). Searches were conducted using default parameters for both methods. Identified matches were ranked based on structural similarity and geometric alignment of all motif residues (**Figure S5**). To ensure functional relevance, only the nearest neighbor motifs in which all query ligands were present were reported. Motifs lacking complete ligand representation or exhibiting incomplete coordination environments were discarded.

*Phase I: Metalloprotein design and characterization*

Computational design of metalloproteins.

An initial set of backbone structures was downloaded from Lu et. al.[6]. In brief, all samples (both "designable" and "undesignable" structures) from the Protpardelle (*39*, *40*) stepscale 0.8 set were used. Structures that lacked any beta-strand content or were >75% helical content as determined by DSSP (as calculated by BioPython) were discarded. One sequence per backbone for all remaining backbones was designed using the ProteinMPNN with default parameters and then further clustered by TM-score of 0.7 using Foldseek. All cluster representatives were taken as the initial design set and designed using the dEVA algorithm applied to metalloprotein design. CATH topology for the cluster representatives was determined using methods described in Eguchi et. al. (*62*). One design per CATH topology was chosen for each of the hypotheses described in the main text. This made up the final eight candidate structures that were further characterized in Phase I.

Prospectively, we found that without retraining Metal3D (and likely, LigandMPNN), most of the designed and converged metal binding sites were coordinated by two or more carboxylate residues.

Metalloprotein protein expression and purification.

Genes encoding the designed protein sequence (**Table S1A**) were synthesized and cloned into pET-24a(+) *Escherichia coli* plasmid expression vectors (Genscript, TEV cleavage tag, 6xHis-tag). Plasmids were then transformed into chemically competent BL21(DE3) E. coli (Zymo Research). The cells were cultured in 2xYT medium at 37 °C until the optical density (OD) reached 0.6–0.8. Protein expression was then induced with 1 mM IPTG at 16 °C. After overnight expression, cells were centrifuged (4,000xg, 30min, 4ºC) and resuspended with 50 mM NaPi 300 mM NaCl (pH 8.0) and frozen at −80 °C until extraction and purification.

The cell pellet was thawed and supplemented with final concentration of 1mM PMSF dissolved in isopropyl alcohol. The cells were lysed by sonication (20s on, 40s off, 20% amplitude, 9:30 minutes). The lysate was immediately centrifuged (18,000g, 30min, 4ºC) and the supernatant was collected and purified by nickel affinity chromatography followed by size-exclusion chromatography (SEC) (Superdex 75 10/300GL, GE Healthcare).

To cleave the His-tag, purified protein samples were incubated with 1:100 (w/w) of TEV Protease (New England Biolabs) at room temperature for 1hr and then 20ºC for two nights. The samples were purified by nickel purification by HisTrap (HisTrap HP 1mL, Cytiva) on an AKTA Pure. Cleaved proteins were incubated in 1mM of EDTA (Santa Cruz Biotechnology) for 3hr at 37ºC, filtered through a 2.5µm syringe filter, and subsequently buffer exchanged on SEC (Superdex 75 10/300GL, GE Healthcare) into 20mM MOPS 150mM NaCl, pH 7.4. All protein samples were characterized by commaise stain SDS–PAGE (BioRad) (**Figure S6**). Protein concentrations were determined by absorbance at 280 nm measured with a Nanodrop spectrophotometer (Thermo Fisher Scientific) using the predicted extinction coefficient and molecular weight from ProtParam Expasy.

Single-site alanine mutagenesis.

Single-site alanine mutations were introduced by PCR-based site-directed mutagenesis. Primers were designed to substitute each codon for the target residue with an alanine codon (GCT) (**Table S1B**). Primers were synthesized by Integrated DNA Technologies (IDT) and checked for melting temperature (Tm ~ 60ºC), GC content, and absence of strong secondary structure. Mutagenesis reactions were performed using Phusion High-Fidelity DNA Polymerase (New England Biolabs) following manufacturer instructions, with the plasmid DNA encoding the wildtype design as template. Following PCR amplification, the reaction is treated with Dpn1 restriction enzyme (New England Biolabs) for 1 hour at 37ºC. Digested PCR products were transformed into chemically competent E. coli DH5α cells. They were recovered in SOC medium and plated on LB agar supplemented with kanamycin. Colonies were picked and grown overnight, and plasmid DNA was purified using a miniprep kit.

All constructs were verified by Plasmidsaurus across the full coding region to confirm the presence of the desired alanine substitution and absence of unwanted secondary mutations. Verified plasmids were purified according to "Phase 1: Metalloprotein protein expression and purification" in Methods.

Circular dichroism and thermal denaturation.

CD spectra were measured on a JASCO CD spectrophotometer in a 1-mm-pathlength cuvette (Hellma). Protein samples were at ~0.2 mg $ml^{-1}$ in the 20 mM MOPS 150mM NaCl, (pH 7.4) buffer. Protein 1 was further prepared with additional 2mM TCEP (Tris(2-carboxyethyl)phosphine), GoldBio) reducing agent. To prepare protein with zinc, equimolar Zinc chloride (Sigma) was added to each sample and equilibrated for 1 hour. Melting temperature ranged from 20 to 95 °C and the absorption signal was monitored at a wavelength between 211 nm and 221 nm (**Figure S7**) in 1 °C increments per minute, with 5 s of equilibration time and 1 s of digital integration time. Wavelength scans (200–260 nm) were collected at 20 and 95 °C and again at 20 °C after fast refolding. To perform the EDTA experiment of protein 7, equimolar EDTA was added at 95ºC followed by an immediate rescan and then re-folding to 20ºC of the same sample. All measurements were baseline corrected.

Metal-binding affinity by MagFura-2 competition titration.

Competition assays using MagFura-2 were performed to determine the kinetic binding affinities of the zinc metal ion. The MagFura-2 was prepared as 1mM stocks in filtered water. The protein used was EDTA-cleaved and buffer exchanged into 20mM MOPS 150mM NaCl (pH 7.4). Protein 1 was further prepared with additional 1mM BME reducing agent. In a quartz-bottomed 96-well plate, 16 wells were prepared for each protein with 5µM of protein and 5µM MagFura-2, with an aliquot of $ZnCl_2$ (400µM stock solution) to reach a final concentration between 0 μM and 15μM. Each protein had 5 replicates of the above titration. The plate was placed in a plate reader (SpectraMax iD3) at 20ºC and equilibrated by shaking low intensity for 10 minutes prior to absorption read. The change in absorption ratio between 335nm and 365nm was used to measure the concentration of metal-bound MagFura-2. Absorption was read three times per wavelength and averaged. The titration curves were fit to a one-site binding model using the 1:1:1 competition fit in PyBindingCurve (*63*) (**Figure S8**).

Cobalt Substitution as a Spectroscopic Probe of Metal-Binding.

Cobalt and zinc are chemically similar ions: divalent cations with similar ionic radii and overlapping ligand preferences. However, cobalt is spectroscopically active when protein-bound due to its partially filled d-orbitals, enabling detection by UV-vis. We therefore used cobalt substitution as an additional probe to confirm metal binding.

To determine whether the designed proteins bound cobalt, we relied on spectroscopic readout of the ligand-to-metal charge-transfer absorbance. Protein used was EDTA-cleaved and buffer exchanged into 20mM MOPS, 150mM NaCl (pH 7.4). The holo protein was prepared by co-incubating with equimolar concentration $CoCl_2$ (Cobalt chloride(II) hexa hydride, Sigma Aldrich) prepared in protein buffer at room temperature for thirty minutes to ensure equilibration.

Spectroscopic measurements were performed using a UV-visible spectrophotometer (Genesys 180, ThermoFisher Scientific) with a 1cm pathlength quartz cuvette. Absorbance spectra of the proteins were collected between 450nm to 600nm with a 1mm step size. Control spectra of the buffer with cobalt alone was recorded under identical conditions and subtracted from all forthcoming spectra. Spectra were then measured for the apo and holo proteins to isolate the protein-associated Co(II) absorbance. Similar measurements were taken for the binding site point mutants. Binding was assessed by the presence of a d-d transition band in the visible region (**Figure S9**).

Determination of zinc content by ICP-MS.

All metalloproteins were EDTA-cleaved then buffer exchanged into 20mM MOPS, 150mM NaCl (pH 7.4) before ICP-MS. Protein 1 was further prepared with additional 1mM BME reducing agent immediately before ICP-MS preparation. These were the background controls in their apo form ("control"). To evaluate metal-binding, 100μL of 7.6μM protein was added to 400μL of 150μM $ZnCl_2$. The samples were left to equilibrate for 1 hour. Then each sample was concentrated down at 6000rpm on a table-top centrifuge to 100uL (the "concentrate"), leaving 400μL in solution (the "flow-through").

For each protein, three samples were given: (a) 100μL stock protein at 7.6uM without zinc (b) the "concentrate" and (c) the "flow-through". Samples were prepared in the SIGMA class 100 metal-free clean lab facility. Sample (a) was a control to ensure there were minimal to no residual free metals (**Table S3**). Concentrated high purity nitric acid and hydrogen peroxide (Fisher Scientific Optima grade) were used for sample digestion. Samples were heated and dried before dilution in 2% (v/v) nitric acid for introduction into the ICPMS instrument.

Analysis was performed on an Agilent 8900 ICP-MS attached to an Agilent SPS-4 autosampler. The sample introduction system includes a standard Scott double pass cooled spray chamber operated at 2ºC, a 2.5mm i.d. Agilent glass torch, and a 400 µl/min Micromist nebulizer. The ICP-MS was fitted with high sensitivity nickel cones and operated in $NH_3$ mode (4 mL/min) to reduce the presence of isobaric interferences from argides generated in the plasma. Samples were further diluted by 50% with a Sc internal standard teed into the sample introduction system; Sc was used to correct for instrumental drift over the course of the analysis. Quality control was facilitated using a well characterized in-house standard (KBL) which was also used to ensure accuracy of the concentration measurements. Samples were run in replicate (n=3) to determine the external reproducibility.

The Differential Factor is calculated as described in equation 1 and reported in **Extended Data Fig. 4**. The baseline 0.20 is the theoretical value if no zinc binds the protein, in which case the zinc is evenly distributed between the concentrate and the flow-through, leaving a 1:0 ratio. For a single-site metal-binding site, a differential factor of ~0.4 is expected for mono-nuclear binding.

$$(1)\ \text{Differential Factor} = \frac{\text{mass of Zinc in "the concentrate"}}{\text{mass of Zinc in "the flow-through"}}$$

Phase II: Metalloenzyme design and characterization

Computational design of metalloenzymes.

An initial set of 10,000 backbone structures was unconditionally generated by Protpardelle-1c between lengths of 200 and 300 residues to diversify the structural search space. Structures that lacked any beta-strand content or were >75% helical content as determined by DSSP were discarded. A sequence for all remaining backbones was designed using the ProteinMPNN to allow for structures to be further clustered by TM-score of 0.7 using Foldseek. To take advantage of a well-defined pocket, cluster representatives that resembled a TIM barrel fold were manually selected for further design. This resulted in a final 20 structures.

To restrain the explored design space, we took advantage of the barrel. For each of the 20 initial scaffolds, one sequence was generated for that backbone (described later) while residues that were in the barrel were manually identified. The identified residues were then used as variable residue positions to be further designed by dEVA while the rest of the sequence was fixed.

For this second round of design, the original Metal3D model was replaced with the newly trained Metal3D-Cat, described above. Since no official LigandMPNN training script was made public at time of use, we attempt to correct for the training dataset bias by applying an amino acid bias: "H:1.5,E:-0.5,D:-0.5". Cysteines were omitted from design. A successful designed sequence was considered if at least four of the designed sequences along Pareto front had a p(catalytic metal) > 0.85.

In an initial round, all the sequences for each candidate backbones were initially designed by one ProteinMPNN sequence (cysteines also omitted from design) followed by further design by dEVA. However, no designs were able to meet the Pareto front cutoff. A subsequent round used ensemble Caliby (*49*) to design 16 sequences (cysteines also omitted from design); the sequence with lowest energy was further design by dEVA with the above described above. Similarly, the designs were made omitting cysteines. The sequence along the Pareto front that maximized the p(catalytic metal) was chosen for further characterization. This resulted in three sequences: desA, desB, and desC.

Metalloenzyme protein expression and purification.

Purification was performed with Strep-Tactin® 4Flow® high-capacity resin, acquired from IBA LifeSciences. To perform the affinity chromatography with Strep-Tactin® resin, three separate buffers were initially prepared with Milli-Q water and filtered as follows: **W1 buffer** (100mM Tris-HCl, 150mM NaCl, 1mM EDTA, pH8.0), **W2 buffer** (100mM Tris-HCl, 150mM NaCl, pH 8.0), **E buffer** (40mM HEPES, 50mM NaCl, 2.5mM desthiobiotin, pH 8.0; the buffer was acquired from IBA-Lifesciences at 10x concentration and diluted in Milli-Q water), **Regeneration buffer** (100mM NaOH, pH 8.0). All buffers were clarified through a 2.5µm filter.

Genes encoding the designed protein sequence were synthesized and cloned into pET-24a(+) *Escherichia coli* plasmid expression vectors (Genscript), wherein the protein sequence contained a C-terminal Strep-tag separated by a "SA" spacer (**Table S5**). We later found that the sequence for desB contained a C-terminal tandem Strep-tag which was a result of unintended duplication during cloning. Plasmids were then transformed into chemically competent BL21(DE3) E. coli (Zymo Research). The cells were cultured in 2xYT medium at 37 °C until the optical density (OD) reached 0.6–0.8. Protein expression was then induced with 1 mM IPTG at 16 °C. After overnight expression, cells were centrifuged (4,000xg, 30min, 4ºC) and resuspended with W1 buffer and frozen at −80 °C until extraction and purification.

The cell pellet was thawed and supplemented with final concentration of 1mM PMSF dissolved in isopropyl alcohol. The cells were lysed by sonication (20s on, 40s off, 20% amplitude, 9:30 minutes). The lysate was immediately centrifuged (18,000g, 30min, 4ºC) and the supernatant was collected and purified by Strep-Tactin® chromatography followed by SEC (Superdex 75 10/300GL, GE Healthcare), described next.

Purification by Strep-Tactin® affinity chromatography was performed in 4ºC. Plastic 25mL gravity flow-through chromatography columns were prepared with Strep-Tactin® resin in a 50% resuspension (1 CV = 1.5mL of resin). The columns were equilibrated 2 times with at least 10 CV of W2 buffer. Once the columns were equilibrated, the lysate supernatant was gently added over the columns and allowed to completely flow-through. The columns were then washed with 6 CV of W2 buffer, and 1 time with 0.5 CV of E buffer. Next, the bound protein in each column was eluted into individual Eppendorf tubes with 5 individual additions of 0.5 CV of E buffer, saved for further purification. The columns were washed with 2 CV of E buffer, washed in water, and then regenerated with 10 CV of regeneration buffer. Between uses, the column was washed with MilliQ water and stored in 20% v/v ethanol and water to prevent bacterial growth, capped at both ends, and stored at 4ºC for future use. If the column was used after being stored in ethanol, it was promptly washed with Milli-Q before regenerating with Regeneration Buffer as previously described.

Individual fractions were filtered through a 2.5µm syringe filter and subsequently then purified by size-exclusion chromatography (SEC) at room temperature on an AKTA Pure with a Superdex 75 10/300GL****, GE Healthcare with a running buffer of 40mM HEPES, 50mM NaCl, pH 8.0. The major, resulting monomeric fraction was collected and immediately used for downstream experiments. All protein samples from this purification were characterized by commaise stain SDS–PAGE (BioRad) (**Figure S14, S18**) . Protein concentrations were determined by absorbance at 280 nm measured with a Nanodrop spectrophotometer (Thermo Fisher Scientific) using the predicted extinction coefficient and molecular weight. The further characterized design *desB* was additionally confirmed by liquid chromatography mass spectrometry (Methods described later).

**** *Note:* In the very initial purification of these designed proteins, the Superdex 75 10/300GL used was same as the one used in Phase I (and by others in our group). We believe this column had residual free phosphate, which we later found through X-ray absorption spectroscopy (described later) to have bound phosphate in high quantities in a Malachite Green Phosphate Assay (described later). Similarly, when we performed ICP-MS on the desB sample, we found residual nickel in the sample. Suspecting this is from residual free phosphate found in common sodium-phosphate buffers and contaminating nickel from NiNTA affinity column chromatography, we purchased a new Superdex 75 10/300GL that was never run with phosphate-containing buffers, to minimize phosphate contamination and potential inhibition. All kinetic characterization, including reported design screens described, for the reported metalloenzymes were purified with this new, phosphate-free, nickel-free Superdex 75.

Single-site alanine mutagenesis.

Single-site alanine mutations were introduced by a PCR based site-directed mutagenesis. Primers were designed to substitute each codon for the target residue with an alanine codon (GCT). Primers were synthesized by Integrated DNA Technologies (IDT) and checked for melting temperature (Tm ~ 60ºC), GC content, and absence of strong secondary structure (**Table S6**). Mutagenesis reactions were performed using Phusion High-Fidelity DNA Polymerase (New England Biolabs) following manufacturer instructions, with the plasmid DNA encoding the wildtype design as template. Following PCR amplification, the reaction is treated with Dpn1 restriction enzyme (New England Biolabs) for 1 hour at 37ºC. Digested PCR products were transformed into chemically competent E. coli DH5α cells. They were recovered in SOC medium and plated on LB agar supplemented with kanamycin. Colonies were picked and grown overnight, and plasmid DNA was purified using a miniprep kit.

All constructs were verified by Plasmidsaurus across the full coding region to confirm the presence of the desired alanine substitution and absence of unwanted secondary mutations. Verified plasmids were purified according to "Phase 2: Metalloenzyme protein expression and purification" in Methods.

Kinetic analyses.

Kinetic  experiments were done on a Tecan Infinite M200 Pro, unless otherwise noted, and fluorescence experiments were performed in a Greiner 96 well, F-bottom (chimney well), black, FluoTrac plate while absorbance experiments were performed in a Corning 384 well clear microplate, Each purified protein sample was diluted to 10x of their desired concentration in the reaction using the **reaction buffer** (40mM HEPES, 50mM NaCl, pH 8.0). Additionally, zinc sulfate (Sigma) dissolved in Milli-Q water was prepared and added to each of protein samples, to reach the final desired [zinc]:[enzyme] ratio (described in each aspect of kinetic screening). The samples containing protein and zinc sulfate were incubated

The fluorophores used to detect **phosphate monoesterase activity** are the fluorogenic monoester 4-methylumbelliferyl phosphate (4-MUP, Sigma) and 6,8-Difluoro-4-methylumbelliferyl phosphate (DiF-MUP, Sigma). The fluorophores used to detect **phosphate diesterase activity** are fluorogenic methyl 4-nitrophenyl phosphate (me-pNPP, synthesized by and acquired from the Herschlag Lab from previous work (*64*)) and bis-(4-nitrophenyl) phosphate (bis-pNPP, Sigma). To generate the standard curve, the fluorescent product of the phosphomonoesters was monitored by a plate reader using an excitation/emission wavelength of 360nm and 445, respectively.

The stock solutions of 4-MUP and 4-MU were made by dissolving each of the solids in 100% DMSO to a final concentration of 10mM. The stock solutions of DiF-MUP and DiF-MU were made by dissolving each of the solids in 5% DMSO to a final stock concentration of 10mM. The me-pNPP was acquired from Herschlag Lab as a pre-synthesized stock, where the solids were dissolved in MilliQ water**.** The stock solution of bis-pNPP was made by dissolving the solid in MilliQ water.

Data regarding kinetics can be found in the main text and **Figures S16-19**.

<u>Initial design screen.</u>

Each purified protein sample was diluted to 10µM protein stock using the reaction buffer. For the initial screen, 1µM of protein was supplemented with the expected zinc molar equivalence (1x for desA and desC, 2x for desB) in the plate. Briefly, this means adding 84µL of reaction buffer, followed by 10µL of the 10µM protein stock, and 2µL of the appropriate zinc sulfate stock into each well. The plate containing protein and zinc was incubated for 15 minutes in the plate reader to equilibrate. The final protein sample concentration is 1µM.

Then 4µL of an 800µM stock of the 4-MUP fluorogenic ester was added to each sample well (including background traces in water, buffer, and buffer+zinc), resulting in a final substrate concentration of 32µM. Reads were measured every 15 seconds for the first 15 minutes of the reaction.

<u>Kinetic screen under variable pH.</u>

To characterize the relative pH profile of desB, initial reaction rates (in RFU/min) were recorded. All measurements were collected on a SpectraMax iD3 microplate reader in a Thermo Scientific 96-well black flat bottom plate. Default settings, opaque plate, and medium gain were used for this screen.

To assess the effect of pH on desB activity, nine different conditions were prepared in advance. The conditions are as follows:

40mM HEPES, 50mM NaCl, pH 7.0
40mM HEPES, 50mM NaCl, pH 7.5
40mM HEPES, 50mM NaCl, pH 8.0
40mM CHES, 50mM NaCl, pH 8.5
40mM CHES, 50mM NaCl, pH 9.0
40mM CHES, 50mM NaCl, pH 9.5
40mM CHAPS, 50mM NaCl, pH 10.0
40mM CHAPS, 50mM NaCl, pH 10.5
40mM CHAPS, 50mM NaCl, pH 11.0

where HEPES, CHES, and CHAPS were all acquired from Sigma. Purified protein desB was diluted to a 10µM stock in the reaction buffer. For each well, 84µL of the varying pH buffer condition was added to an individual well, along with 10µL of 10µM stock (final concentration of 1μM enzyme), and 2μL of a 96μM zinc sulfate stock. Prior to the addition of substrate, the plate was incubated in the plate reader for 15 minutes to allow for equilibration. 4μL of an 800μM stock of the fluorogenic ester was added to each sample well, resulting in a final substrate concentration of 32μM.

Reads were measured every 10 seconds for the first 15 minutes of the reaction. The resulting fluorescence curves were manually set to start at y=0 relative fluorescence units such that a linear regression fit $y = mx + b$, where the y-intercept b=0, therefore only needing to fit the slope. The calculated slopes for the initial velocities were reported in units RFU/min.

Kinetic screen under variable temperature.

To characterize the relative pH profile of desB, initial reaction rates (in RFU/min) were recorded. All measurements were collected on a SpectraMax iD3 microplate reader in a Thermo Scientific 96-well black flat bottom plate. Default settings, opaque plate, and medium gain were used for this screen.

To assess the effect of temperature on desB activity, six different wells were prepared as described in the "Initial design screen". Prior to the addition of substrate, the plate was incubated in the plate reader for 15 minutes at each respective temperature to allow for equilibration. Then for one well at each temperature, 4μL of an 800μM stock of the fluorogenic ester was added, resulting in a final substrate concentration of 32μM.
Reads were measured every 10 seconds for the first 15 minutes of the reaction. The resulting fluorescence curves were manually set to start at y=0 relative fluorescence units such that a linear regression fit $y = mx + b$, where the y-intercept b=0, therefore only needing to fit the slope. The calculated slopes for the initial velocities were reported in units RFU/min.

Kinetic screen under variable ionic strength.

To characterize the relative ionic strength profile of desB, initial reaction rates (in RFU/min) were recorded. All measurements were collected on a SpectraMax iD3 microplate reader in a Thermo Scientific 96-well black flat bottom plate. Default settings, opaque plate, and medium gain were used for the screen.

To assess the effect of ionic strength on desB activity, five different conditions were prepared in advance. The conditions are as follows:

40mM HEPES, 50mM NaCl, pH 8.0
40mM HEPES, 150mM NaCl, pH 8.0
40mM HEPES, 500mM NaCl, pH 8.0
40mM HEPES, 1M NaCl, pH 8.0
40mM HEPES, 2M NaCl, pH 8.0

Reactions were assessed under two different substrate regimes: saturating conditions where $[E] << [S]$ and at $[E] = [S]$. Purified protein desB was diluted to a 10μM stock in the reaction buffer. For each well, 84μL of the varying ionic strength buffer condition was added to an individual well, along with 10µL of 10µM protein (final concentration of 1µM enzyme), and 2µL of a 96µM zinc sulfate stock. Prior to the addition of substrate, the plate was incubated in the plate reader for 15 minutes to allow for equilibration. Then to assess [E] << [S] reaction conditions, 4µL of an 800µM stock of the fluorogenic ester was added to each sample well, and to assess [E] = [S], 4µL of a 25µM stock of the fluorogenic ester was added to each sample well.

Reads were measured every 10 seconds for the first 15 minutes of the reaction. The resulting fluorescence curves were manually set to start at y=0 relative fluorescence units such that a linear regression fit $y = mx + b$, where the y-intercept $b=0$, therefore only needing to fit the slope. The calculated slopes for the initial velocities were reported in units RFU/min.

Phosphomonoesterase: 4-MU standard curve.

One set of serial dilutions of the 4-MU (stock is at 80µM to 625nM in two-fold dilutions, final reaction concentrations of 3.2µM to 25nM) was made into anhydrous DMSO to construct a fluorescence calibration curve. Then 4µL of each 4-MU dilution was added to 96µL of reaction buffer; the final concentration of DMSO in each well is 4% v/v. Each point was measured at 25ºC in a 96-well plate was measured in triplicate and averaged over 5 minutes of measurements. Instrument was set to 60 gain, 25 flashes, and 20µs integration time. The calibration curve was fit with a linear regression equation to convert relative fluorescence units into the product concentration of 4-MU.

Phosphomonoesterase: 4-MUP kinetics.

Serial dilutions of the 4-MUP substrate (stock at 80µM to 625nM in two-fold dilutions, final reaction concentrations of 3.2µM to 25nM) were made in anhydrous DMSO.
The desB protein sample was selected for further kinetic characterization with 4-MUP. Three biological replicates were purified on three separate days using the protein purification process described above. Between preparation days, the already purified protein samples were stored at 4ºC. The three biological replicates were diluted to 250nM concentration, for a final enzyme concentration of 25nM. The zinc sulfate was prepared to 500nM, for a final concentration of 4µM, in other words [E]:[Zn] = 1:2.

In a 96 well plate, the protein sample and zinc were added and equilibrated – 84µL of reaction buffer, 10µL of protein stock, and 2µL of the stock zinc sulfate. In separate rows, background hydrolysis reactions were carried out without any enzyme at 25ºC in three separate conditions: in reaction buffer, in reaction buffer and zinc, and in MilliQ water. The protein sample was equilibrated for 15 minutes, followed by immediate addition of 4µL of each dilution of 4-MUP. The plate was sealed and the fluorescence progress of each biological replicate was monitored overnight, with 1-minute reads. Instrument was set to 60 gain, 50 flashes, and 50 integration time.

Phosphomonoesterase: DiF-MU standard curve.

One set of serial dilutions of the DiF-MU was used to construct a fluorescence calibration curve. Then 4µL of each DiF-MU dilution was added to 96µL of reaction buffer. Each point was measured at 25ºC in a 96-well plate was measured in triplicate and averaged over 5 minutes of measurements. The instrument was set to a gain of 60, 50 flashes, and 50µs integration time. The calibration curve was fit with a linear regression equation to convert relative fluorescence units into the product concentration of DiF-MU.

<u>Phosphomonoesterase: DiF-MUP kinetics.</u>

Protein and substrate were prepared analogously to the 4-MUP kinetics. Three biological replicates were diluted to 250nM concentration, for a final enzyme concentration of 25nM. The zinc sulfate was prepared to 500nM, for a final concentration of 4µM, in other words [E]:[Zn] = 1:2. The plate was sealed and the fluorescence progress of each biological replicate was monitored with 1-minute reads. The instrument was set to a gain of 60, 50 flashes, and 50µs integration time.

<u>Phosphomonoesterase: desB mutants.</u>

Thirteen mutants of desB were synthesized, purified, and prepared according to methods described above. For each construct, a kinetic screen was performed analogous to the initial screen. In brief, 1µM of the protein was equilibrated with 2µM of zinc sulfate. This means adding 84µL of reaction buffer, followed by 10µL of the 10µM protein stock, and 2µL of the zinc sulfate stock into each well. The plate containing protein and zinc was incubated for 30 minutes in the plate reader to equilibrate. The final protein sample concentration is 1µM. Then 4-MUP fluorogenic ester was added to each sample well (including background traces in water, buffer, and buffer+zinc), resulting in a final substrate concentration of 1µM of the ester. Reads were measured every 15 seconds for the first 15 minutes of the reaction. This screen was performed on the SpectraMax plate reader. We note there was significantly less background in this screen, and we attribute this to the significantly lower 4-MUP concentration than in the initial design screen.

<u>Phosphodiesterase: me-pNPP kinetics.</u>

Serial dilutions of the me-pNPP substrate (final reaction concentrations of 80µM to 5µM in two-fold dilutions) were made in MilliQ water. For each sample, two biological replicates were diluted to a stock concentration 10µM concentration, with a final enzyme concentration of 1µM. The zinc sulfate was prepared to 100µM, for a final concentration of 2µM, in other words [E]:[Zn] = 1:2. The plate was sealed and the 400nm absorbance progress of each biological replicate was monitored with 1-minute reads. The instrument was set to 50 flashes.

<u>Phosphodiesterase: bis-pNPP kinetics.</u>

Serial dilutions of the bis-pNPP substrate (final reaction concentrations of 100µM to 6.25µM in two-fold dilutions) were made in MilliQ water. For each sample, two biological replicates were diluted to a stock concentration 10µM concentration, with a final enzyme concentration of 1µM. The zinc sulfate was prepared to 100µM, for a final concentration of 2µM, in other words [E]:[Zn] = 1:2. The plate was sealed and the 400nm absorbance progress of each biological replicate was monitored overnight, with 1-minute reads. Instrument was set to 50 flashes.

Michaelis-Menten kinetics.

The resulting curves were analyzed by first subtracting the background reaction at each substrate concentration point for the matching concentration in the reaction buffer. For 4-MUP and DiF-MUP, the resulting fluorescence curves were manually set to start at y=0 relative fluorescence units such that a linear regression fit y = mx + b, where the y-intercept b=0, therefore only needing to fit the slope. The calculated slopes for the initial velocities at each substrate concentration point were transformed to be expressed in terms of [4MU]/[E] using the calibration curve and known enzyme concentration.

The steady-state parameters $k_{cat}$ and $K_M$ were determined for enzyme hydrolysis by non-linear regression fitting initial data points to the Michaelis-Menten equation

$$\frac{v_o}{[E]} = \frac{k_{cat}[S]}{(K_M+[S])},$$

where $v_o$ denotes the initial velocity, [E] denotes the enzyme concentration, and [S] denotes the substrate concentration. Standard deviations were calculated from the standard error of the fit consider each biological triplicate value as an individual data point.

To determine the uncatalyzed reaction rate ($k_{uncat}$), the rate constant was determined by fitting all data points to the equation of $v_0 = k_{uncat}[S]$ using linear regression.

Determination of $k_{cat}/K_M$ from $k_{obs}$

The resulting curves were analyzed by first subtracting the background reaction at each substrate concentration point for the matching concentration in the reaction buffer. Initial rates for me-pNPP and bis-pNPP were measured under substrate concentrations well below the Michaelis constant ($[S] << K_M$), such that the reaction follows a pseudo-first-order kinetics with respect to the substrate. Under these conditions, we use the simplified equation

$$v_o = \frac{k_{cat}[E][S]}{K_M}$$

The molar extinction coefficient of product p-nitrophenolate is approximately 16,000 $M^{-1}$ $cm^{-1}$. Absorbance values were corrected for effective path length in the microplate format. For each condition, product formation was fit over the initial linear time window to obtain the initial rate $v_0$. Rates were normalized by the enzyme concentration to extract $k_{obs}$.

Under low-substrate conditions, linearly depends on substrate concentration; therefore, the $k_{cat}/K_M$ was obtained as the slope of a linear fit. Linear regression was performed using least-squares fit and uncertainties in $k_{cat}/K_M$ were estimated from the standard error of the slope.

Determining Apparent Inhibition Constants ($K_{i,app}$).

To determine the appear constants of desB, a dilution series of two inhibitors (inorganic phosphate and sodium tungstate, Sigma) were prepared to a final concentration of 25nM and assayed against 250nM of 4-MUP. Fluorescence was monitored over time as described above, and initial linear rates (RFU/s) were extracted. A background fluorescence rate was measured in the absence of enzyme and subtracted from each experimental rate.

Background-corrected rates were plotted as a function of inhibitor concentration and fit to a hyperbolic decay model:

$$v = \frac{m_1}{m_2 + [I]}$$

where v is the observed rate, [I] is the inhibitor, and $m_2$ corresponds to the apparent inhibition constant ($K_{i,app}$).

Attempting to prepare apo-desB by zinc chelation.

Multiple attempts at chelating zinc out of *desB* were taken. In brief, 10mM of EDTA was added to the purified protein sample and left to equilibrate for 1 hour. Following equilibration, the sample was filtered in a 2.5µm syringe filter and subsequently buffer exchanged by size-exclusion chromatography to remove any residual EDTA prior to analyses. Protein concentration was determined by absorbance at 280 nm measured with a Nanodrop spectrophotometer using the predicted extinction coefficient and molecular weight.

Two liters of culture were purified and chelated, which eventually yielded high enough concentration for biophysical characterization (0.2mg/ml at 200µL). Further characterization on CD and temperature melt for the EDTA-treated sample was performed. However, we note that a significant amount of protein was lost during chelation, suggesting tight-binding affinity of the zinc site(s) and resulting spectra indicate significant loss in stability and secondary structure.

Circular dichroism and thermal denaturation.

A circular dichroism (CD) spectrum for each designed metalloenzyme was measured on a JASCO CD spectrophotometer in a 1-mm-pathlength cuvette (Hellma). Protein samples were at ~0.2 mg $ml^{-1}$ in PBS buffer (137mM NaCl, 2.7mM KCl, 10mM $Na_2HPO_4$, 1.8mM $KH_2PO_4$, pH 7.4). Melting temperature ranged from 20 to 90°C and the temperature was increased in 1 °C increments per minute, with 5 s of equilibration time and 1 s of digital integration time. Wavelength scans (200–260 nm) were equilibrated for 2 minutes at every 10ºC interval, and three accumulations were taken per spectra.

Preparation of zinc-supplemented sample for ICP-MS and XAS.

To prepare a zinc-supplemented sample, 2L of desB culture were purified following Phase II purification protocol. Subsequently, all the protein sample was combined and supplemented with 50x zinc sulfate (Sigma). The sample is left to equilibrate for 20 minutes followed by concentration using a Thermo Scientific Pierce Protein Concentrator (10kDa). Prior to concentrating, the concentrator was washed 2x with reaction buffer. The sample was subsequently exchanged on SEC to remove excess zinc.

Determination of zinc content by ICP-MS.

Rather than performing zinc chelation, we sought to determine zinc content via ICP-MS in an alternative method than that described in Phase I. Two samples were prepared. Sample one, referred to as "desB (purified)" is the final protein stock purified after size-exclusion chromatography. Sample two, referred to as "desB (zinc)" is the zinc-supplemented sample, described above. We note that these proteins were purified before switching to the brand new Superdex 75 (described as a note in Methods of Phase II purification). An aliquot of the reaction buffer was provided for further reference with ICP-MS to determine trace metals in the buffer.

The desB (purified) sample was concentrated to ~10mg/ml and 7µL was provided to the facility, leading to a final protein concentration of 422µM. We can then calculate the expected [E]:[Zn] concentration and convert to nanograms. For desB (purified), we expect ~193ng of detected zinc for 1:1 stoichiometry and ~386ng of detected zinc for 1:2 stoichiometry.

The desB (zinc) sample was concentrated to ~12mg/ml and 8.5µL was provided to the facility, leading to a final protein concentration of 506µM. We can then calculate the expected [enzyme]:[zinc] concentration and convert to nanograms. For desB (zinc), we expect ~281ng of detected zinc for 1:2 stoichiometry and ~562ng of detected zinc for 1:2 stoichiometry.

Concentrated high purity nitric acid and hydrogen peroxide (Fisher Scientific Optima grade) was used for sample digestion. Samples were heated and dried before dilution in 2% (v/v) nitric acid for introduction into the ICPMS instrument. Analysis was performed on an Agilent 8900 ICP-MS attached to an Agilent SPS-4 autosampler. The sample introduction system includes a standard Scott double pass cooled spray chamber operated at 2º C, a 2.5mm i.d. Agilent glass torch, and a 400 µl/min Micromist nebulizer. The ICP-MS was fitted with high sensitivity nickel cones and operated in $NH_3$ mode (4 mL/min) to reduce the presence of isobaric interferences from argides generated in the plasma. Samples were further diluted by 50% with a Sc internal standard teed into the sample introduction system; Sc was used to correct for instrumental drift over the course of the analysis. Quality control was facilitated using a well characterized in-house standard (KBL) which was also used to ensure accuracy of the concentration measurements. Samples were run in replicate (n=3) to determine the external reproducibility (**Table S7**).

We note the trace amounts of nickel in the purified sample is hypothesized to come from residual nickel on the SEC Superdex 75 column, and note the elevated amount of copper and iron, which we hypothesize is potential contamination in the zinc sulfate powder (Sigma) from leaching in industrial production as the zinc sulfate used was not of high purity or mass spec grade.

Quantifying free orthophosphate in native and denatured desB.

The presence of free orthophosphate ($PO_4^{-3}$) present in the zinc-supplemented desB sample was quantified using a malachite green colorimetric assay according to the manufacturer's protocol (Sigma-Aldrich MAK307). Samples were analyzed in a 96 well-late in a clear Greiner Bio-One CELLSTAR flat bottom plate. Absorbance spectra were measured from 600nm to 660nm using a SpectraMax iD3 plate reader. Reactions contained protein at a final concentration of 4μM. Two conditions were tested. For the native condition, the desB (zinc) sample (described above) was analyzed intact. For the denatured condition, the same desB (zinc) sample was acid-denatured by addition of 2μL of 1M HCl followed by heating at 95ºC for 5 minutes prior to phosphate measurement. After treatment, 80μL of sample was combined with 20μL of malachite green working reagent, incubated for 30min at room temperature for color development. Phosphate concentration was determined from a phosphate standard curve prepared in parallel with the protein samples following blank subtraction (**Figure S20**).

Liquid-chromatography mass spectrometry (LC-MS).

For LC-MS characterization, 45μM of final, purified desB sample (one of the biological replicates used in kinetics) was provided to the Stanford Mass Spectroscopy Facility. The sample was analyzed by electrospray ionization mass spectrometry (ESI-MS) using a Waters Acquity UPLC system coupled to a Thermo Exploris 240 BioPharma Orbitrap mass spectrometer. It was injected as received and separated on a BioResolve RP mAb Polyphenyl column (450 Å, 2.7 μm, 100 × 2.1 mm; Waters) maintained at 50 °C, with a flow rate of 0.2 mL/min and an injection volume of 3 μL. Mobile phases consisted of solvent A (0.1% formic acid in water) and solvent B (0.1% formic acid in acetonitrile), and separation was achieved using the following gradient: 0–2 min, 95% A / 5% B; 3 min, 85% A / 15% B; 8.5 min, 35% A / 65% B; 10–12 min, 5% A / 95% B; 12.5 min, 95% A / 5% B; 13 min, 5% A / 95% B; 13.5–20 min, 95% A / 5% B. Mass spectra were acquired in full scan mode over an m/z range of 500–4000 at a resolution of 120,000, and data were deconvoluted using Byos Intact software (Protein Metrics, version v5.9.121-g8dcdb07d19 x64) to obtain intact mass distributions (**Figure S15**).

Structural determination methods.

X-ray crystallography.

desDEH (#2): The purified protein was concentrated to around 12 mg/mL went through screening of around 400 crystallization conditions. Additionally, drops were also setup with varying the protein concentration. The best crystals were obtained from SG1 screen well A2 (2M Ammonium Sulfate). Crystals were grown by sitting drops at 16°C using a 1:0.95 ratio of protein to well solution and took around 3 weeks to grow. Crystals were transferred to a well solution supplemented with 25% ethylene glycol and cryocooled in liquid nitrogen. Diffraction data was collected at the SSRL BL12-2 beamline using Dectris PILATUS EIGER 2XE 16M PAD detector. The crystals belonged to space group $P4_12_12$ with dimensions a=97.53Å, 97.53Å, 73.45Å, α=90º, β=90º, γ=90º. There were two monomers in the asymmetric unit.

All data were processed with AUTOPROC. The structure was solved by molecular replacement using MOLREP (*65*) with an *in silico* protein model as the search model. The best solution gave an R-factor of 51%. Multiple rounds of manual model building using coot and model building using the buccaneer program were needed to successfully complete the model. The structure was refined by using Refmac (*66*) and Phenix (*67*) and manually fitted using the Coot (*68*) program. The details of data collection and refinement are given in **Supplementary Table 10A**.

desE2D (#7): The purified protein was concentrated to around 14 mg/mL. The crystallization trials with around 800 conditions using protein supplemented with different amounts of Zinc. The best crystals were obtained from a pH selective screen developed at SSRL well A8 (0.15M Calcium Acetate, 0.25 M Citrate (pH 5.8), 0.25M HEPES (pH 7.5), 025 M ADA (pH 6.5), 0.25 M Tris (pH 8.0), 25% PEG 2KMME) diffracted to 1.55Å. Diffraction data was collected at the SSRL BL12-2 beamline using Dectris PILATUS EIGER 2XE 16M PAD detector. The crystals belonged to space group $P2_12_12_1$ with dimensions a=37.88Å, 56.40Å, 92.92Å, α=90º, β=90º, γ=90º.

All data were processed with AUTOPROC. The structure was solved by molecular replacement using MOLREP & Phaser (*69*) with an *in silico* protein model as the search model. The structure solution attempts were first tried with 2Å data collected from a crystal from a different crystallization that belonged to a different space group. This helped build some of the models correctly. The availability of higher resolution data enabled better model building and tracing of the model. Few model building attempts using Buccaneer (*70*) and manual building reduced the Rfree to around 33%. The structure was further refined by using Refmac and Phenix and manually fitted using the Coot program. The details of data collection and refinement are given in **Supplementary Table 10A**.

desB: The purified protein was concentrated to around 38 mg/mL went through screening of around 800 crystallization conditions at 16°C. Drops were also setup with varying the protein-well drop ratio. All trials were unsuccessful. Then, crystallization screens were setup at 4°C and the crystals were obtained from few crystallization conditions of SSRL's Buffer Selective screen. The best crystals were from well F3 using HAT buffer (15% PEG20K, 0.15 M Sodium Malonate (pH 7.0), 0.033 M HEPES (pH 7.5), 0.033 M ADA (pH 6.5), and 0.033 M Tris-HCl (pH 8.0)). Crystals were grown by sitting drops at 4°C using a 1:1 ratio of protein to well solution and took around 4 weeks to grow. Crystals were transferred to 100 mM $ZnSO_4$ soaking solution (Soaking solution is 100 mM $ZnSO_4$ in well solution containing 2% higher 20K and 20% glycerol) for 90 Sec prior to cryocooling cryocooled in liquid nitrogen. Diffraction data was collected at the SSRL BL12-2 beamline using Dectris PILATUS EIGER 2XE 16M PAD detector. The crystals were thin plates and the diffraction along the thin edge had higher anisotropy in diffraction. Therefore, several data sets were collected, and the best diffraction data set was used for the structure solution. The crystals belonged to space group $P2_1$ with dimensions a=40.44Å, 78.84Å, 73.72Å, α=90°, β=92.4°, γ=90°. There were 2 monomers in the asymmetric unit.

All data were processed with AUTOPROC. The structure was solved by molecular replacement using MOLREP with an in-silico protein model as the search models. The best solution gave an R-factor of 50%. Multiple rounds of manual model building using coot and model building using the buccaneer program were needed to successfully complete the model. The structure was refined by using Refmac and Phenix and manually fitted using the Coot program. The details of data collection and refinement are given in **Supplementary Table 10A**.

desHE2 (#8): The purified protein was concentrated to around 20 mg/mL. The crystallization trials with around 600 conditions with 16 mg/mL and 12 mg/mL protein concentrations. The crystallization behavior of the protein was very strange and required number of crystallization attempts. Due to variability in diffraction, few crystals were screened to collect good data. The best crystals were obtained from SG1 screen well B7 (0.2M Sodium formate and 20% PEG 3350) diffracted to 1.42Å. The crystals belonged to space group P212121 with dimensions a=38.11Å, 75.29Å, 88.18Å, α=90°, β=90°, γ=90°. There was one monomer in the asymmetric unit. Diffraction data was collected at the SSRL BL12-2 beamline using Dectris PILATUS EIGER 2XE 16M PAD detector.

All data were processed with XDS (*71*). The structure was solved by molecular replacement using MOLREP with an *in silico* protein model as the search models. Few rounds of manual building and refinement were needed to successfully complete the structure. The structure was further refined by using Refmac and Phenix and manually fitted using the Coot program. The details of data collection and refinement are given in **Supplementary Table 10B**.

X-ray absorption spectroscopy (XAS).

Additional preparation of the metalloproteins.

Proteins desH2C2 (#1), desDEH (#2), and desE2D (#7) were prepared as previously described in Phase I and exchanged into a final buffer of 20mM MOPS, 150mM NaCl, 20% (w/v) glycerol (pH 7.4) to a final concentration of ~1mM. $ZnCl_2$ was added to achieve a concentration equal to 0.8 molar equivalents relative to protein. desH2C2 was supplemented with reducing agent as previously described. The samples were concentrated in a Thermo Fisher Pierce 3kDa Concentrator.

Additional preparation of metalloenzyme desB.

The sample used for desB (zinc) was prepared according to methods described in Phase II (Preparation of zinc-supplemented sample for ICP-MS and XAS). The sample was concentrated in a Thermo Fisher Pierce 10kDa Concentrator.

Performing XAS.

Purified protein samples were loaded into cuvettes sealed with 25 µm-thick Kapton tape, flash frozen, and stored in liquid nitrogen until measurement. XAS spectra were measured at the Stanford Synchrotron Radiation Lightsource on beamline 9-3 and 7-3. A Si(220) double crystal monochromator, oriented at Φ=0, was used for energy selection at Zn K-edge. The harmonic rejection was provided by a rhodium-coated mirror on BL 9-3 and by detuning the monochromator for 40% at BL 7-3. The scattering signal was reduced using a Cu filter and Soller slits. Samples were maintained at 10 K in He atmosphere during data collection using an Oxford liquid helium cryostat. Spectra were collected in fluorescence mode using a multi-element Canberra Ge detector, and a Zn foil reference spectrum was collected in transmission mode simultaneously for energy calibration with each scan. Four spots on each sample were measured (two to six scans each) to minimize beam damage, and no beam damage (such as reduction in whiteline intensity) was observed during data collection.

For data processing, channels with large background signals from ice diffraction were removed. The rest of the channels were then averaged in ATHENA (*72*). Data were normalized and calibrated by setting the edge energy (first maximum in first derivative) of the Zn foil to 9659 eV. The normalized spectra were imported into PySpline (*73*) for post-edge background subtraction. EXAFS was extracted by setting E0 to 9670 eV and fitting the background using a 3-region spline (with polynomial orders 2, 3, and 3). Fitting of $k^3$-weighted EXAFS was performed using EXAFSPAK (*74*), and the theoretical phase and amplitude of scattering paths were generated using FEFF7 (*75*). Starting geometry of the models were adopted from structures predicted by the protein design algorithm.

Greater uncertainty is associated with the Zn-Zn distance of desB+Pi, as EXAFS is inherently less sensitive to more distant scatters. Models with Zn-Zn distances ranging from 4.0 to 4.3Å yielded comparable fit quality. Representative multiple scattering paths are included in the model, and the number of paths used for fitting the desB+Pi and apo desB samples are kept the same. The number of independent points in the spectra is given by $2\times\Delta k\times\Delta R/\pi+2 = 2\times(11.2-2)\times(4-1)/3.14+2\approx20$, and 15 parameters were used for the fitting. All final fits are reported in **Table S4.**

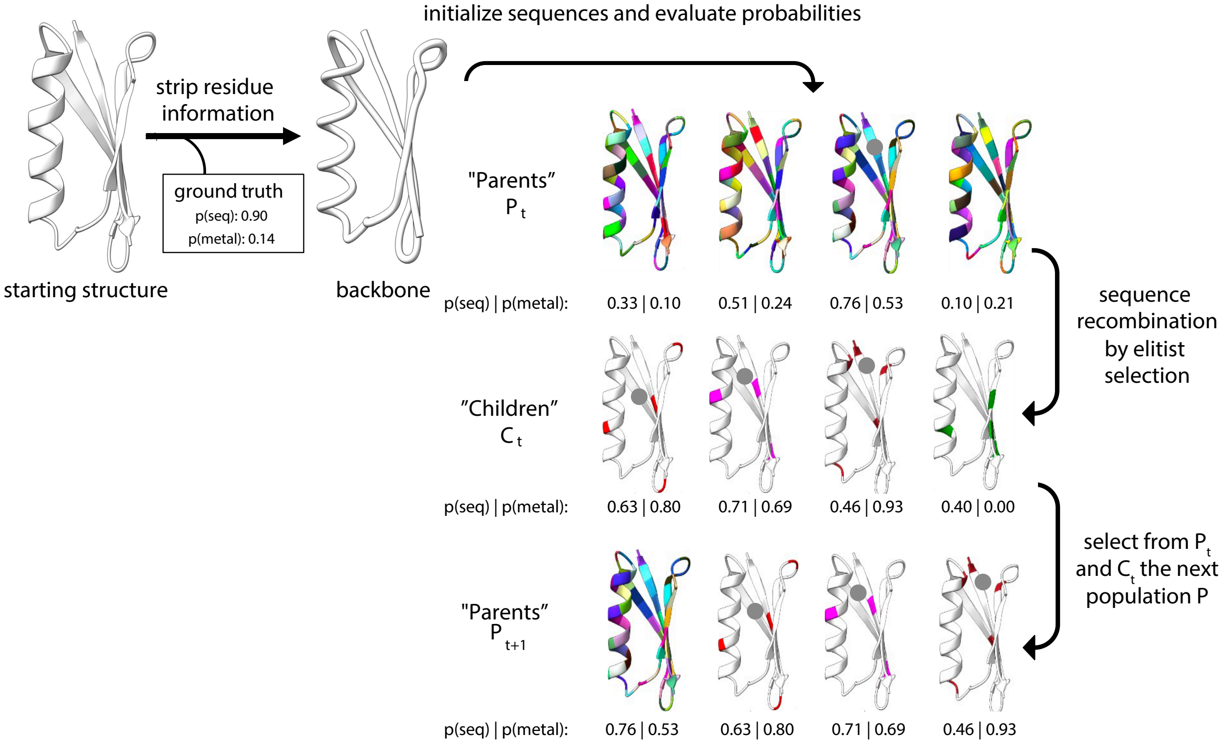


**Figure S1.** Detailed outline of the dEVA algorithm with an example of one generation of designs following a genetic algorithm. Mutations are made as residue-level changes based on LigandMPNN while the predicted metal position is predicted by Metal3D at each iteration.

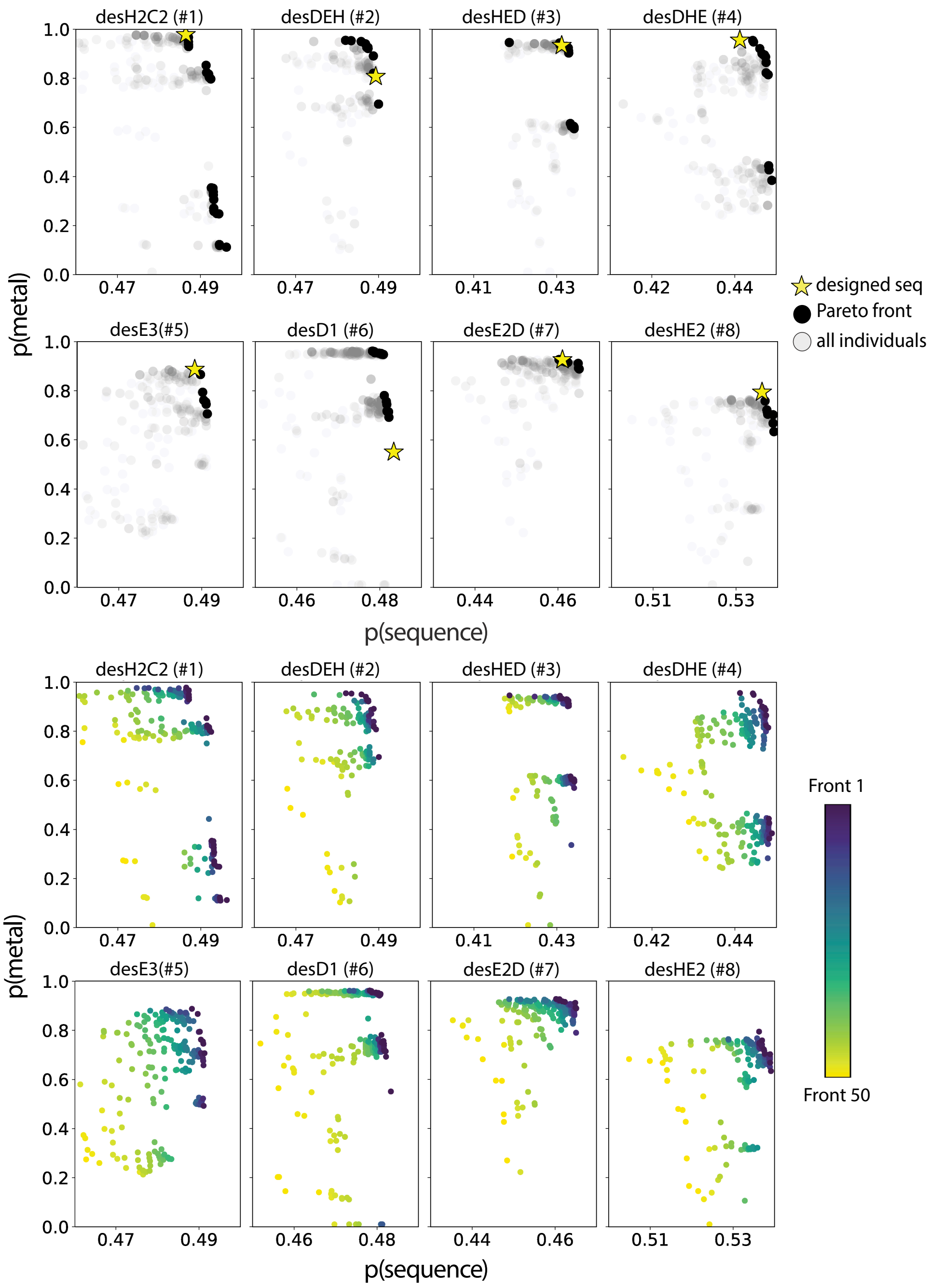


**Figure S2.** Computational design using dEVA. (a) All fifty generations explored. Final Pareto front population is shown in black, the experimentally tested sequence is indicated by a yellow star, and all other sampled sequences are shown in light grey. (b) Iterative progression of each population across generations. dEVA efficiently converges towards the optimal trade-off between sequence likelihood and metal-binding probability, with successive fronts moving closer to the knee point of the Pareto front.

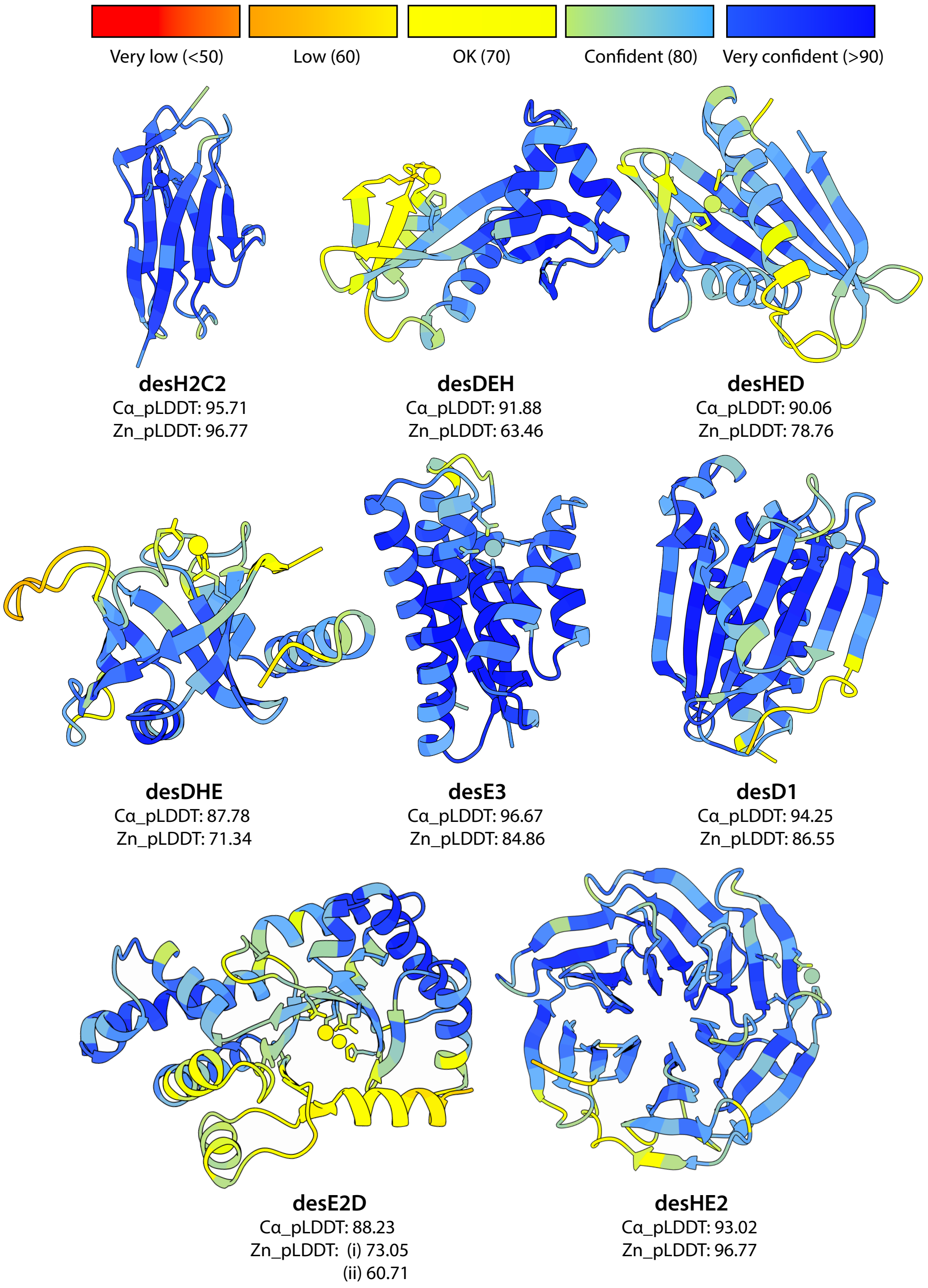


**Figure S3**. Predicted structures of the metalloprotein designs colored by AlphaFold-3 pLDDT. All designs exhibit varying backbone and zinc confidence indicated by Ca_pLDDT and Zn_pLDDT, respectively.

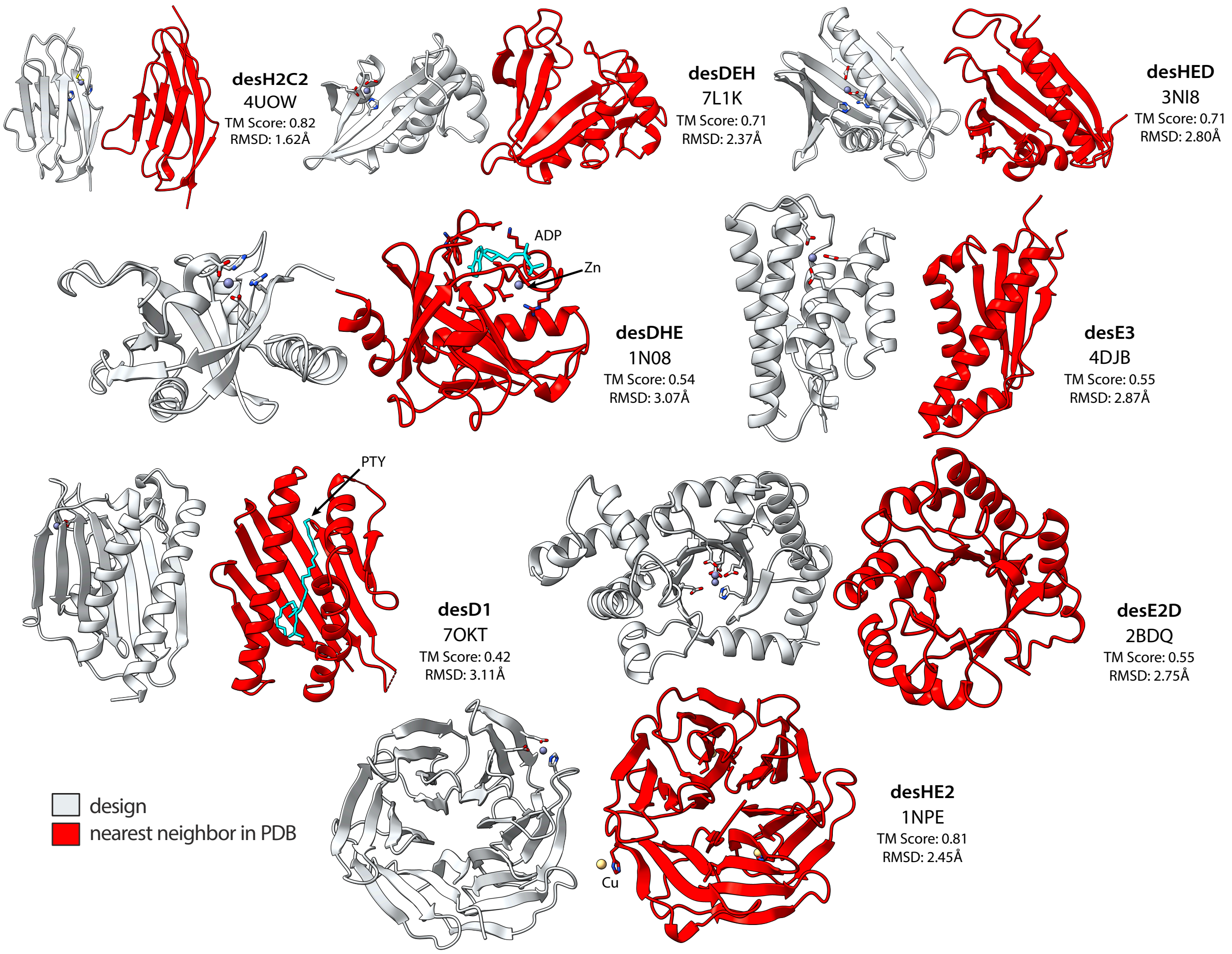


**Figure S4.** Comparison of nearest neighbor structure in the PDB (red) and designed structures (white). RMSD is evaluated using *cealign* and TM-score is evaluated using TM-align.

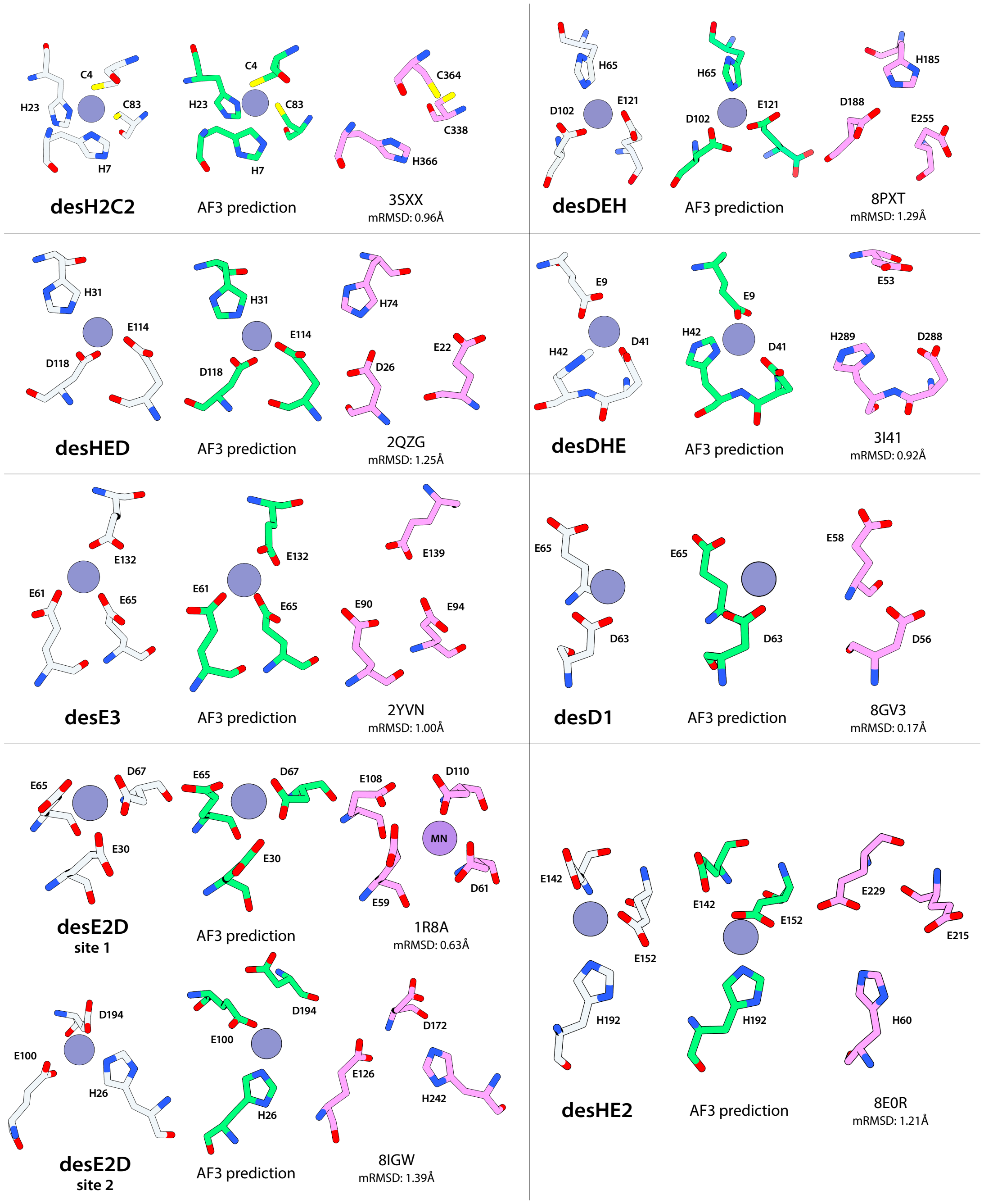


**Figure S5**. Comparison of designed metal-binding site (grey) to AlphaFold-3 predicted metal-binding site (green) and nearest neighbor from RCSB search (pink). Overall, first-shell coordinating residues are predicted by AlphaFold-3.

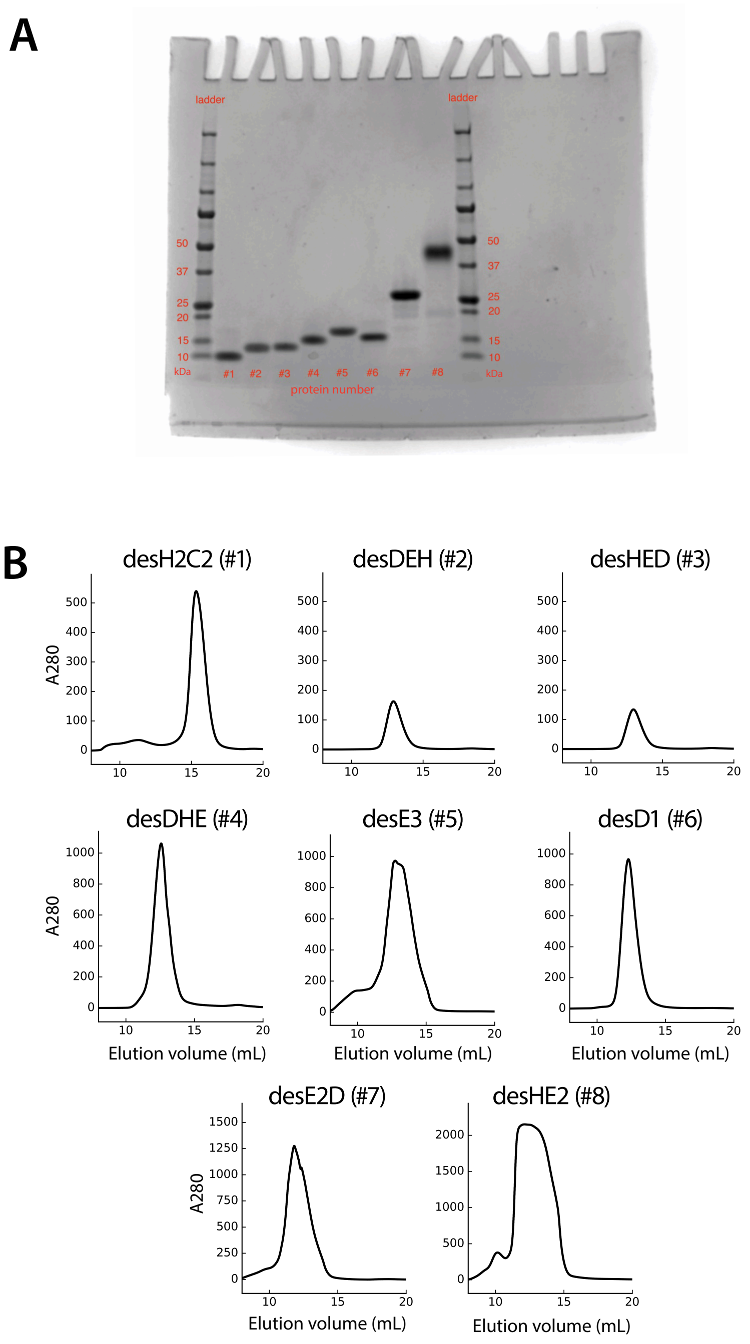


**Figure S6**. (A) SDS-page gel of each protein after SEC post-EDTA incubation. We note that protein 8 (27kDa) shows up at ~50kDa. Size-exclusion chromatography and X-ray structure confirmed the protein is a monomer, suggesting that the protein forms non-covalent, likely ionic bonds as DNA on the gel. (B) Prior to TEV-cleavage, protein 8 had a monomeric band at ~27kDa. Purification of all eight proteins on SEC after TEV-cleavage and EDTA incubation. For each protein, we take the major peak.

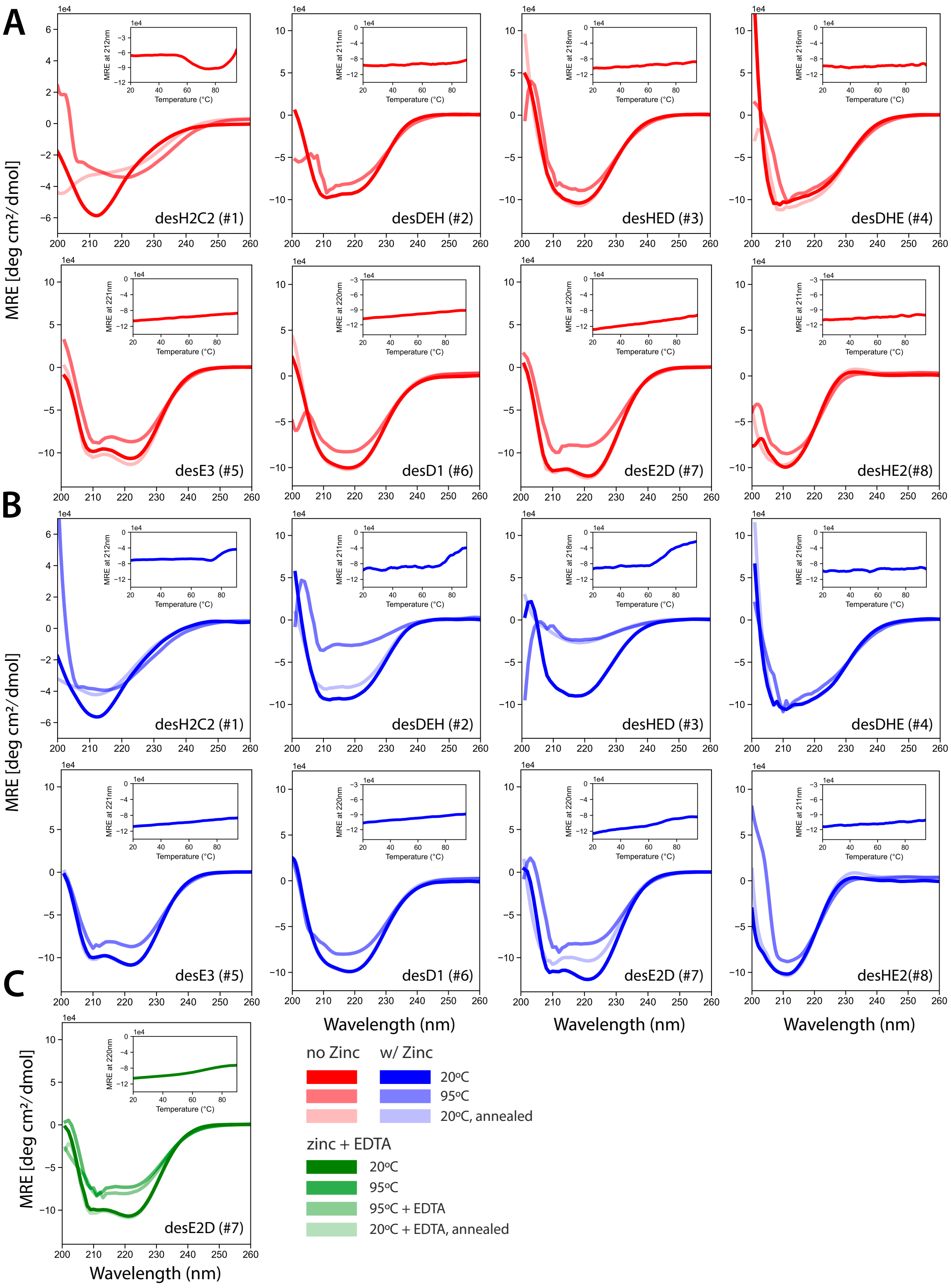


**Figure S7**. Circular dichroism (CD) and temperature melt curves. (A) The CD curve of each protein was collected in the absence of zinc at 20ºC, directly after temperature melt at 95ºC, and directly after re-folding by temperature annealing to 20ºC. The temperature melt curve is in the top right of each respective protein. (B) The same data as in panel A, but in the presence of equimolar zinc. Fresh protein was used for each spectra. Notably, the addition of zinc stops aggregation for desH2C2 (#1), destabilizes desDEH (#2), prevents the refolding of desHED (#3), and causes alternate refolding in desE2D (#7). (c) Upon addition of EDTA after thermal denaturation, desE2D (#7) refolds to its original fold, indicating metal-dependent refolding.

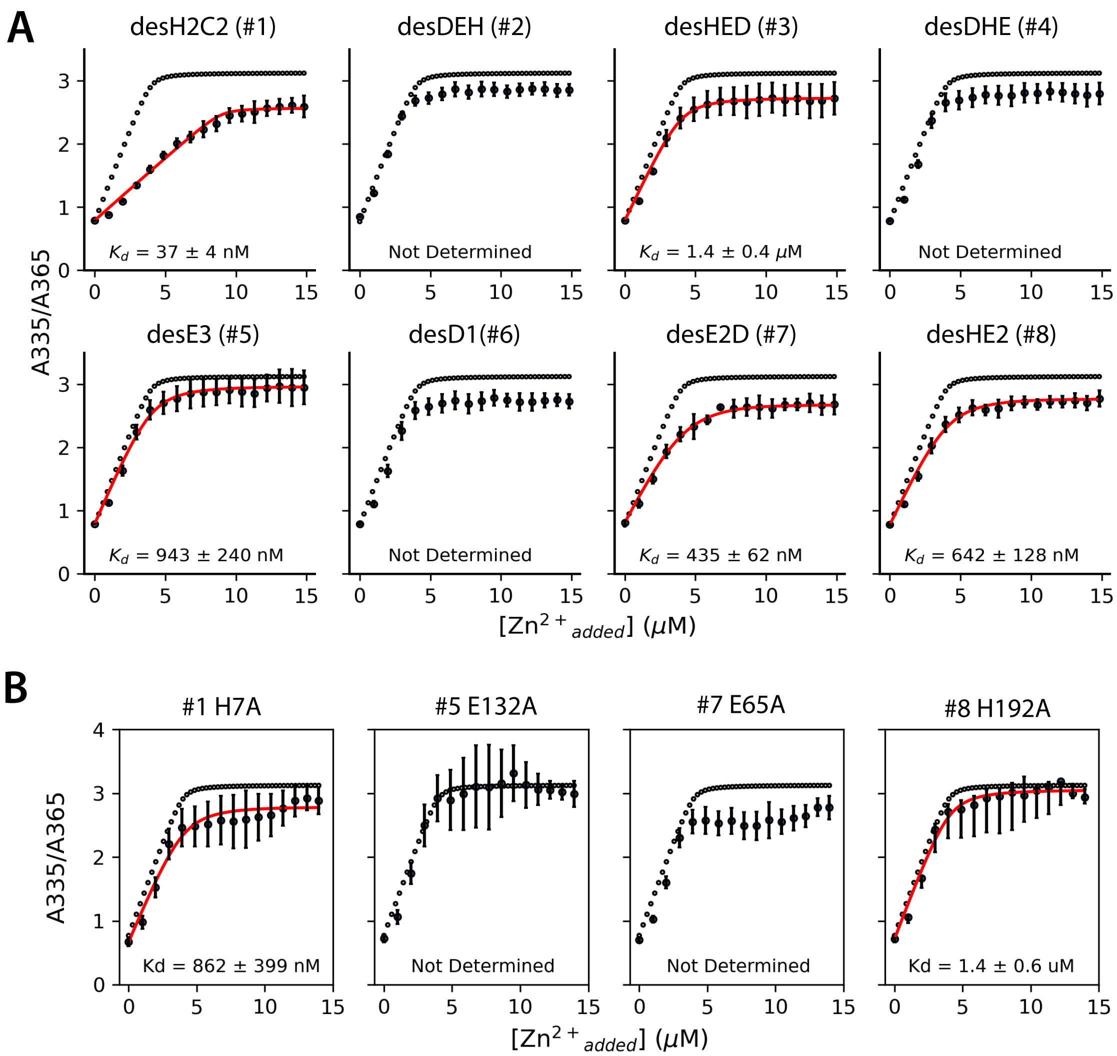


**Figure S8.** Experimental assays on computationally designed metalloproteins. (A) Competition titration assay with MagFura-2. Open circular dots represent experimental curve of MagFura-2 alone and red represents best curve fit. Error bars represent relative error across n=5 replicates. (B) Point mutations of designed proteins. Point mutations are made at a key ligating residue in the designed binding sites. In a competition titration assay with MagFura-2, all mutants have weakened activity.

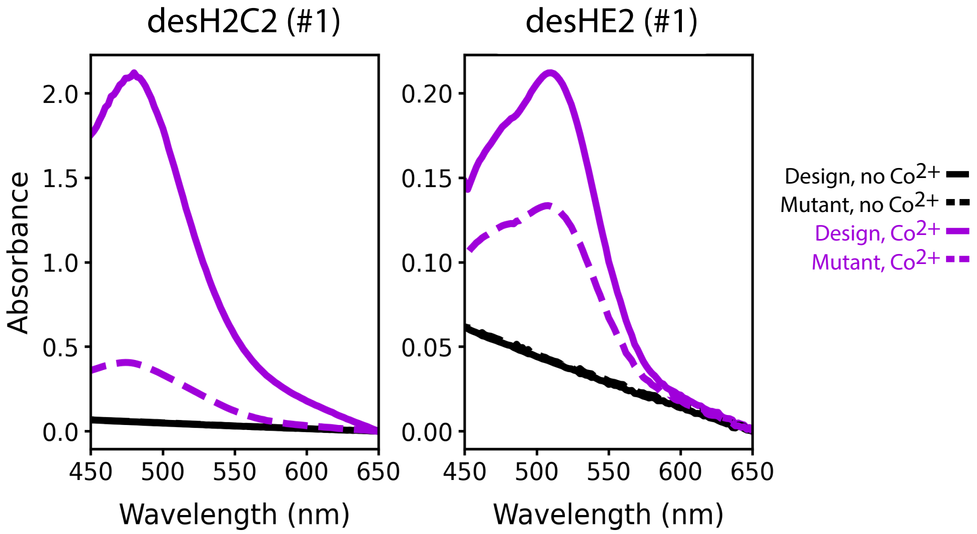


**Figure S9.** Cobalt substitution assay. UV-vis absorbance spectra reveal characteristic Co(II) ligand-to-metal charge transfer bands. desH2C2 exhibits a strong cobalt-dependent absorbance peak while desHE2 exhibits a weaker absorbance intensity, consistent with designed motifs. Mutations of key a designed ligating residue displayed diminished absorbance.

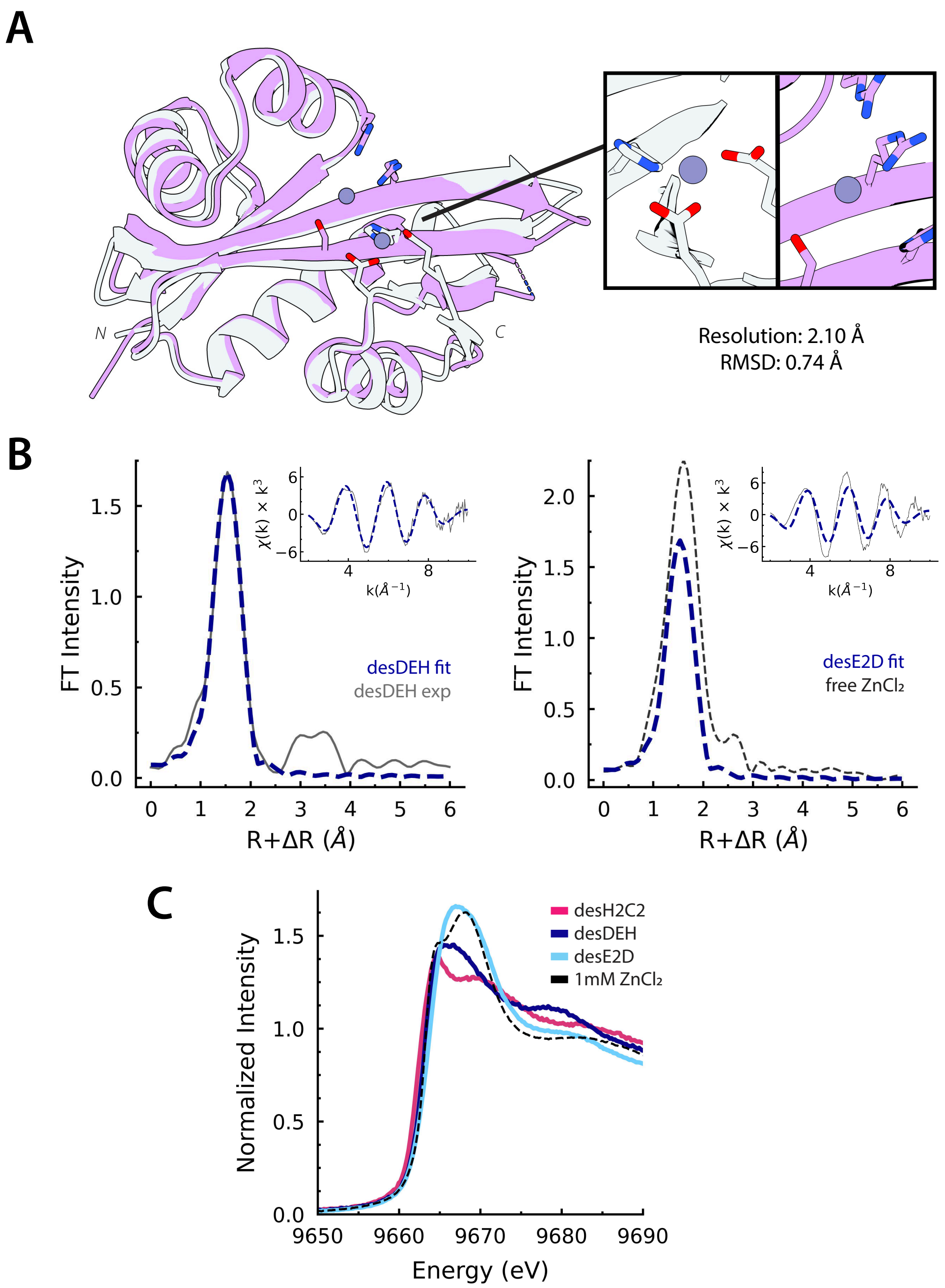


**Figure S10.** (A) Crystal structure of desDEH closely matches the design (Resolution: 2.10Å, RMSD: 0.74Å) demonstrating that dEVA can reliably guide sequence design of *de novo* folds. Although the design model predicts zinc coordination at the C-terminus, this region is disordered in the crystal structure, leaving the zinc without coordinating ligands and further confirming the weak metal binding. (B) Zn K-edge EXAFS spectra of desDEH. The non-phase corrected Fourier transform and the corresponding EXAFS (inset). Fit shown in blue (dashed line) overlayed with the experimental data (solid grey line, left) and 1mM free ZnCl (black dashed line, right). (C) Normalized Zn K-edge spectra of desH2C2, desE2D, desDEH, and free zinc chloride.

### LigandMPNN Analysis

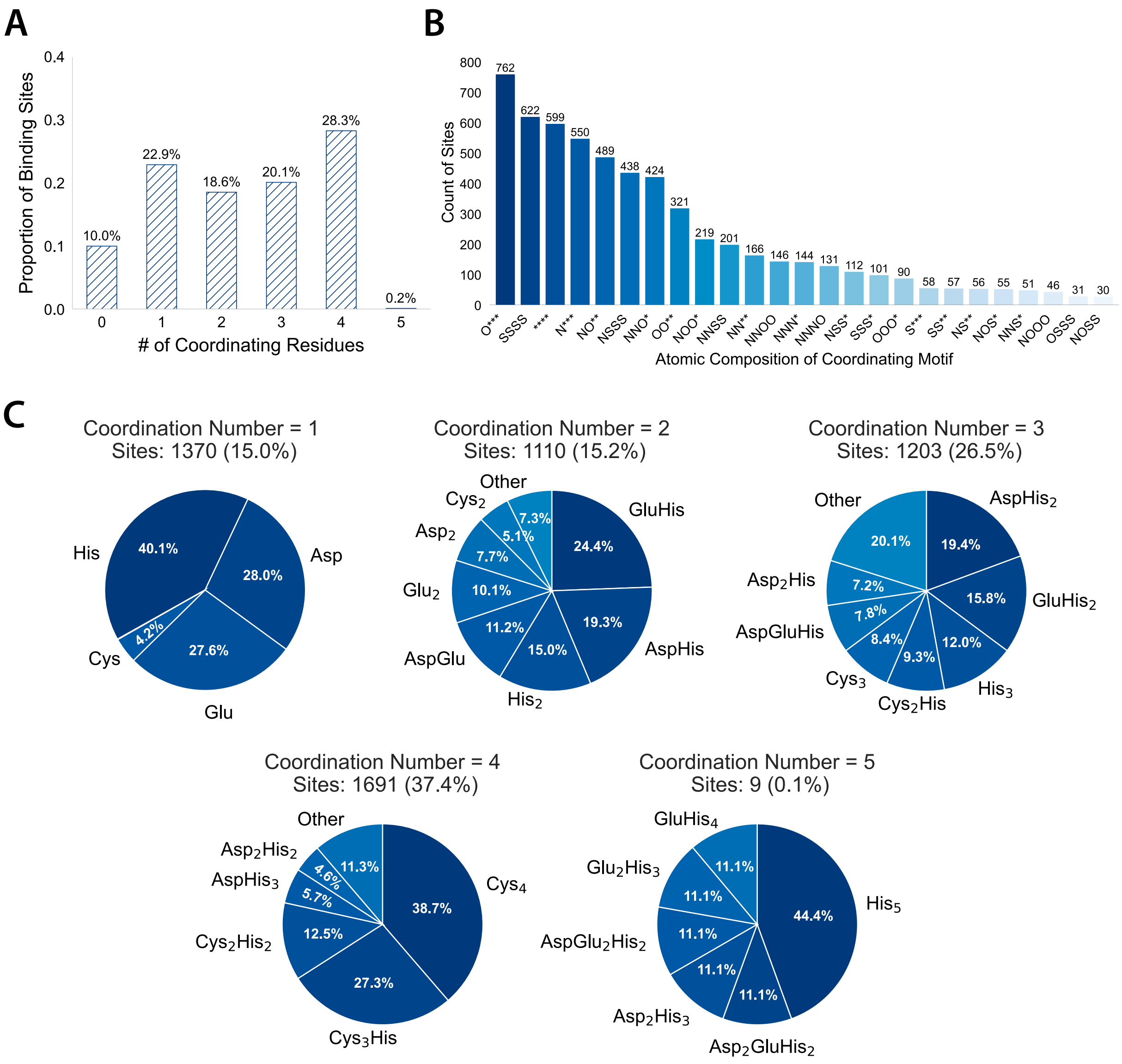


**Figure S11.** Analysis of the LigandMPNN training dataset. (a) The number of coordinating residues per zinc binding site in representative zinc-containing proteins structures. (b) Analysis of the atomic composition of coordination motifs across representative metal-containing proteins. The 25 most common compositions are depicted in descending order of relative frequency. Single- residue carboxylates dominate the binding motifs. Atoms represented are nitrogen (N), oxygen (O), sulfur (S), and an open coordination site (*). (c) Break down of the individual residue contributions per metal binding site.

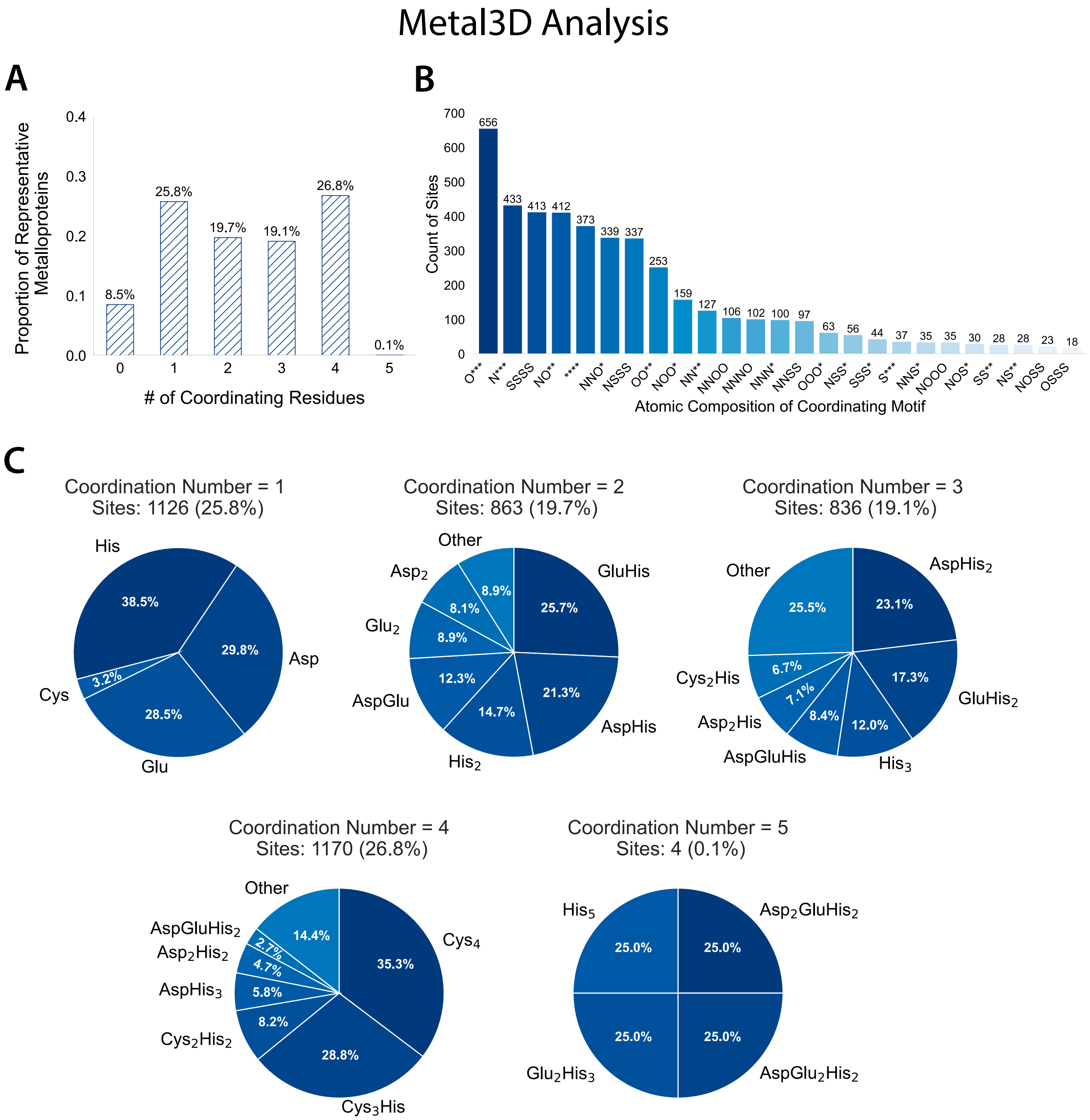


**Figure S12**. Analysis of the Metal3D training dataset. (a) The number of coordinating residues per zinc binding site in representative zinc-containing proteins structures. (b) Analysis of the atomic composition of coordination motifs across representative metal-containing proteins. The 25 most common compositions are depicted in descending order of relative frequency. Single-residue carboxylates dominate the binding motifs. Atoms represented are nitrogen (N), oxygen (O), sulfur (S), and an open coordination site (*). (c) Break down of the individual residue contributions per metal binding site.

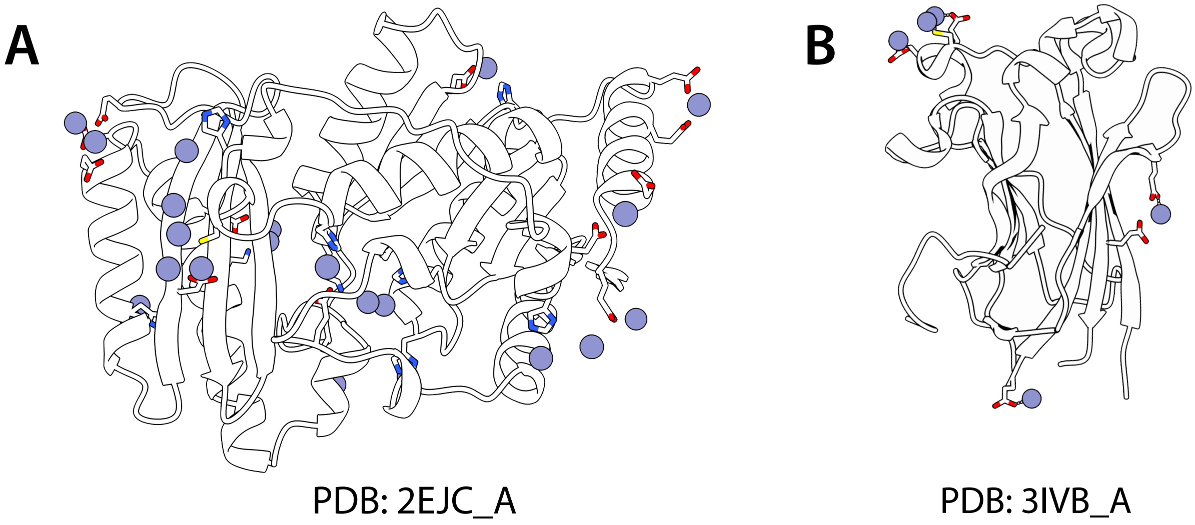


**Figure S13**. Examples of protein structures from the PDB containing metal crystallographic artefacts. (A) structure of PDB entry 2EJC and (B) 3IVB shown as cartoon representations. Blue spheres indicate zinc ions with nearby residues shown as sticks. The metals constitute examples of the crystallographic artefacts that exist in the training dataset of current deep learning models.

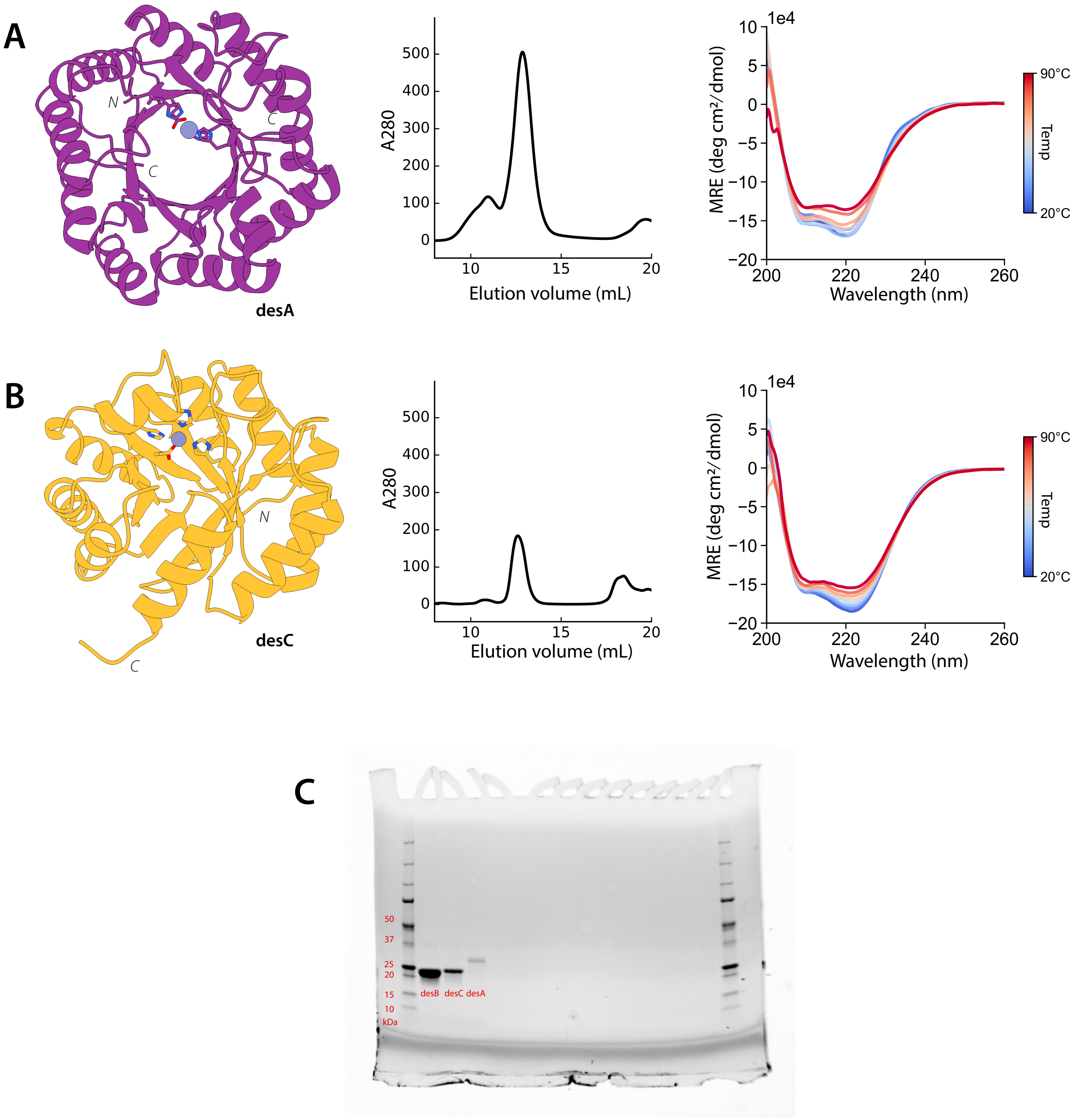


**Figure S14**. Characterization of designed metalloenzymes desA and desC. (A) Designed structure of desA (purple), shown alongside SEC profile and CD thermal melt spectra collected from 20ºC to 90ºC. The SEC trace shows a dominant monodisperse peak, and the CD spectra indicates the protein is folded and stable across temperatures. (B) Designed structure of desC (gold), with corresponding SEC and CD melt. SEC profile shows a monomeric peak and CD spectra confirms secondary structure with high thermostability. (C) SDS-PAGE gel of purified desB, desC, and desA, all migrating to the expected molecular markers, consistent with their expected size.

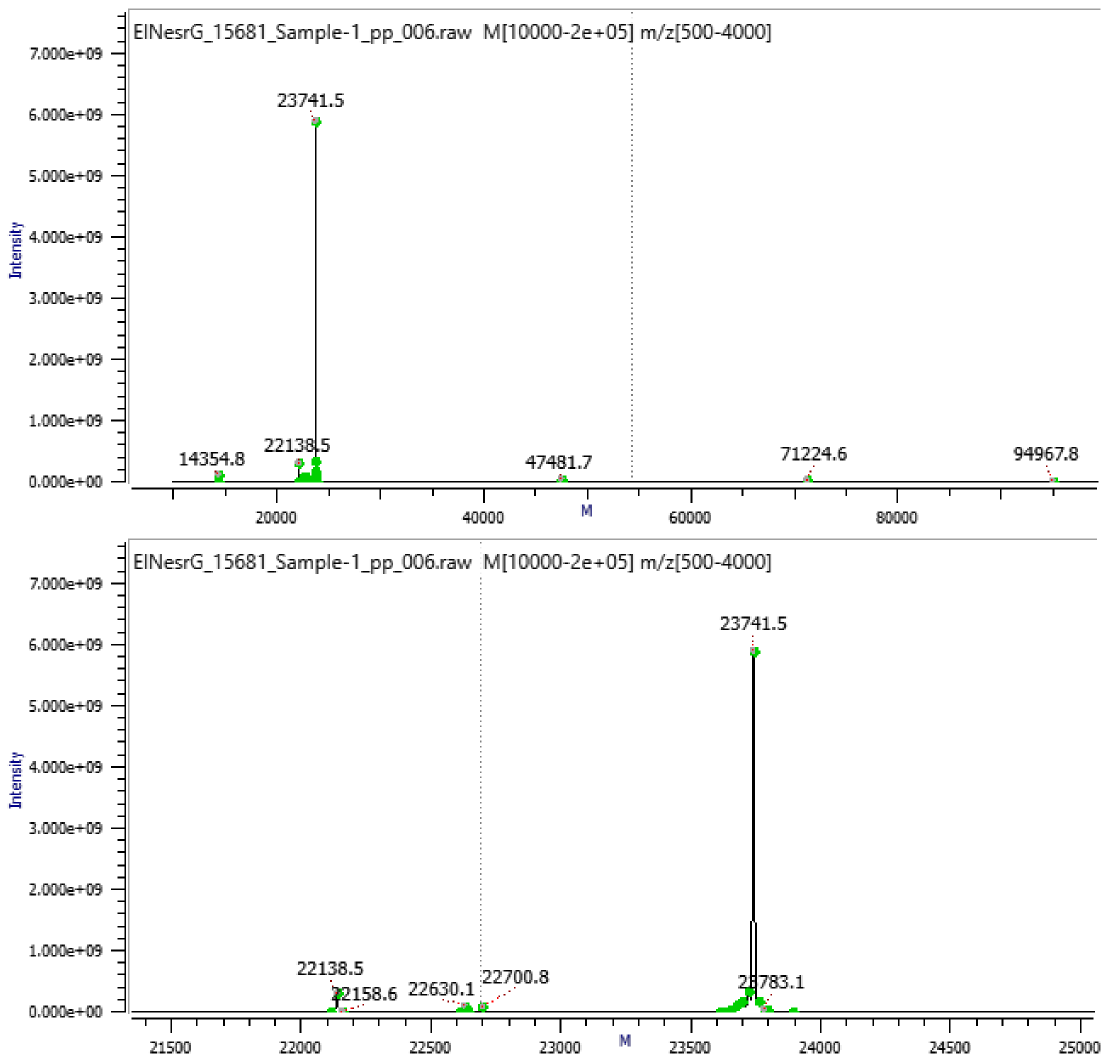


**Figure S15**. LC/MS of desB. The expected molecular weight 23741 Da. The highest peak is 23741, consistent with the expected molecular weight.

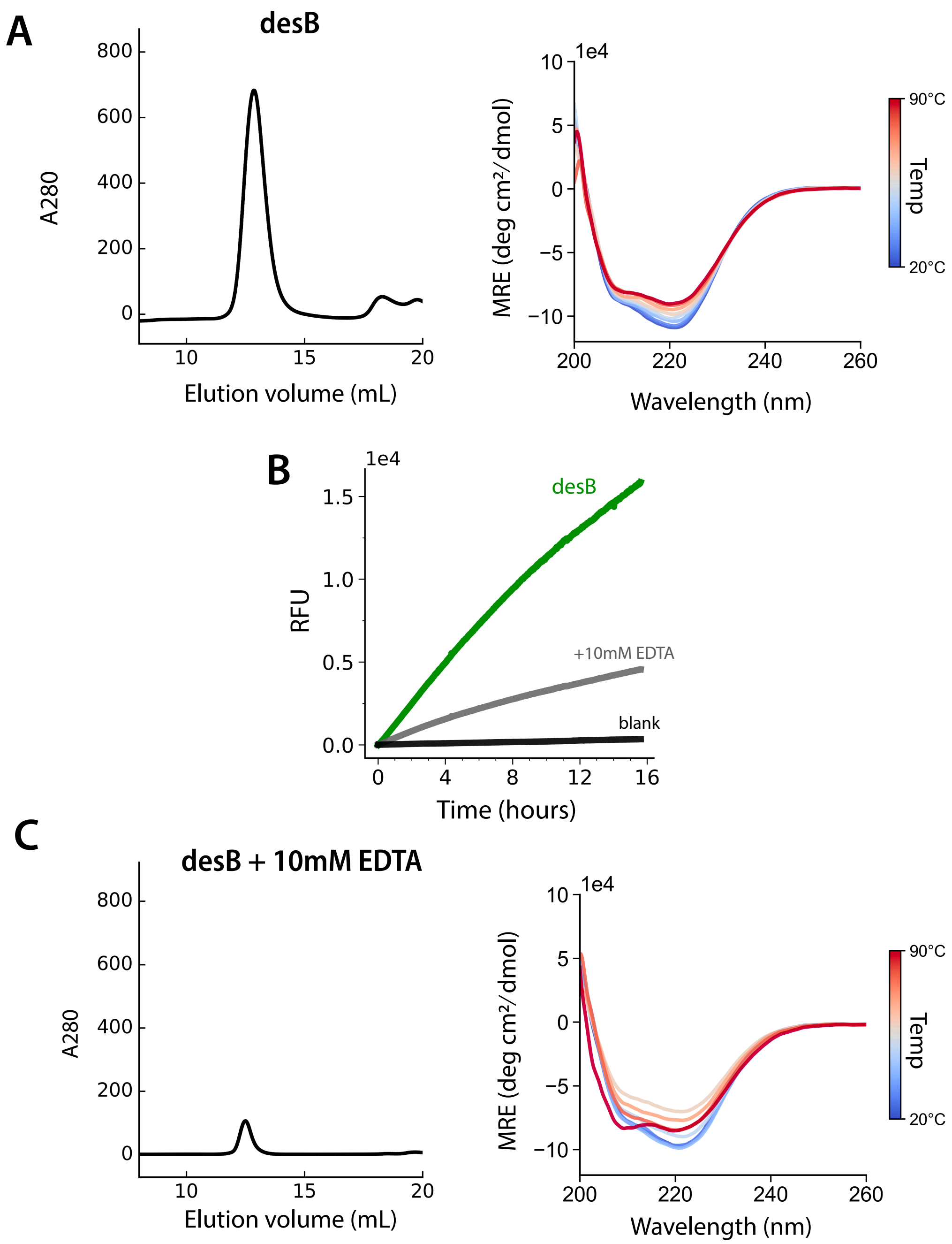


**Figure S16**. Evaluating the zinc-dependence of desB. (A) The size-exclusion chromatography spectra (left) and circular dichroism spectra (right) of desB directly after purification. (B) The addition of 10mM EDTA directly before reading kinetics decreases the turnover of 4MUP. (C) The size-exclusion chromatography spectra (left) and circular dichroism spectra (right) of desB directly after purification. The protein is destabilized after cleavage of any bound metals by EDTA.

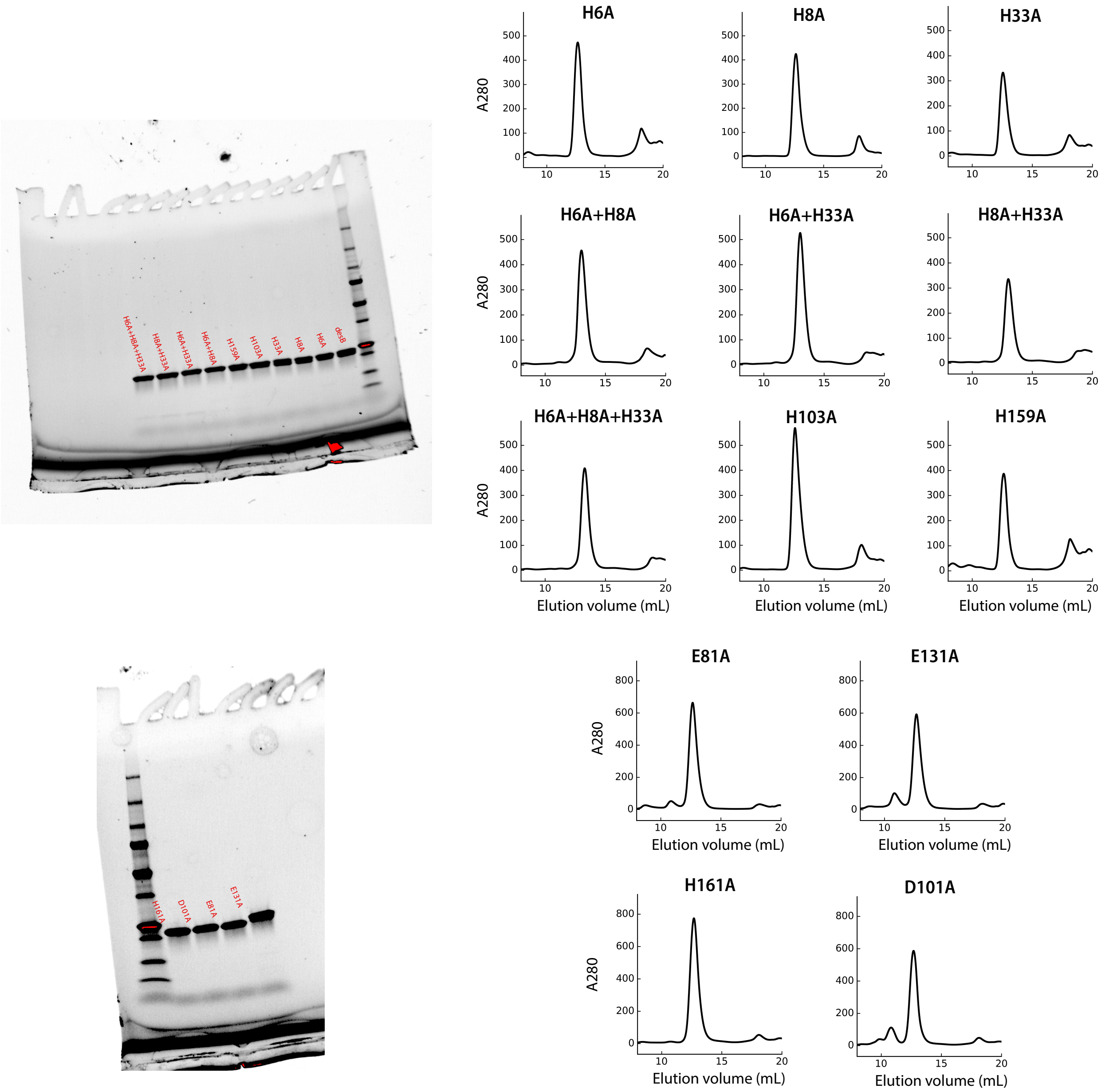


**Figure S17**. SDS-PAGE gel and size exclusion chromatography spectra of all thirteen mutants of desB. All mutants are monomeric and run at the expected size as the original desB design.

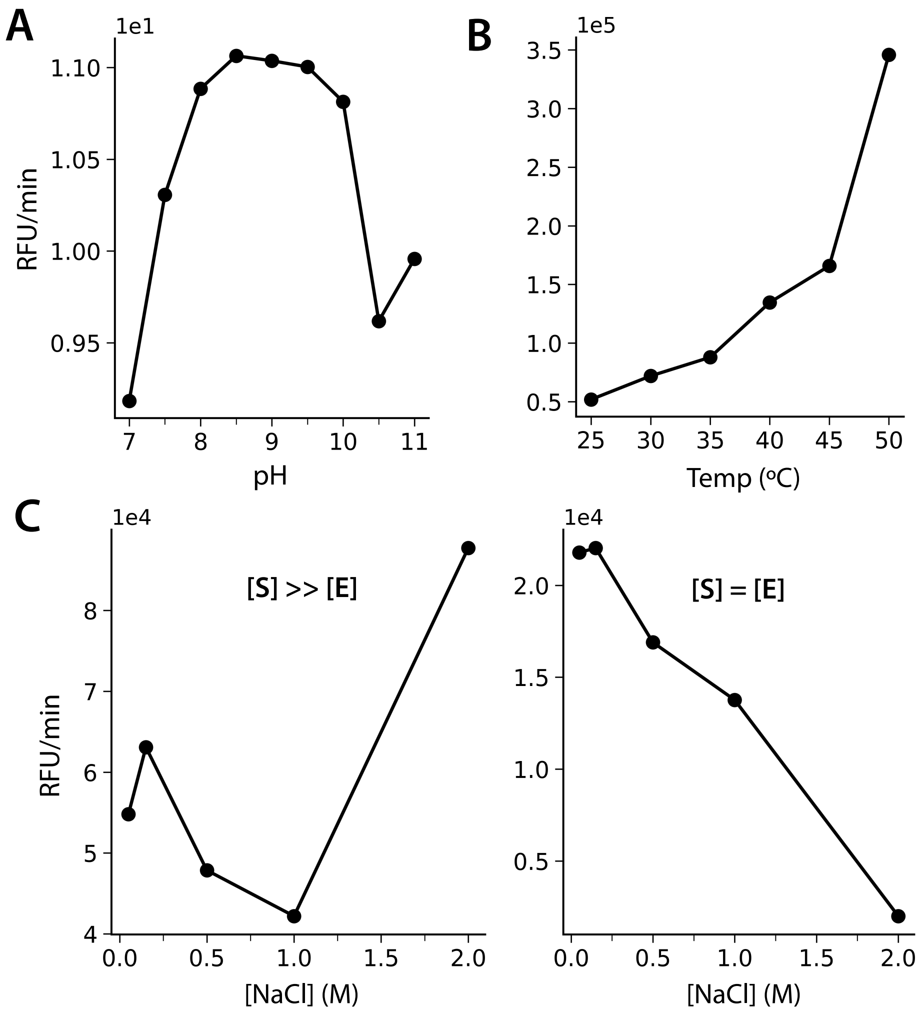


**Figure S18**. Assessing the pH dependence (A), temperature dependence (B), and ionic strength dependence (C) of desB's activity as a phosphate monoester. The improved catalysis at pH 8.5 indicates that hydroxide is involved in hydrolysis.

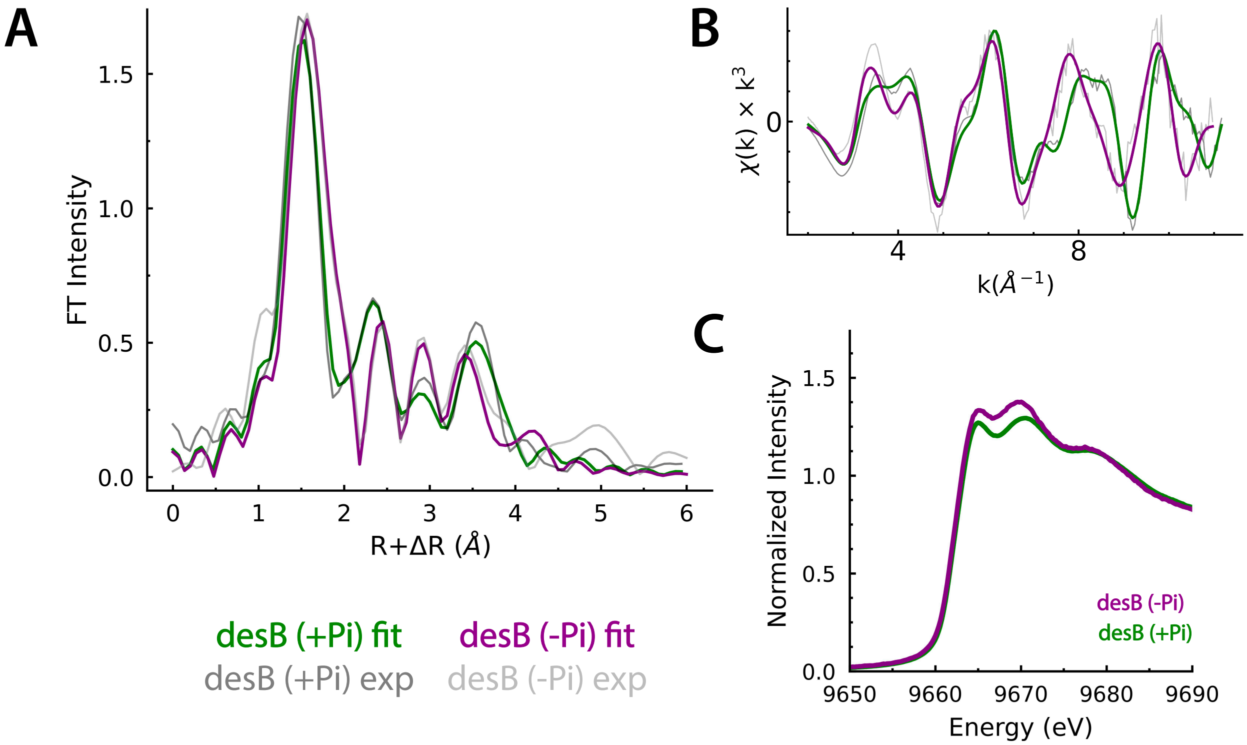


**Figure S19**. EXAFS and XANES analysis of desB in the presence and absence of orthophosphate. (A) Fourier-transformed and (B) Zn K-edge EXAFS spectra of desB (+Pi) (green) and apo desB (-Pi) (purple), with experimental data in gray and fits overlaid. The spectra oscillate in phase below k=7$Å^{-1}$ but diverge at higher k, and a feature near 7.5Å in the +Pi spectrum is absent in the apo sample. The first-shell Zn bond distance is 0.04Å longer in the apo sample, consistent with weak water ligands in the coordination sphere. Features between R'=2-4Å are fit to single-and multiple-scattering paths from coordinating histidine residues. (C) Comparison of the Zn K-edge XANES spectra.

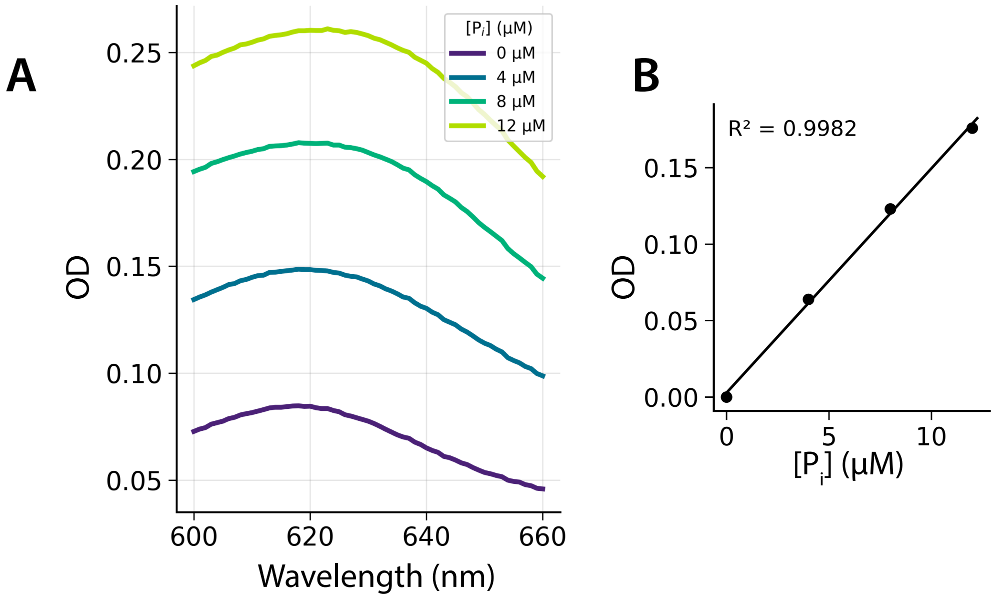


**Figure S20.** Absorbance spectra (600-660nm) of malachite green reagent at increase inorganic concentration (0-12µM), showing characteristic absorption peaks near 620nm. (B) The standard curve relating $P_i$ concentration (µM) to absorbance (optical density, OD) at 620nm), demonstrating excellent linear fit ($R^2 = 0.9982$).

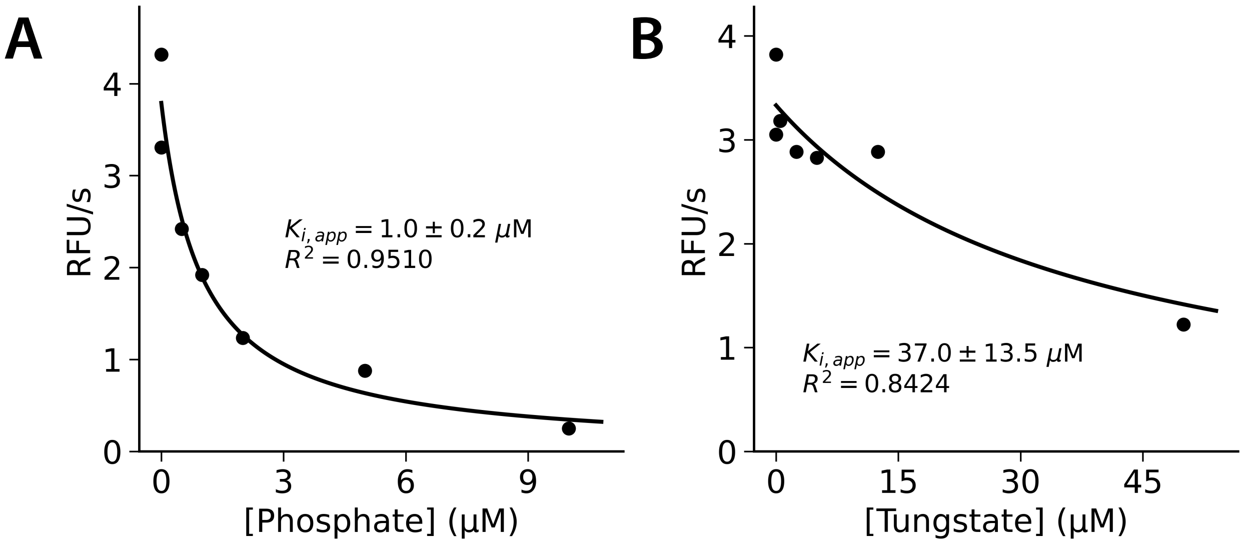


**Figure S21**. Inhibition of phosphomonoesterase activity by phosphate and tungstate. Enzyme activity (RFU/s) as a function of orthophosphate (A) and tungstate (B) concentration for the same biological replicate (n=1). Data is fit to a hyperbolic inhibition model, with fitted Ki,app corrected for substrate competition.

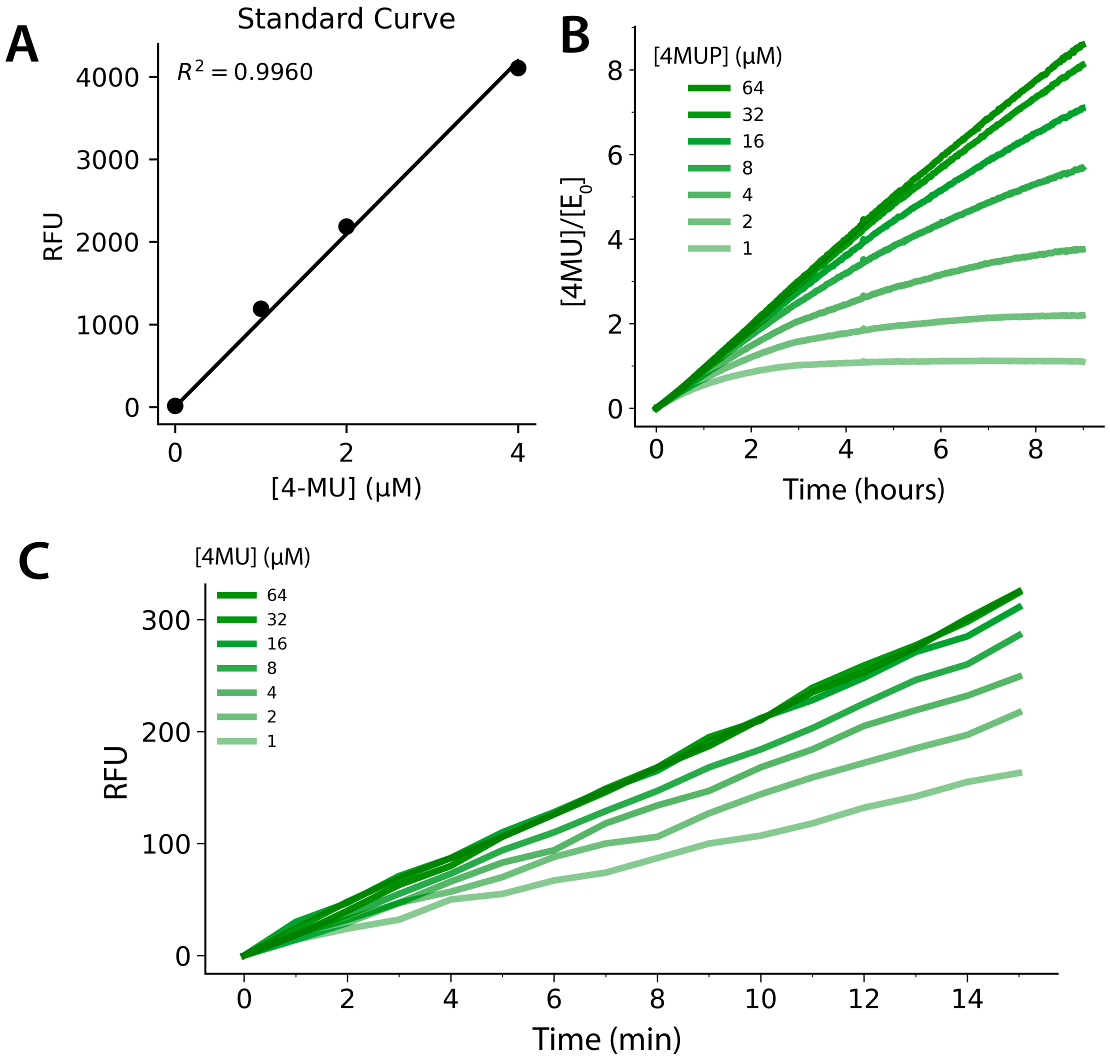


**Figure S22**. Kinetics of enzymatic activity measured by 4MU fluorescence. (A) The standard curve relating 4MU concentration (µM) to relative fluorescence units (RFU) with a strong linear relationship ($R^2 = 0.9960$). (B, C) Time-course fluorescence measurements at varying 4MUP substrate (1=64µM) where [E] = 1µM. Higher substrate concentrations (dark green) produce greater fluorescence signal, consistent with increased production of 4MU.

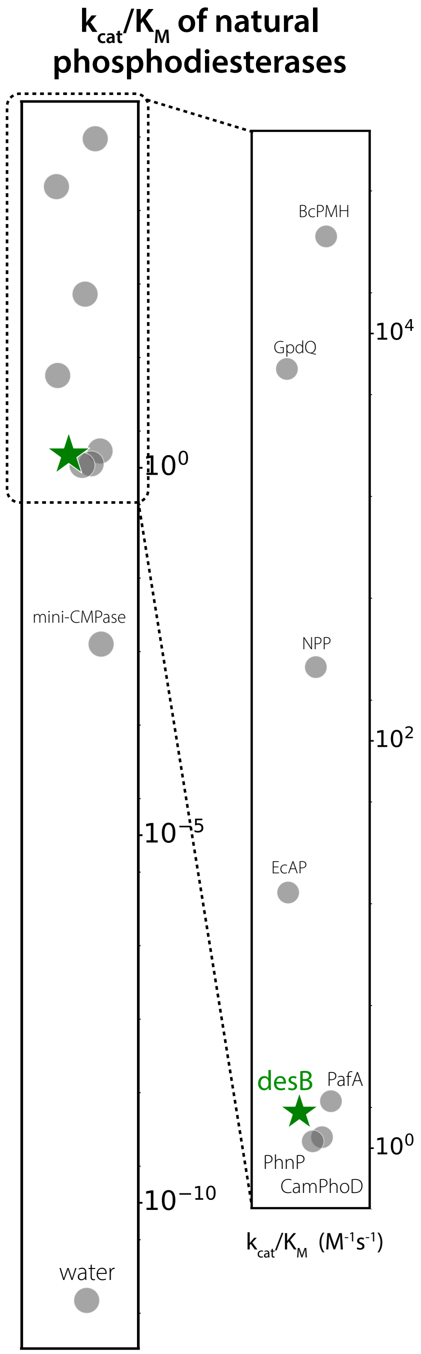


**Figure S23**. Comparison of catalytic efficiency ($k_{cat}/K_M$) of phosphodiesterase activity literature values to desB (dark green star).

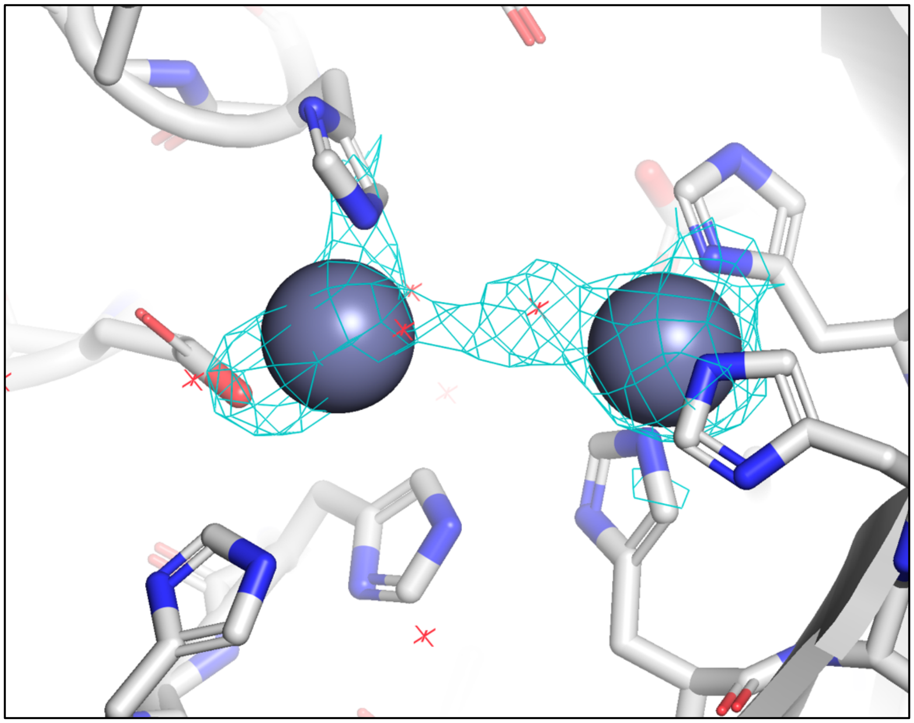

**Figure S24.** X-ray structure showing electron density [composite omit map (σ = 2.5)]. Structure shows density at the di-zinc site and surrounding coordinating residues, shown in cyan wire.

**Table S1A.** Sequences and primers ordered in Phase I for the design of metalloproteins. In bold are the TEV cleavage site and 6x His tag.

| Protein Name | Sequence |
|---|---|
| desH2C2 | MAAVCTNHPSEVTVKRGEAATVTHTFTGLTADPTVFFNGQLVKPGDPDFEVTISGNTVTITIKAGAEIEGGGTITVTAPGGETCTTKIVVVP**SAENLYFQSHHHHHH** |
| desDEH | **MHHHHHHENLYFQSA**MLEVRPMTEADLPALRALARAAGLSDEELEAMLARSEVMVGYVDGEPVAFAFGRLVRRDGKAYIHVSSLADPDTPEELLEMMFRALIEDLLPQAGRAIFVTDGDELAERLARELGLEVEDE |
| desHED | **MHHHHHHENLYFQSA**MNTFTITLTLAGAKVGDGLDLLVATLSSLPHVTLTVSADKKTYTFEVKVGDKTITFTLTVVERTPTTTVLALTYGDIKGTVTITDSTGADGNTVVTITFVSDADAETHKAIANELASDLERGFQEHPPVPGVTLVSIELTF |
| desDHE | **MHHHHHHENLYFQSA**DPIEFTRFEQCLVLDARLLTDPAQPEAPWVVELDLDPSCGDHIRPGDILAFAVDGELQAIGRVLGPDPSIPDKPPGESVLVELASREEADALIAMKGKEVLIMIVRAPGLSTEEALAYVLSPEGRALLTAEIQRALAENPDLL |
| desE3 | MKPAVGLIPPELMETLGAAFRAGGAPLSPEEVGVIAGLAIGAIVSAMNRGVDAEEAAQEVEELLEGMLAEGASLEEIIATIEKRYPGVRVLSIDEDGITVSFLVPDDPEKATELIEEATEAAVAMMADIAREAGASDAQVATVREAVREIGENFLAGALAS**SAENLYFQSHHHHHH** |
| desD1 | MAMLPASDFFEETVPGTVTFKEVVADPESPGSQKVLDRLIDLALDAASKVPGRIVTFIFRFRQDPEDPNGWQIDILIYVDGKPVAVAGVTFNTDGTLTFYRRGITAEEYDALEALPAILRITIVVDVEAAKILEREVEVSPLVSELGLSKEEALASAEIIADFATKKLVAWARELRAA**SAENLYFQSHHHHHH** |
| desE2D | **MHHHHHHENLYFQSA**SPKQREIIAAMREVAPRLEIYLLGDHIVLEVSEEAYRADPEGILAELAAAIRAAAELGGKNIVIEFDFEPGTDVIALLREIREGLARAPGITPEMLKKVEFMVGMSLIGLAELGLRAAGLDLDLAEALADPELKATLLAAVKALLTELAEEHGVDAITFVIPPSSTPISREEALRVLAALGADVVAHAQSLGLDVRVFSPEKALTPEEAAAANAAGLPGIYSAEDLEEERRRFE |
| desHE2 | **MHHHHHHENLYFQSA**TPRLLAVGINSAKKKVHLLDASTNKILATFTVPDGSTPTAATLADGILYIGTDNGKIFAFDVETKELVRTIQAHPEGTVITSIAVGPDGKIYTGTSNGLWAILDGETGEILHTGQVPEPIVSVTFSPDGKTLYLATPNAIYEWDVASRELVEVIELPAEYQGKVTDIAMGDDGNLYFVTSDGDIIRLDVETHKFTKATSLGGGKPTAIAAGNGKIYVADAKRRQILEFDERTFELERVEATLPSDVTVVSIVVV |

**Table S1B.** Sequences and primers ordered in Phase I for the design of metalloproteins. In bold are the TEV cleavage site and 6x His tag.

| Protein | Mutant | Forward Primer | Reverse Primer |
|---|---|---|---|
| desH2C2 | H7A | TACAAATGCACCCTCAGAAGTCACGGTG | GAGGGTGCATTTGTACATACTGCAGCCATCTCG |
| desH2C2 | H23A | CGTGACGGCAACCTTCACCGGTTTGACC | AAGGTTGCCGTCACGGTGGCTGCCTC |
| desHED | H31A | CCTGCCGGCAGTGACCTTAACTGTCAGCGCC | GTCACTGCCGGCAGGCTAGAAAGGGTC |
| desE3 | E61A | GGAGGTGGCAGAATTGCTGGAGGGCATGCTG | AATTCTGCCACCTCCTGCGCAGCCTC |
| desE3 | E132A | CGCGAGAGCAGCTGGCGCGAGCGACGCC | CCAGCTGCTCTCGCGATGTCTGCCATCATTGC |
| desE2D | E65A | CGTGATCGCATTCGATTTCGAACCGGGAACC | TCGAATGCGATCACGATGTTTTTACCGCCAAGC |
| desE2D | H26A | GGGTGATGCAATTGTCCTGGAAGTGAGCGAAGA | ACAATTGCATCACCCAGCAGGTAGATTTCCAG |
| desHE2 | H192A | AGAGACTGCAAAATTTACCAAAGCGACGTCCCT | AATTTTGCAGTCTCTACGTCCAGACGAATGATG |

**Table S2.** Computational metrics of the sequences for each of the eight computationally designed metalloproteins. The p(metal) is calculated by Metal3D, and p(seq) is calculated by LigandMPNN.

| **Protein** | **desH2C2** | **desDEH** | **desHED** | **desDHE** | **desE3** | **desD1** | **desE2D** | **desHE2** |
|---|---|---|---|---|---|---|---|---|
| Length | 91 | 121 | 141 | 143 | 161 | 177 | 234 | 254 |
| p(metal) | 0.98 | 0.80 | 0.93 | 0.96 | 0.89 | 0.56 | 0.93, 0.87 | 0.79 |
| p(seq) | 0.49 | 0.49 | 0.43 | 0.44 | 0.49 | 0.48 | 0.46 | 0.54 |
| FoldSeek TM-score | 0.82 | 0.71 | 0.71 | 0.54 | 0.55 | 0.42 | 0.55 | 0.81 |
| BLAST seq sim | 43% | 32% | 29% | 31% | 28% | 38% | 27% | 32% |

**Table S3.** ICP-MS detection of transition metals in the apo metalloproteins confirms borderline to minimal trace transition metals in each designed protein.

| **Protein** (apo) | **Cr (ng)** | **Mn (ng)** | **Fe (ng)** | **Co (ng)** | **Ni (ng)** | **Cu (ng)** | **Zn (ng)** |
|---|---|---|---|---|---|---|---|
| 1 | 0.05 ± 0.00 | bdl | bdl | bdl | 0.56 ± 0.01 | bdl | 0.97 ± 0.49 |
| 2 | 0.92 ± 0.03 | bdl | 2.78 ± 0.31 | bdl | 58.41 ± 2.05 | bdl | 5.95 ± 1.66 |
| 3 | 0.31 ± 0.02 | bdl | bdl | bdl | 0.88 ± 0.04 | bdl | 0.49 ± 0.62 |
| 4 | 0.11 ± 0.01 | bdl | bdl | bdl | 17.95 ± 0.25 | bdl | 1.23 ± 0.65 |
| 5 | 0.04 ± 0.04 | 0.11 ± 0.01 | bdl | bdl | 9.02 ± 0.20 | bdl | 169.15 ± 2.16 |
| 6 | 0.51 ± 0.27 | bdl | 0.18 ± 0.05 | bdl | 13.66 ± 0.13 | bdl | 1.51 ± 0.65 |
| 7 | 0.03 ± 0.01 | bdl | bdl | bdl | 10.72 ± 0.41 | bdl | 4.21 ± 2.16 |
| 8 | 0.03 ± 0.04 | bdl | bdl | bdl | 4.31 ± 0.13 | bdl | 1.58 |
| *detection limit* | 0.125 | 0.018 | 1.260 | 0.012 | 0.075 | 0.676 | 1.354 |

**Table S4.** Zn K-edge EXAFS fitting results for *de novo* designed proteins.

| Protein | Coordination/path | $R$ (Å)* | $\sigma^2$ (Å$^2$)† | $\Delta E_0$ (eV) | F‡ |
|---|---|---|---|---|---|
| desH2C2 (#1) | 2 Zn-N | 1.97 | 462 | -11.844 | 0.441 |
| | 2 Zn-S | 2.27 | 383 | | |
| | 4 Zn-C | 2.50 | 462 | | |
| desDEH (#2) | 2 Zn-N/O | 1.92 | 147 | -1.72 | 0.256 |
| | 2 Zn-N/O | 2.08 | /147 | | |
| | 2 Zn-C/O | 2.53 | 244 | | |
| des E2D (#7) | 2 Zn-N/O | 1.97 | 269 | -1.30 | 0.205 |
| | 2 Zn-N/O | 2.12 | 229 | | |
| | 2 Zn-C/O | 2.88 | 894 | | |
| desB (+Pi) | 4 Zn-N | 1.96 | 367 | -2.04 | 0.193 |
| | 1 Zn-P | 2.79 | 234 | | |
| | 4 Zn-C | 2.97 | 447 | | |
| | 8 Zn-C-N | 3.19 | /447 | | |
| | 4 Zn-C | 4.04 | 99 | | |
| | 12 Zn-N-N | 4.14 | /99 | | |
| | 1 Zn-Zn | 4.31 | 128 | | |
| | 8 Zn-C-N | 4.70 | 564 | | |
| desB (apo) | 4 Zn-N | 2.01 | 290 | -2.71 | 0.283 |
| | 1 Zn-O | 2.41 | 54 | | |
| | 4 Zn-C | 3.00 | 107 | | |
| | 8 Zn-C-N | 3.18 | /107 | | |
| | 4 Zn-C | 3.28 | 1621 | | |
| | 8 Zn-N-N | 4.15 | 474 | | |
| | 8-Zn-C-N | 4.58 | /457 | | |
| | 16 Zn-C-N | 4.66 | 987 | | |

*The estimated standard deviations for the bond lengths are on the order of ±0.02 Å.
†A slash indicates that the $\sigma^2$ value was linked to the previous path.
‡The error is given by $[\Sigma k^6(\chi_{exptl}-\chi_{calcd})^2/\Sigma k^6\chi_{exptl}^2]^{1/2}$. The $S_0^2$ factor was set at 1.

**Table S5.** Sequences ordered in Phase II for the design of metalloenzymes. In bold is the Strep-Tag(s). The tandem tag in desB was an unintended cloning duplication.

| Protein Name | Sequence |
|---|---|
| desA | MARVEFWVHAGPLSGVTGEDVARLIEAGAAVVVLWVVGMDGVMPEETLREIAEALRRHPEVKVVAWVSAGGALKSGIDETVRALRLAMELAGVDRVMLYGGVNEKGEPLPLEDFLRIARAVAEVAREHGAEVWFHLSGGAKTPISEEELLRYAEAAVREAGVDGVHVTPRDENGNKIRPSPEEVARLIRRLKELGAKHVVVSVDASTPVEEIVRIAKVAQEEGGMVEIHADPSLPVEEAARRMAEQARALKEAGVDAIVTTPSGATPEEQLAMLEALREAGIEAVPL**SAWSHPQFEK** |
| desB | MRLWGHFHARSREEARRAVRLAKLAGFDGILVHGSTPEVVAEATRVIKEETDLPVIAEISGGVEKAREARAKTDADEVVAETNSLEEALQLVREGAVDAVDLHNALKRYSLEEVVEMVRAIKAAGGRVFLEVDTGPGGATLEEVLEAIRASGGAIEGVHTHDATPEVIAALREAGVKEIFVSLPETEEEARRLGELAR**SAWSHPQFEKSAWSHPQFEK** |
| desC | MRAPRLPRGALLVGLSVGGPDNPTLEAVIERARRARELGAAGIVLLASVTPHKGLPTATPEQLAAAVRAIREETGLPVFLHGVALTPEQADRLVEMAEAAGVDAIVVPSSTPLEALRRIRERGIPVFLELHDEETAKALHPGLKPVKPEEIEELIRRAKEAGFEGVVLSTNDPELARKGAEAARREGVPFFVRTNGDVETVRAIAEIPEVEGIILDGSWEKFEEAARALR**SAWSHPQFEK** |

**Table S6.** Primers to introduce active site mutations into desB.

| Forward Primer | Reverse Primer |
|---|---|
| ATGGGGAGCGTTTCACGCTAGGTCTCGTGAGGAGG | CGTGAAACGCTCCCCATAGTCTCATATGTATATCTCCTTC |
| ACATTTTGCGGCTAGGTCTCGTGAGGAGGCCCGC | ACCTAGCCGCAAAATGTCCCCATAGTCTCATATGTATAT |
| TCTGGTTGCGGGTAGCACCCCGGAAGTTGTTGCAG | GCTACCCGCAACCAGAATACCATCAAAACCTGCCAGCTT |
| ATGGGGAGCGTTTGCGGCTAGGTCTCGTGAGGAG | CCGCAAACGCTCCCCATAGTCTCATATGTATATCTCCTTC |
| ACATTTTGCGGCTAGGTCTCGTGAGGAGGCCCGC | ACCTAGCCGCAAAATGTCCCCATAGTCTCATATGTATAT |
| ATGGGGAGCGTTTCACGCTAGGTCTCGTGAGGAGG | CGTGAAACGCTCCCCATAGTCTCATATGTATATCTCCTTC |
| TCTGGTTGCGGGTAGCACCCCGGAAGTTGTTGCAG | GCTACCCGCAACCAGAATACCATCAAAACCTGCCAGCTT |
| ACATTTTGCGGCTAGGTCTCGTGAGGAGGCCCGC | ACCTAGCCGCAAAATGTCCCCATAGTCTCATATGTATAT |
| TCTGGTTGCGGGTAGCACCCCGGAAGTTGTTGCAG | GCTACCCGCAACCAGAATACCATCAAAACCTGCCAGCTT |
| ATGGGGAGCGTTTGCGGCTAGGTCTCGTGAGGAG | CCGCAAACGCTCCCCATAGTCTCATATGTATATCTCCTTC |
| ACATTTTGCGGCTAGGTCTCGTGAGGAGGCCCGC | ACCTAGCCGCAAAATGTCCCCATAGTCTCATATGTATAT |
| TCTGGTTGCGGGTAGCACCCCGGAAGTTGTTGCAG | GCTACCCGCAACCAGAATACCATCAAAACCTGCCAGCTT |
| GGACCTGGCGAACGCGCTGAAACGTTACAGCTTGG | GCGCGTTCGCCAGGTCCACAGCGTCCACCG |
| AGGCGTGGCGACCCATGATGCGACTCCGGAAGTG | CATGGGTCGCCACGCCTTCAATCGCGCCACCG |
| CGTGGCCGCGACCAATTCCCTTGAGGAGG | AATTGGTCGCGGCCACGACTTCATCGGCATCG |
| TTTTTTGGCGGTTGACACCGGTCCGGGTG | TGTCAACCGCCAAAAAAACACGGCCACCCG |
| GCACACCGCGGATGCGACTCCGGAAGTGATCGC | TCGCATCCGCGGTGTGCACGCCTTCAATCGCGC |
| CGCTGTGGCGCTGCACAACGCGCTGAAAC | TGTGCAGCGCCACAGCGTCCACCGCGC |

**Table S7.** ICP-MS of desB after purification and after zinc supplement.

| Sample | ICP-MS Replicate | Mg (ng) | K (ng) | Ca (ng) | Fe (ng) | Co (ng) | Ni (ng) | Cu (ng) | Zn (ng) | Expected for 1:1 | Expected for 1:2 |
|---|---|---|---|---|---|---|---|---|---|---|---|
| SIGMA blank | 1 | bdl | bdl | bdl | bdl | bdl | bdl | bdl | bdl | n/a | n/a |
| | 2 | 121.344 | bdl | 167.132 | bdl | bdl | bdl | bdl | bdl | | |
| | 3 | 11.913 | bdl | bdl | bdl | bdl | bdl | bdl | bdl | | |
| desB (purified) | 1 | bdl | bdl | bdl | bdl | bdl | 26.194 | 6.803 | 125.186 | **~193** | ~386 |
| | 2 | 39.579 | 2.375 | 33.719 | bdl | bdl | 19.791 | 4.291 | 135.003 | | |
| | 3 | 1.645 | bdl | bdl | bdl | bdl | 19.914 | 3.911 | 133.198 | | |
| desB (zinc) | 1 | bdl | bdl | bdl | 12 | bdl | bdl | 92.355 | 549.462 | ~281 | **~562** |
| | 2 | 17.874 | bdl | bdl | 15.34 | bdl | bdl | 75.263 | 568.763 | | |
| | 3 | bdl | bdl | bdl | 13.929 | bdl | bdl | 69.563 | 567.695 | | |
| reaction buffer | 1 | bdl | 43.649 | bdl | bdl | bdl | bdl | bdl | bdl | n/a | n/a |
| | 2 | 7.7 | 41.33 | bdl | bdl | bdl | bdl | bdl | bdl | | |
| | 3 | 28.988 | 35.67 | bdl | bdl | bdl | bdl | bdl | bdl | | |

**Table S8**. Curated $k_{cat}/K_M$ of characterized phosphomonoesterases in literature, listed from lowest to highest catalytic efficiency.

| **Enzyme** | **$k_{cat}/K_M$ ($M^{-1}s^{-1}$)** | **Substrate** | **Type** | **DOI** | **Origin** |
|---|---|---|---|---|---|
| p34 | 6.40E-01 | pNPP | p34 | 10.1021/bi0255064 | discovered, native |
| NPP | 1.1 | pNPP | nucleotide phosphodiesterase | 10.1021/bi4010045 | native |
| GpdQ | 5 | pNPP | phosphodiesterase | 10.1021/acs.biochem.6b00297 | native |
| Cdc25A | 7.86 | pNPP | dual specificity dephosphorylate and cytokine-dependent kinase | 10.1074/jbc.M109636200 | native |
| CamPhoD | 7.99 | pNPP | *Cobetia amphilecti* KMM 296 | 10.3390/md17120657 | native |
| BcPMH | 2.20E+01 | pNPP | *Burkholderia caryophilli* PG2952 | 10.1073/pnas.0903951107 | native |
| MKP5 | 1.09E+02 | pNPP | Mitogen-activated protein kinase phosphatase-5 | 10.1074/jbc.M203969200 | native |
| VHR | 3.96E+02 | pNPP | Vaccinia H1-related (VHR) phosphatase | 10.1074/jbc.M203969200 | native |
| PP2CA | 8.59E+02 | pNPP | Serine/Threonine Phosphatase | 10.1074/jbc.274.29.20336 | native |
| **desB** | **1288** | **4MUP** | ***de novo*** | **This work** | **designed** |
| Stp1 | 2.60E+04 | pNPP | Protein tyrosine phosphatase | 10.1021/bi500765p | native |
| PTP1B | 9.30E+04 | pNPP | Protein tyrosine phosphatase | 10.1021/bi500765p | native |
| PafA | 1.60E+06 | pNPP | *Chryseobacterium meningosepticum* alkaline phosphatase | 10.1074/jbc.M117.788240 | native |
| PAP | 5.8E+06 | pNPP | Purple acid phosphatase | 10.1006/abbi.1997.0250 | native |
| EcAP | 5.80E+08 | pNPP | *E. coli* alkaline phosphatase | 10.1021/bi500765p | native |
| water ($k_w$) | 5.00E-11 | pNPP | | 10.1021/bi0028892 | |

**Table S9**. Curated $k_{cat}/K_M$ of characterized phosphodiesterase in literature, listed from lowest to highest catalytic efficiency.

| **Enzyme** | **$k_{cat}/K_M$ ($M^{-1}s^{-1}$)** | **Substrate** | **Type** | **DOI** | **Origin** |
|---|---|---|---|---|---|
| mini-cAMPase | 4.0E-03 | bis-pNPP | Miniaturized protein | 10.1038/s41557-024-01490-4 | designed |
| PhnP | 1.08 | bis-pNPP | Purine nucleoside phosphorylase | 10.1021/bi2005398 | native |
| CamPhoD | 1.133 | bis-pNPP | *Cobetia amphilecti* KMM 296 | 10.3390/md17120657 | native |
| **desB** | **1.5** | **me-pNPP** | ***de novo*** | **This work** | **designed** |
| PafA | 1.7 | me-pNPP | *Chryseobacterium meningosepticum* alkaline phosphatase | 10.1021/jacs.6b06186 | native |
| EcAP | 1.80E+01 | me-pNPP | *E. coli* alkaline phosphatase | 10.1016/j.jmb.2008.09.059 | native |
| NPP | 2.30E+02 | me-pNPP | nucleotide phosphodiesterase | 10.1021/bi060847t | native |
| GpdQ | 6.70E+03 | me-pNPP | *Enterobacter aerogenes* glycerol-phosphodiesterase | 10.1021/bi700561k | native |
| BcPMH | 3.00E+04 | diphenyl phosphate | *Burkholderia caryophilli* PG2952 | 10.1073/pnas.0903951107 | native |
| water ($k_w$) | 4.70E-12 | me-pNPP | | 10.1021/ja051603j | |

**Table S10A.** Crystallographic parameters, data collection and refinement statistics.

| | **desDEH** | **desE2D** | **desB** |
|---|---|---|---|
| **Crystallographic parameters** | | | |
| Space group | $P4_{1}2_{1}2$ | $P2_{1}2_{1}2_{1}$ | $P2_{1}2_{1}2_{1}$ |
| Unit-cell dimensions | 97.53, 97.53, 73.45Å | 37.88, 56.40, 92.92Å | 37.88, 56.40, 92.92Å |
| | 90, 90, 90˚ | 90˚, 90˚, 90˚ | 90˚, 90˚, 90˚ |
| **Data collection statistics** | | | |
| Resolution limits (outer shell) (Å) | 37.5-2.09(2.17-2.09) | 35.9-1.55 (1.62-1.55) | 35.9-1.55 (1.62-1.55) |
| No: of observed reflections (outer | 271206 (12852) | 207863 (11984) | 207863 (11984) |
| shell) | 15907 (795) | 26391 (1320) | 26391 (1320) |
| No: of unique reflections (outer shell) | | | |
| | 94.5 (96.4) | 94.8 (65.5) | 94.8 (65.5) |
| Completeness - ellipsoidal (outer | 74.2 (37.4) | 88.7 (34.7) | 88.7 (34.7) |
| shell) | 99.7 (60.4) | 99.6 (39.3) | 99.6 (39.3) |
| Completeness - spherical (outer shell) | | | |
| CC1/2 (outer shell) | | | |
| | 22.9 (192.8) | 10.0 (438.5) | 10.0 (438.5) |
| $R_{sym}$ (outer shell) (%)* | | | |
| Mean I/σ(I) (outer shell) | 10.0 (2.0) | 9.6 (1.6) | 9.6 (1.6) |
| **Refinement statistics** | | | |
| Resolution limits (Å) | 37.50-2.09 | 35.7-1.55 | 35.7-1.55 |
| Number of reflections (%) | 15880 (74.1) | 26381 (88.7) | 26381 (88.7) |
| Reflections used for $R_{free}$ | 814 | 1316 | 1316 |
| $R_{factor}$ (%)† | 21.0 | 18.4 | 18.4 |
| $R_{free}$ (%) | 25.5 | 23.2 | 23.2 |
| Model contents (average B(Å$^2$)) | | | |
| Protein atoms | 1804 (36.7) | 1700 (41.8) | 1700 (41.8) |
| Ligand | 0 | 0 | 0 |
| Ion/bugger | 13 (45.6) | 0 | 0 |
| Water molecules | 110 (37.1) | 101 (42.1) | 101 (42.1) |
| RMS deviations | 0.004 | 0.006 | 0.006 |
| Bond length (Å) | 0.699 | 0.80 | 0.80 |
| Bond angle (°) | | | |
| Ramachandran (favored (%)/outliers) | 99/0 | 97/1 | 97/1 |

* $R_{sym} = \Sigma|I_{avg}-I_i|/\Sigma I_i$

† R factor = $\Sigma|F_p - F_{pcalc}|/\Sigma F_p$, where $F_p$ and $F_{pcalc}$ are the observed and calculated structure factors; $R_{free}$ is calculated with 5% of the data.

**Table S10B.** Crystallographic parameters, data collection and refinement statistics.

| | **desHE2** |
|---|---|
| **Crystallographic parameters** | |
| Space group | $P2_12_12_1$ |
| Unit-cell dimensions | 38.11, 75.29, 88.18Å |
| | 90˚, 90˚, 90˚ |
| **Data collection statistics** | |
| Resolution limits (outer shell) (Å) | 38.1-1.42 (1.46-1.42) |
| No: of observed reflections (outer | 427253 (31804) |
| shell) | 47839 (3438) |
| No: of unique reflections (outer shell) | |
| | 98.2 (97.2) |
| Completeness (outer shell) | 99.8 (79.7) |
| CC1/2 (outer shell) | |
| | 5.5 (221.4) |
| $R_{sym}$ (outer shell) (%)* | |
| Mean I/σ(I) (outer shell) | 15.0 (1.3) |
| **Refinement statistics** | |
| Resolution limits (Å) | 38.1-1.42 |
| Number of reflections (%) | 47761 (98.0) |
| Reflections used for $R_{free}$ | 2388 |
| Rfactor (%)† | 18.8 |
| $R_{free}$ (%) | 23.5 |
| Model contents (average B(Å$^2$)) | |
| Protein atoms | 1944 (35.7) |
| Ligand | 0 |
| Ion/bufferer | 5 (45.2) |
| Water molecules | 208 (43.0) |
| RMS deviations | |
| Bond length (Å) | 0.004 |
| Bond angle (°) | 0.67 |
| Ramachandran (favored (%)/outliers) | 100/0 |

* $R_{sym} = \Sigma|I_{avg}-I_i|/\Sigma I_i$

† R factor = $\Sigma|F_p - F_{pcalc}|/\Sigma F_p$, where $F_p$ and $F_{pcalc}$ are the observed and calculated structure factors; $R_{free}$ is calculated with 5% of the data.
